# Phenotyping of *Klf14* mouse white adipose tissue enabled by whole slide segmentation with deep neural networks

**DOI:** 10.1101/2021.06.03.444997

**Authors:** Ramón Casero, Henrik Westerberg, Neil R Horner, Marianne Yon, Alan Aberdeen, Vicente Grau, Roger D Cox, Jens Rittscher, Ann-Marie Mallon

**Affiliations:** Mammalian Genetics Unit, MRC Harwell Institute, Harwell Campus, Didcot, OX11 0RD, UK; Institute of Biomedical Engineering, Department of Engineering Science, University of Oxford, Oxford OX3 7DQ, UK; Ground Truth Labs Ltd, 9400 Garsington Road, Oxford Business Park, Oxford, OX4 2HN, UK

**Keywords:** White adipose tissue, Deep learning, Segmentation, *Klf14*, Mouse, Phenotyping

## Abstract

White adipose tissue (WAT) plays a central role in metabolism, with multiple diseases and genetic mutations causing its remodeling. Quantitative analysis of white adipocyte size is of great interest to understand physiology and disease, but previous studies of H&E histology have been limited to a subsample of whole depot cross-sections. In this paper, we present the deep learning pipeline DeepCytometer, that can segment mouse and human whole slides (≃40,000 cells per mouse slide on average) using an adaptive tiling method, correct for cell overlap and reject non-white adipocytes from the segmentation. Using quantile colour maps we show intra- and inter-depot cell size heterogeneity with local correlation; quantile estimates also suggest significant differences in population estimates from 75 whole slides compared to smaller data sets. We propose three linked levels (body weight BW, depot weight DW and cell area quartiles) for exploratory analysis of mouse *Klf14* phenotypes in gonadal and subcutaneous depots. We find a rich set of phenotypes when stratifying by sex, depot and three genotype strata: (1) WTs/Hets with a Het father (Controls), (2) WTs with a Het mother, and (3) Hets with a Het mother (functional KOs or FKOs). Namely, at BW level, mean difference testing suggests that female FKOs are similar to Controls, but WTs with a Het mother are significantly larger. At DW and cell levels, linear models with interaction terms and BW or DW covariates, respectively, reveal phenotypes not shown by difference of means tests. For example, at DW level, gonadal and subcutaneous models are similar, and female FKOs have lower fat percentage than Controls due to both an offset and the DW/BW slope in the linear model. Meanwhile, female WTs with a Het mother have on average similar fat percentage to Controls, but as their slopes are close to zero, their DWs are uncorrelated to BW, suggesting that larger female WTs with a Het mother have lower fat percentage than smaller ones. In contrast to depot level, at cell level female gonadal phenotypes diverge from subcutaneous ones. Furthermore, male Controls and FKOs have similar average area values in subcutaneous depots, but area~DW slope flattening in FKOs suggests that larger DWs could be caused by cell size increase in Controls and by cell count increase in FKOs. Thus, DeepCytometer and associated exploratory analysis reveal new insights into adipocyte heterogeneity and phenotyping.

## 1. Introduction

White adipose tissue (WAT) provides the body’s main long-term energy storage and plays a central role in metabolism. It is distributed in subcutaneous (SAT) and visceral (VAT) depots, with SAT depots located abdominally, gluteofemorally, intramuscularly and in the upper body above and below the fascia, whereas VAT depots are omental, mesenteric, perirenal, retroperitoneal, gonadal and pericardial (Bjørndal et al., 2011; Parlee et al., 2014). The main components of WAT are unilocular lipid-filled white adipocytes or fat cells, comprising 90% of WAT mass, but less than 20% to 25% of cell population count (Eto et al., 2009; Vernon and Flint, 2003). WAT also contains blood vessels, adipocyte precursor cells, lymph nodes and nerves of the sympathetic nervous system and immune cells (Lenz et al., 2020), which interact with white adipocytes in adipose expansion and in metabolic disease (Nishimura et al., 2007; Rajbhandari et al., 2019; Skelly et al., 2019; Ye, 2011).

Healthy white adipocytes may present inter-depot heterogeneity in size, with SAT cells generally larger than VAT (Boumelhem et al., 2017; Fang et al., 2015; Glastonbury et al., 2020); adipocyte diameter is associated with BMI in both depots, but with obesity traits adjusted for BMI only in VAT (Honecker et al., 2021). There is also a sex effect, as females have significantly smaller adipocytes than males in VAT (Glastonbury et al., 2020). Females have larger adipocytes than males in SAT (Glastonbury et al., 2020; Small et al., 2018), but not significantly after adjusting for BMI, age and ancestry (Glastonbury et al., 2020). Obesity, or excessive WAT expansion occurs through two mechanisms: initial increase in cell size (hypertrophy) followed by increase in cell numbers (hyperplasia), the latter having poorer prognosis (Hausman et al., 2001). Obesity has reached pandemic levels and is strongly associated with a higher risk of type 2 diabetes, cardiovascular disease and cancer, as well as other chronic diseases such as fatty liver disease, hyperlipidaemia, hypertension, gout, restrictive lung disease, stroke, dementia, gallbladder disease, degenerative arthritis, and infertility (Blüher, 2019; Hausman et al., 2001; Myint et al., 2014; Pulit et al., 2019; Vazquez et al., 2007; Verboven et al., 2018). Where the expansion occurs is also important, as VAT expansion (especially omental and mesenteric) is correlated with higher disease risk, whereas SAT expansion can have a protective metabolic effect (Kim et al., 2007; Kusminski et al., 2012; Tchkonia et al., 2013; Yang and Civelek, 2020). Further, relative anthropometric measures of fat distribution such as waist-hip-ratio adjusted for BMI (WHRadjBMI) capture body fat percentage, and cardiovascular disease and type 2 diabetes risk, and may be better than BMI measures (Dale Caroline E. et al., 2017; Pulit et al., 2019; Winkler et al., 2018). Adipocyte mean area correlates with obesity measured through BMI, more strongly in VAT than SAT (Glastonbury et al., 2020). The study of WAT remodeling is interesting beyond obesity, as WAT expansion or wasting can be caused by certain diseases such as hypogonadism, Cushing’s syndrome, HIV, parasitic infection, or cancer-induced cachexia (Parlee et al., 2014).

The laboratory mouse is the leading model organism for the study of human disease, due to 99% of its genes having human orthologs, a wide catalogue of inbred strains and mutant models, and ease of breeding (Stamatoyannopoulos et al., 2012). Depending on the strain, 16-week-old mice on regular diets vary from 16.4% to 43.1% fat percentage in males and from 16.7% to 43.5% in females (Reed et al., 2007). For the C57BL/6NTac strain that we use in this work, fat percentage varies from 29.0% in males to 23.2% in females on control diets, and from 43.3% in males on a 12 week high fat diet (HFD) to 31.6% in females on a 2 week HFD (Podrini et al., 2013). As in humans, mice present inter-depot heterogeneity, with visceral gonadal rather than SAT or visceral mesenteric expansion being associated with metabolic disorders (van Beek et al., 2015). Common mouse models to study obesity are numerous genetics models such as the ob/ob, db/db, POMC, MC4 knockout, ectopic agouti and AgRP, FABP4-Wnt10b, LXRβ-/-, Sfrp5-/- and Timp-/- models, surgical/chemical models and diet induced obesity (Lutz and Woods, 2012; Parlee et al., 2014). Other models to study WAT phenotypes are R6/2 and CAG140 for Huntington’s disease (Phan et al., 2009) and CRH for Cushing’s syndrome (Williams-Dautovich et al., 2017). In this paper, as an exemplar we focus on the C57BL/6NTac *Klf14* knockout model (*Klf14^tm1(KOMP)Vlcg^*) previously developed by (Small et al., 2018). The *KLF14* (Krüpple-Like family 14) transcription factor is associated with metabolic syndrome and regulates gene expression in adipose tissue. This single exon gene is imprinted and only expressed from the maternally inherited allele (Parker-Katiraee et al., 2007), and is expressed more highly in females than males (Yang and Civelek, 2020). Homozygous females with the *KLF14* risk allele had significantly larger SAT white adipocytes in the Oxford Biobank (BMI-matched) (Small et al., 2018) and GTEx (not BMI-matched) (Glastonbury et al., 2020) data sets, compared to non-risk homozygous females. The ENDOX data set showed the same trend, but no statistical significance possibly due to the smaller data sample (Glastonbury et al., 2020). By contrast, VAT did not show significant adipocyte size differences in the GTEx (Glastonbury et al., 2020) data set.

Therefore, for human or animal model analysis, unbiased, high-throughput, global WAT quantitative analysis is of great interest. Adipocytes can be collagenase-isolated, counted and measured with a series of methods: microscope (Di Girolamo et al., 1971; Tchoukalova et al., 2003), hemocytometer (Bradshaw et al., 2003), Coulter counter (Hirsch and Gallian, 1968) and flow cytometry (Majka et al., 2014). However, those approaches consume the tissue and discard its spatial information, whereas whole slide imaging using fluorescent or conventional brightfield microscopy preserves tissue architecture within their slices. Given a sufficiently large field of view, ideally surveying the entire depot area, 2D assessment provides an excellent approximation of cell morphology. Hence our work focuses on the analysis of digitised Hematoxylin and Eosin (H&E) stained tissue sections.

In the first part of this work, we tackle adipocyte segmentation. Manual segmentation of adipocytes is highly accurate (Nishimura et al., 2007), but also labour intensive and slow. Larger scale studies require semi- or automatic instance segmentation methods, that need to be robust against preanalytical variation in histopathology (e.g. staining variability, tissue deformations, tears) and imaging artefacts (e.g. out of focus regions, bubbles). Traditional approaches were either based on hand-tailored features combined in *ad hoc* pipelines, including colour conversion, median filters, mathematical morphological operators, thresholding and watershed algorithms (Chen and Farese, 2002; Galarraga et al., 2012; Maguire et al., 2020; Osman et al., 2013; Zampirolli et al., 2010; Zhi et al., 2018); or based on training a pixel classifier with feature vectors extracted by a set of predetermined general-use filters (Arganda-Carreras et al., 2017).

Advances in deep learning have revolutionised biomedical image analysis, for instance cell detection and segmentation (Falk et al., 2019; Moen et al., 2019). In addition, these methods provide new ways for phenotyping specific cell types (Sirinukunwattana et al., 2020). DeepCell (Bannon et al., 2019; Van Valen et al., 2016) replaces predetermined filters (Arganda-Carreras et al., 2017) by deep convolutional neural networks (DCNNs) that learn optimal feature extraction for the target cell and microscopy modality, although its fully connected layers restrict input images to a pre-arranged size. To overcome this, Adipocyte U-Net (Glastonbury et al., 2020) uses a Fully Convolutional Network (FCN) (Long et al., 2015), so that images of variable size can be processed, up to the GPU memory limit. DCNNs successfully tackle stain variability and other colour variations using their generalisation capabilities, transfer learning, data augmentation (geometric transformations and colour) or a combination thereof. Even so, a DCNN-based whole H&E slide WAT segmentation pipeline needs to address several design decisions that we briefly review next.

Segmentation by pixel classification (typically as background, cell boundary or cell interior) is widely used (Glastonbury et al., 2020; Sirinukunwattana et al., 2020; Van Valen et al., 2016), but the results may need to be regularised, for example with an active contour (Van Valen et al., 2016). In addition, segmentation results tend to be worse where membranes touch, have less definition or are damaged. Segmentation results have been shown to improve by replacing pixel-classification by regression of the Euclidean distance transform (EDT) (Casero et al., 2019; Wang et al., 2019), i.e. distance of each pixel to the closest membrane point (see Appendix B.1 for insight in pixel-classification vs. EDT). Then, watershed seeds can be computed with peak detection (Casero et al., 2019) or with a DCNN trained as an object detector on the EDT (Wang et al., 2019), although neither can differentiate between adipocytes or other objects in the EDT. In this paper, we propose an EDT DCNN followed by a contour detector DCNN and watershed for full segmentation of H&E images.

The aforementioned segmentation approaches do not tackle white adipocyte overlap (see Fig. 1). Cell overlap has been tackled with a number of options such as a Physics model of light attenuation through the cytoplasm (Zhang et al., 2016) to ISOODL (Böhm et al., 2018), which rotates the plane of each cell in 3D space so that they do not overlap, but those increase the problem complexity. A general solution is using a single Shot Multibox Detector followed by individual cell segmentation with a U-Net (Sirinukunwattana et al., 2020), but this approach does not suit our whole-tile segmentation. Instead, we propose a DCNN based on QualityNet1 (Huang, Wu, and Meng 2016) that corrects each segmented object to account for cell overlap.

**Fig. 1.**
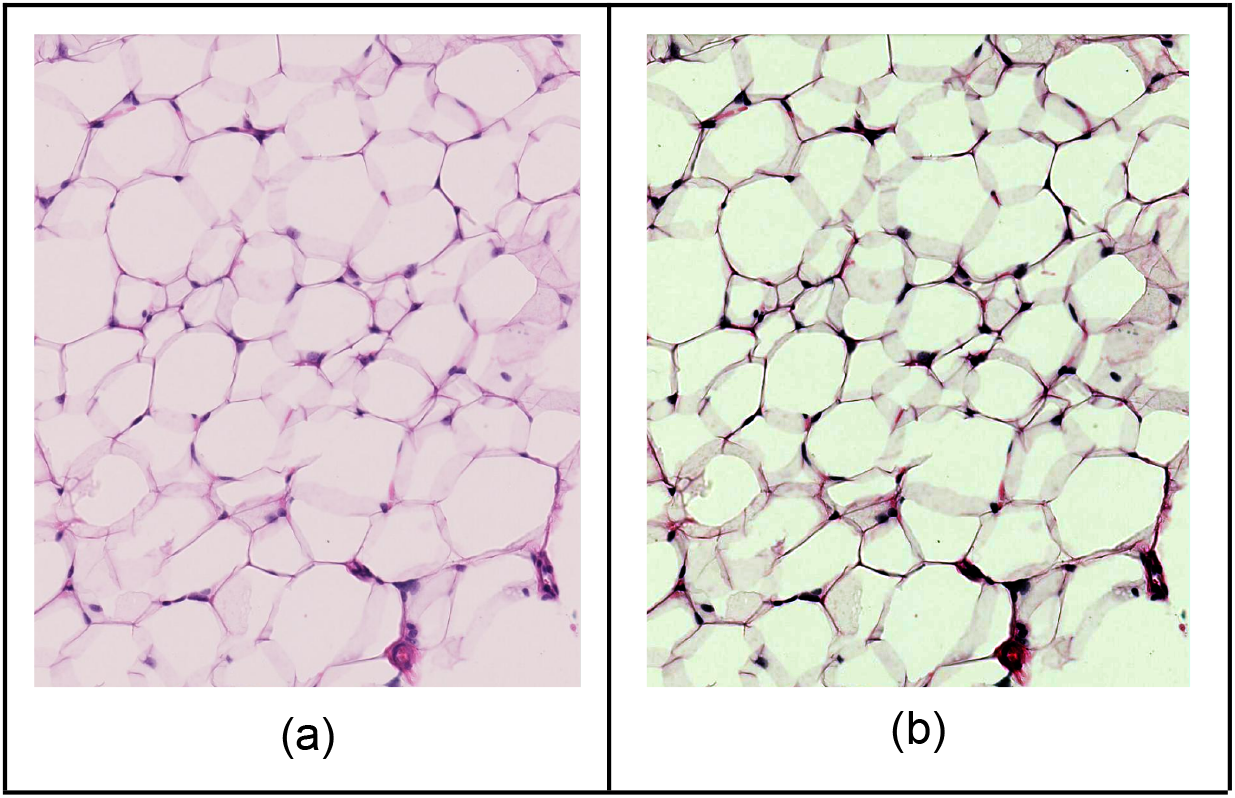
H&E histology section of white adipose tissue (WAT) displaying clear cytoplasm overlap between adipocytes. Cut thickness is 8 μm, but we have observed similar overlaps at 4 μm, 6 μm and 10 μm. These overlaps are effectively the 2D projection of adjacent cells slightly mounting each other in 3D. (a) Original microscope image. (b) Enhanced contrast by automatic levels rescaling and manual curves adjustment for better visualisation of overlaps.

Efficient computation requires pre-segment tissue areas to avoid segmenting large areas of empty background space (Kleczek et al., 2020; Muñoz-Aguirre et al., 2020); we tackle this problem with traditional image processing techniques. In addition, tissue areas are typically too large at full resolution to process in random-access or GPU memory, and need to be tiled and the results stitched together. Uniform titling is commonly used (Coudray et al., 2018; Glastonbury et al., 2020; Muñoz-Aguirre et al., 2020) as it is simple to implement, but the tile overlap needed to avoid dropping cells on the tile seams produces redundant computations; in this work, we propose an adaptive tiling algorithm to reduce that burden. Furthermore, it is necessary to differentiate between the cells of interest (mature white adipocytes) and other WAT components and image artefacts. For this, tiles can be accepted or rejected as a whole by an InceptionV3 network (Glastonbury et al., 2020; Honecker et al., 2021). This, however, has poor granularity and favours areas away from tissue edges and where other components are less prevalent. For full granularity, one can first detect valid cells and then segment them (Sirinukunwattana et al., 2020), or as we do in this paper, first compute a whole-tile segmentation and then classify each object as a valid or invalid white adipocyte.

Building a training data set for training/validating DCNNs can be done by cropping small histology windows, labelling them as valid/invalid for processing, and/or manually hand tracing the cells they contain. Previous multiple-cells-per-window approaches (Glastonbury et al., 2020; Van Valen et al., 2016; Wang et al., 2019) need that all training pixels are labelled as either background / membrane / cell interior, because they all contribute to the network’s loss function. But in practice, windows often contain ambiguous pixels, due to damaged, overlapping or unclear membranes. Furthermore, windows with non-adipocytes are precluded, or those objects need to be labelled as background or a new class, something laborious if they present intricate boundaries. Alternatively, one-cell-per-window approaches (Sirinukunwattana et al., 2020) can be trained granularly, as each training image contains a single cell. However, this also introduces redundant computation, as each training image must allocate space around the cell to provide spatial context. In this paper we search for a compromise, with a multiple-cells-per-window approach that can leave pixels unlabelled.

We address the challenges above to propose a whole slide white adipocyte segmentation pipeline called DeepCytometer. The challenges include: 1) whole slide processing of all tissue, ignoring the background; 2) tiling overlap compromise between segmenting all cells and reducing redundant computations; 3) colour variability in the slide; 4) cell segmentation considering that white adipocytes present as mostly background surrounded by a thin membrane, and that membranes can touch, overlap, be damaged or have poor definition; 5) differentiate between adipocytes and other WAT components, as well as image artefacts; 6) choice of training scheme, e.g. one-window multiple-cells vs. one-window one-cell, full vs. partial segmentation. We validate the segmentation results with summary statistics (Dice coefficient and relative area error) as in previous literature, but also propose to examine segmentation errors as a function of cell size, to assess whether all subpopulations are equally well segmented. We integrate DeepCytometer with the open source web-based application AIDA (Aberdeen and Malacrino, 2021) to navigate whole slides with the segmentation results.

In the second part of this paper, we phenotype *Klf14^tm1(KOMP)Vlcg^* C57BL/6NTac mice WAT, analysing DeepCytometer segmentations of 147 whole slides of H&E histology. We extend previous approaches that quantify median cell area from BMI-matched subjects (Small et al., 2018) or mean area (Glastonbury et al., 2020), and propose a phenotype framework with three interconnected linear model levels (body weight, depot weight and quartile cell area). We assess whether the linear model framework reveals more phenotypes than traditional approaches such as mean differences tests. Finally, we provide heat maps of cell area in whole slides, for qualitative assessment of spatial heterogeneity of subpopulations.

We provide all the code for the pipeline and experiments (Casero, 2021), as well as the pipeline supplementary files, histology images, hand-traced and pipeline segmentations (Casero et al., 2021a, 2021b).

## 2. Data and Methods

### 2.1. Mouse description, tissue acquisition and imaging

To develop and evaluate our methods we used *Klf14^tm1(KOMP)Vlcg^* C57BL/6NTac (B6NTac) mice tissue samples and additional data generated as part of the (Small et al., 2018) study. It should be noted that the single exon *Klf14* gene is imprinted and only expressed from the maternally inherited allele (Parker-Katiraee et al., 2007). This was taken into account by (Small et al., 2018) by crossing a Het parent with a WT parent, so that each offspring inherited a WT allele from the WT parent, and the *Klf14* gene knockout or a WT allele from the other parent (from the father, PAT, or the mother, MAT). We also take *Klf14* imprinting into account by using as controls the PAT mice and comparing them to the MAT WT and MAT Het (or functional KO, FKO) mice.

We used a total of 76 *Klf14*-B6NTac mice (n_female_=n_male_=38), of which 20 mice from the Control and FKO groups were used for training and testing the DeepCytometer pipeline, as well as the hand traced population experiment (summary in Table 1). Body and depot weight were measured with Sartorius BAL7000 scales. The mean ± standard deviation mouse body weights of those 20 mice were 26.6 ± 3.7 g (female PAT), 26.2 ± 2.9 g (female MAT), 37.6 ± 1.9 g (male PAT), 39.9 ± 3.8 g (male MAT).

**Table 1.** Description of mouse cohort used for CNN training (black), and hand traced cell population studies (blue). All slides acquired from subcutaneous tissue. Genot.: Parent and genotype (MAT/PAT: maternally/paternally inherited allele. WT: Wild type. Het: Heterozygous). BW: Body weight. Oth.: Other. Back.: Background. Cells/Oth./Back.: Number of hand traced white adipocytes/other tissue/background objects. Total win.: Number of 1,001×1,001 pixel windows extracted from the full slice at maximum resolution for training and testing. Win. that contain cells: Number of those windows that contain hand traced white adipocytes.

| ID | Sex | Genot. | BW (g) | Fold | Cells |  | Oth. | Back. | Total win. |  | Win. that contain cells |  |
| --- | --- | --- | --- | --- | --- | --- | --- | --- | --- | --- | --- | --- |
| 16.2d | m | MAT Het | 46.19 | 3 | 55 | 89 | 9 | 5 | 7 | 8 | 4 | 5 |
| 17.1c | f | MAT Het | 22.07 | 2 | 165 | 86 | 15 | 3 | 5 | 5 | 1 | 1 |
| 17.2c | f | MAT Het | 26.39 | 1 | 34 | 31 | 7 | 0 | 4 | 4 | 1 | 1 |
| 17.2f | m | MAT Het | 40.87 | 9 | 150 | 135 | 14 | 0 | 7 | 7 | 3 | 3 |
| 18.1a | f | MAT Het | 30.65 | 6 | 63 | 48 | 13 | 1 | 6 | 6 | 1 | 1 |
| 18.1e | m | MAT Het | 40.02 | 8 | 25 | 128 | 9 | 0 | 7 | 11 | 2 | 6 |
| 18.2b | f | MAT Het | 24.28 | 0 | 49 | 44 | 12 | 0 | 5 | 5 | 2 | 2 |
| 18.2d | f | MAT Het | 27.72 | 0 | 190 | 148 | 12 | 0 | 5 | 5 | 2 | 2 |
| 18.2g | m | MAT Het | 41.98 | 9 | 0 | 0 | 9 | 0 | 3 | 3 | 0 | 0 |
| 18.3b | m | MAT Het | 34.52 | 8 | 83 | 73 | 11 | 3 | 8 | 8 | 4 | 4 |
| 18.3d | m | MAT Het | 36.08 | 2 | 199 | 179 | 16 | 1 | 8 | 8 | 5 | 5 |
| 36.1a | f | PAT Het | 31.42 | 5 | 96 | 82 | 5 | 0 | 5 | 5 | 2 | 2 |
| 36.1b | f | PAT WT | 29.25 | 5 | 65 | 56 | 8 | 0 | 6 | 6 | 2 | 2 |
| 36.1c | f | PAT Het | 27.18 | 7 | 187 | 159 | 8 | 1 | 5 | 5 | 2 | 2 |
| 36.1i | m | PAT Het | 36.55 | 1 | 251 | 218 | 11 | 2 | 11 | 11 | 7 | 7 |
| 36.3d | m | PAT Het | 40.77 | 6 | 111 | 100 | 21 | 1 | 8 | 8 | 5 | 5 |
| 37.1c | f | PAT WT | 23.69 | 3 | 157 | 122 | 13 | 2 | 5 | 5 | 2 | 2 |
| 37.1d | f | PAT WT | 21.20 | 4 | 17 | 13 | 15 | 0 | 5 | 5 | 2 | 2 |
| 37.2g | m | PAT Het | 36.98 | 4 | 147 | 126 | 13 | 0 | 8 | 8 | 5 | 5 |
| 37.4a | m | PAT Het | 36.11 | 7 | 73 | 66 | 11 | 5 | 8 | 8 | 3 | 3 |
| Total |  |  |  |  | 2,117 | 1,903 | 232 | 24 | 126 | 131 | 55 | 60 |

The histopathology screen involved fixing, processing and embedding in wax, sectioning and staining with Hematoxylin and Eosin (H&E) both inguinal subcutaneous and gonadal adipose depots. For paraffin-embedded sections, all samples were fixed in 10% neutral buffered formalin (Surgipath) for at least 48 hours at RT and processed using an Excelsior™ AS Tissue Processor (Thermo Scientific). Samples were embedded in molten paraffin wax and 8 μm sections were cut through the respective depots using a Finesse™ ME+ microtome (Thermo Scientific). Sampling was conducted at 2 sections per slide, 3 slides per depot block onto simultaneous charged slides, stained with haematoxylin Gill 3 and eosin (Thermo scientific) and scanned using an NDP NanoZoomer Digital pathology scanner (RS C10730 Series; Hamamatsu).

### 2.2. Ground truth hand traced data set for CNN training and cell population studies

We created two slightly different hand traced data sets. For DeepCytometer training and validation, we randomly sampled each of the 20 training histology slides to extract a total of 55 histology training windows with size 1001×1001 pixels and traced white adipocytes, other types of tissue (connective, vessels, muscle, brown adipocytes, etc.) and background areas. Another 71 windows were manually selected to add more examples of only other types of tissue to train the tissue classifier. This first data set is summarised in Table 1 in black.

For cell population studies, we added 5 random windows from 2 mice whose cell population was undersampled, and we removed those white adipocyte objects with ambiguous interpretation, namely fully overlapped, suffering from image artifacts or comprising very small gaps between clear adipocytes. This second data set is summarised in Table 1 in blue.

The total number of hand traced white adipocytes is 2,117 and 1,903, respectively, with a roughly balanced split between the four groups formed by the female/male and PAT/MAT partitions (Table 2). The total number of “other tissue” objects is 232, and of “background”, 24. (Note that cell objects tend to be much smaller than “other” or “background” objects). Each window contained one or more of these types of objects. Hand tracing was performed with the image editor Gimp (The GIMP Development Team, 2019) over a month. Automatic levels correction and manual curves adjustment was temporarily applied to each window to improve image contrast for the human operator. Contours were drawn as linear polygons on the outermost cell edge, accounting for cell overlap (Fig. 1), and exported with an *ad hoc* plugin as SVG files for further processing. For “other” or “background”, representative linear polygons were drawn, avoiding complex boundaries. This approach produced partially labelled training windows.

**Table 2.**
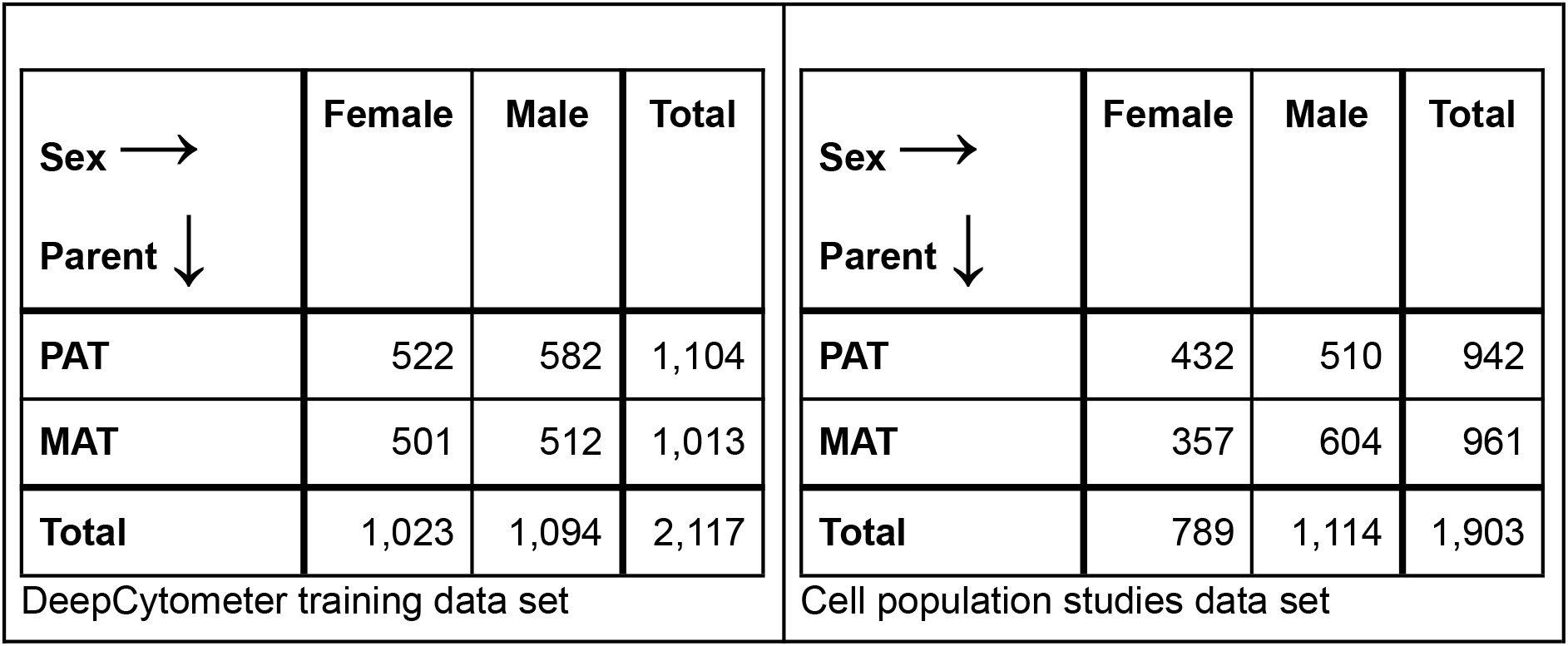
Number of hand traced white adipocytes stratified by sex and parent. These tables are summaries of Table 1.

### 2.3. Coarse tissue mask and adaptive tiling for full processing of whole slides

#### 2.3.1. Coarse tissue mask

Histology slides in Hamamatsu NDPI format can be read by blocks of arbitrary size at a fixed number of precomputed resolution levels with OpenSlide (Satyanarayanan et al., 2013). At the highest resolution level, our images have pixel size 0.454 μm. We read the whole histology slide at the precomputed ×16 downsampling level (7.26 μm pixel size), applied contrast enhancement and computed the mode *Mo_colour_* and standard deviation *σ_colour_* in each RGB channel of the image. We assume that the background colour is centered around *Mo_colour_*, as background pixels are more numerous than tissue pixels. To segment the tissue, we thresholded the downsampled image for pixels that are darker than *Mo_colour_* – *k_σ_σ_colour_* in all RGB channels (see Fig. 2b), where *k*_σ_ = 0.25. Then, we applied morphological closing with a 25×25 kernel at ×16 downsampling level, filled holes smaller than 8,000 pixels (421,759 μm^2^), and removed connected components smaller than 50,000 pixels (2,635,994 μm^2^).

**Fig. 2.**
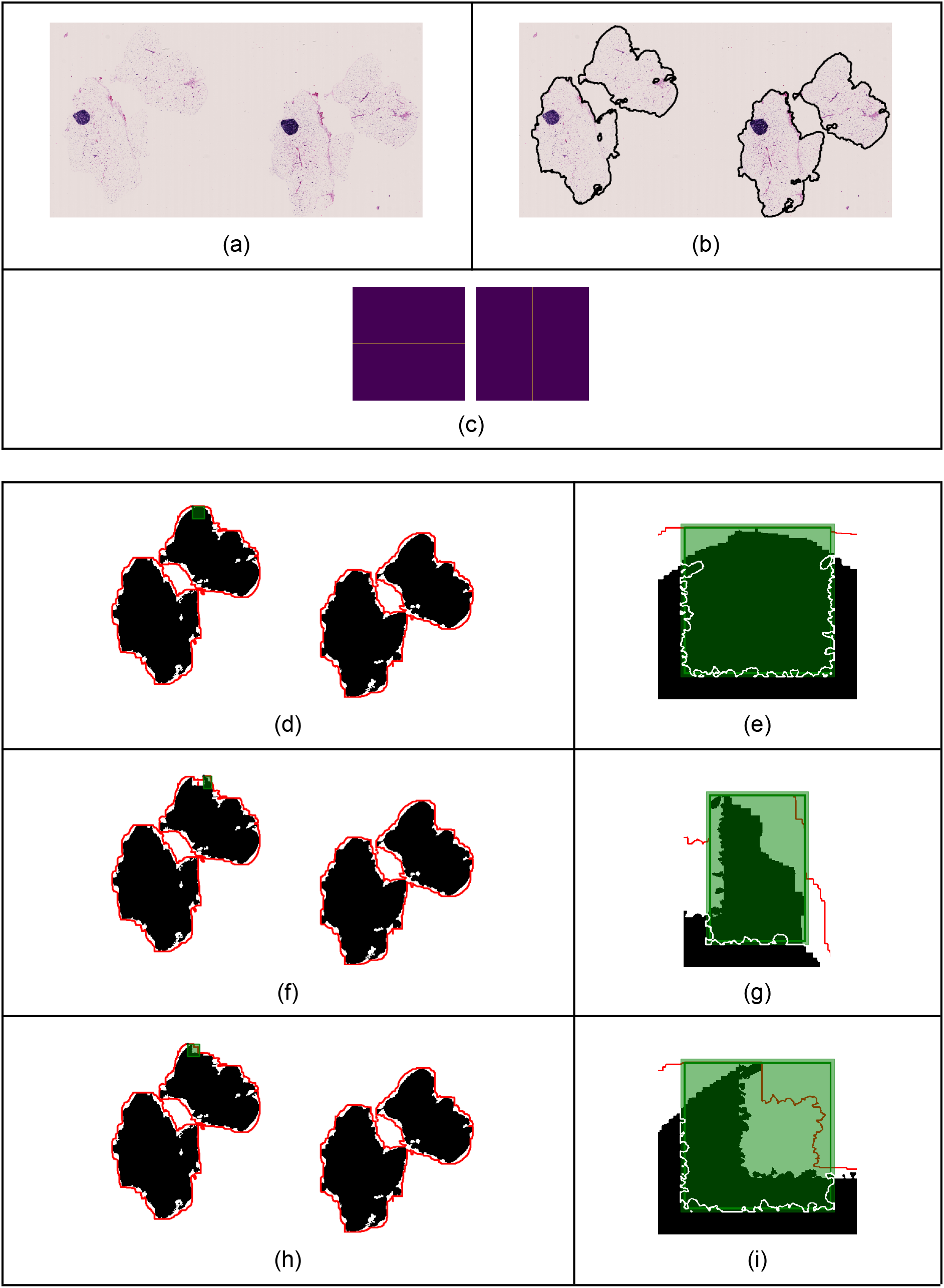
Coarse tissue segmentation and adaptive tiling. (a) Whole histology slide, female MAT. (b) Coarse tissue mask. (c) Horizontal and vertical line convolution kernels. (d)-(i) Black mask: Coarse tissue mask at three consecutive iterations. Red contour: Boundary of convolutions with horizontal and vertical line kernels. Green square/rectangle: Block chosen for histology segmentation at each iteration. Green solid line: Separates inner part of the block from added border to account for ERF. White contour: Edges that are not removed from the coarse mask. (d), (f), (h): Whole slide. (e), (g), (i): Detail around processing block.

#### 2.3.2. Adaptive tiling

When using uniform tiling, to guarantee that any cell will be processed whole in at least one tile, adjacent tiles need to overlap by *RF* + *D_max_*, where *RF* is the receptive field or diameter of input pixels that affect each output pixel, and *D_max_* is the diameter of the largest cell. This overlap introduces repeated processing of the same pixels and multiple segmentations of the same cells from adjacent tiles, but when it is ignored (Glastonbury et al., 2020), cells cropped by tile edges need to be discarded.

To ameliorate this problem, we propose an iterative tiling algorithm that adapts the block size and overlap of each new tile according to the local cell size and tissue mask, such that the whole coarse tissue mask is covered, all cells are segmented, and redundant computations are reduced.

For this, first we flag all pixels in the coarse tissue mask as “to be processed”. To find the location of the first block, the mask is convolved with two small linear kernels. Pixels > 0 in both outputs are potential locations for the block’s top-left corner. Any of those locations guarantee that the block has at least one mask pixel on the top and left borders. The algorithm chooses the first of the candidate pixels, in row-column order. The bottom-right corner of the block is initially chosen to obtain a block with maximum size 2751×2751 pixels (maximum allowed by our NVIDIA TITAN RTX GPU memory). The block’s right side and bottom are then cropped to remove empty columns or rows, producing an adaptive size. Finally, the block is extended half the effective receptive field on each side, to prevent border effects. (This extra border is discarded after image processing operations.) If the block overflows the image, it is cropped accordingly. The image block is then segmented; pixels from cells cropped by the edges keep their “to be processed” flag, so they will be included in another block. The rest of the mask pixels are cleared, and the process is repeated iteratively choosing new adaptive blocks until the whole tissue mask is cleared. Pseudocode and details for the algorithm are provided in Appendix B.3, and an example of its behaviour is shown in Fig. 2:

### 2.4. Deep CNN architectures

In this section we describe the function of the four different DCNNs used in this work, with illustrative examples (Fig. 3–6), a summary of their architectures (Table 4), and the calculation of their effective receptive fields (ERFs) (Table 9). Training and validation details are provided in Appendix B.4. These networks are components of the segmentation sub-pipeline that we describe in the next section. The networks are fully convolutional (Long et al., 2015), so that tile size can be adjusted to available GPU memory and the needs of the adaptive tiling. We used a stride of 1 to avoid downsampling followed by deconvolution, and thus preserve high-resolution segmentation following (Sherrah, 2016). We used atrous or dilated convolution (Chen et al., 2016, 2014; Yu and Koltun, 2015) to enlarge the ERF following (Sherrah, 2016; Van Valen et al., 2016). RGB 8-bit unsigned integer values in the image files were converted to 32-bit float type and scaled to [0, 1] or [-1, 1] depending on the specifications of the network they are being fed to.

**Fig. 3.**
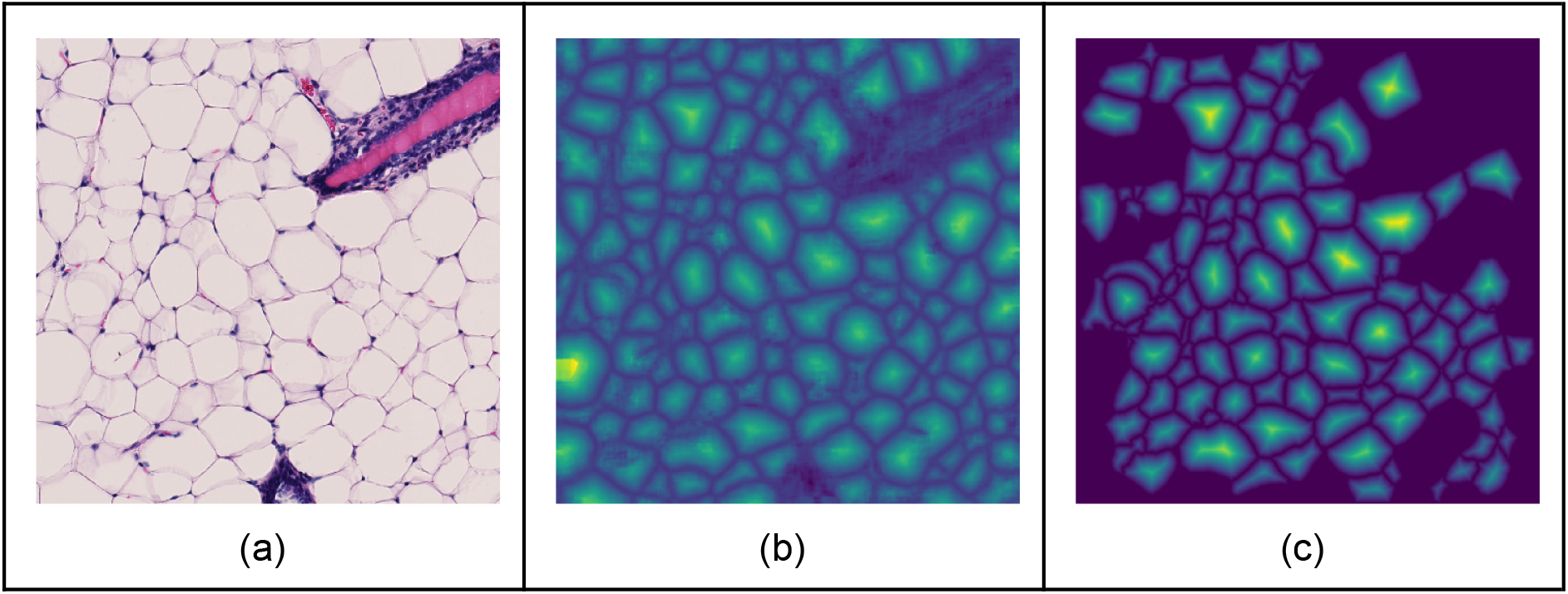
Illustration of EDT CNN. (a) Input histology image. (b) EDT computed by the network, trained without the test data. (c) Ground truth EDT computed from the contours in Fig. 4b.

#### 2.4.1. Histology to EDT regression CNN (EDT CNN)

This network (see architecture in Table 4a), based on previous work by (Casero et al., 2019; Cientanni and Casero, 2018) and similar to (Bai and Urtasun, 2017; Wang et al., 2019) —as discussed in Section 1—takes an input histology image and estimates the Euclidean Distance Transform (EDT) as the Euclidean distance of each pixel to the closest label boundary point (see Section B.4.3 for details). This produces an image similar to an elevation map (Fig. 3), where each “hill” defines an object (e.g. a white adipocyte, an area of muscle tissue or a vessel cross-section). If the object is a white adipocyte, troughs represent the cell’s membrane boundary or a compromise boundary between overlapping cells. Trough points are all critical points (extrema or saddle points) but have different values or “elevations”. We found that troughs in our whole slides could not be segmented with simple segmentation methods like in (Bai and Urtasun, 2017; Wang et al., 2019). Instead, we trained the following Contour network for that task.

#### 2.4.2. EDT to Contour detection CNN (Contour CNN)

This network (see architecture in Table 4b) classifies each EDT pixel as either belonging or not to a trough (Fig. 4).

**Fig. 4.**
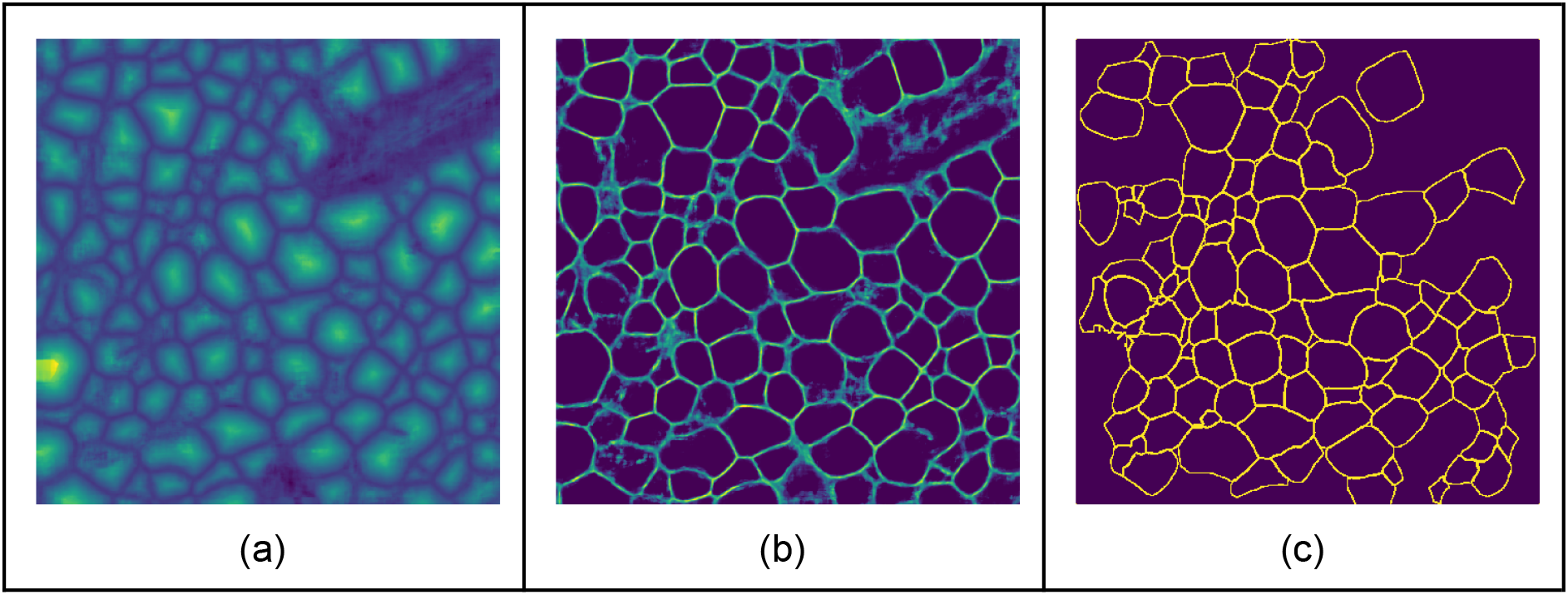
Illustration of Contour CNN. (a) The Input to the network is the output from Fig. 3c. (b) Contours computed by the network, trained without the test data. (c) Ground truth contours derived from hand segmentations with overlaps removed (the hand segmentation leaves unlabelled pixels).

#### 2.4.3. Pixel-wise tissue classifier CNN (Tissue CNN)

This network (see architecture in Table 4c) classifies each pixel from the histology block as “other type of tissue” vs. “white adipocyte” or “background” (see Fig. 5). “White adipocyte” and “background” are combined in one class because a gap in the tissue and the inside of a white adipocyte have the same appearance in the histology. The output of this network is used to classify a segmented object as white adipocyte or not, according to the proportion of “white adipocyte” / “background” pixels it contains (see Section 2.5 for details).

**Fig. 5.**
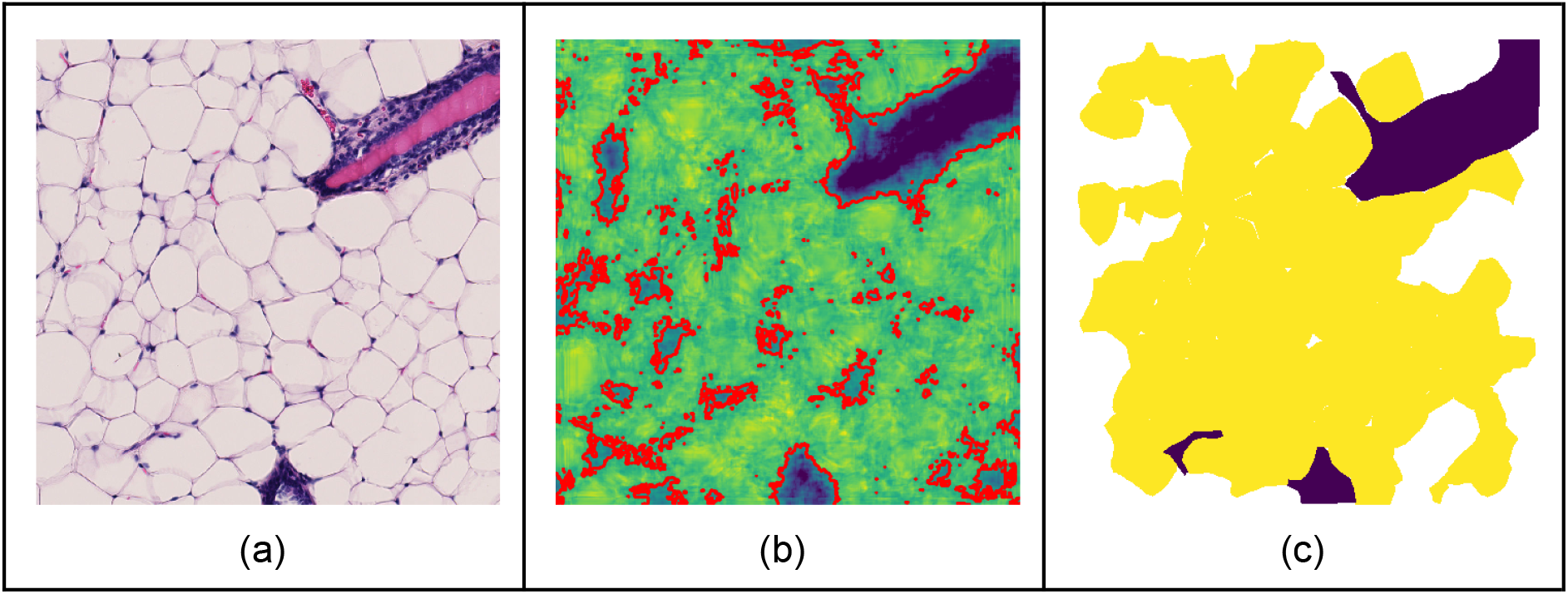
Illustration of Tissue CNN. (a) Input histology image. (b) Classification computed by the network from 0 (dark blue) to 1 (yellow). Red contours correspond to classification threshold 0.5. (c) Ground truth for classification. In white, unlabelled pixels.

#### 2.4.4. Segmentation correction regression CNN (Correction CNN)

This network (see architecture in Table 4d) takes the cropped and scaled histology of a single object, multiplied by a mask derived from its segmentation, and estimates which pixels underestimate (detection error) or overestimate (false positive) the segmentation. This output is then used to correct the object’s segmentation (see Section 2.5 for details). Because each object’s segmentation is corrected separately, corrected boundaries can overlap. To create an input for the network, the histology tile is cropped using a square box with at least twice the size as the segmentation mask’s bounding box, and scaled to a fixed size of 401×401 pixels to make the network blind to cell size. The scaling factor is s=401/L, where L×L is the size of the cropping window. Then, the histology RGB values are multiplied by +1 within the segmentation mask, and by −1 without. By contrast, (Huang et al., 2016)‘s QualityNet1 multiplied the histology RGB values by 0 without. Our approach preserves the information outside the segmentation, while still partitioning the histology image into two sets of inside/outside pixels. Moreover, instead of estimating one single quality measure for the whole input image as in QualityNet1, we estimate whether the segmentation is correct *per pixel*, computing a value between −1 (undetected pixel) to +1 (false positive pixel) through 0 (correctly segmented pixel).

**Table 3.**
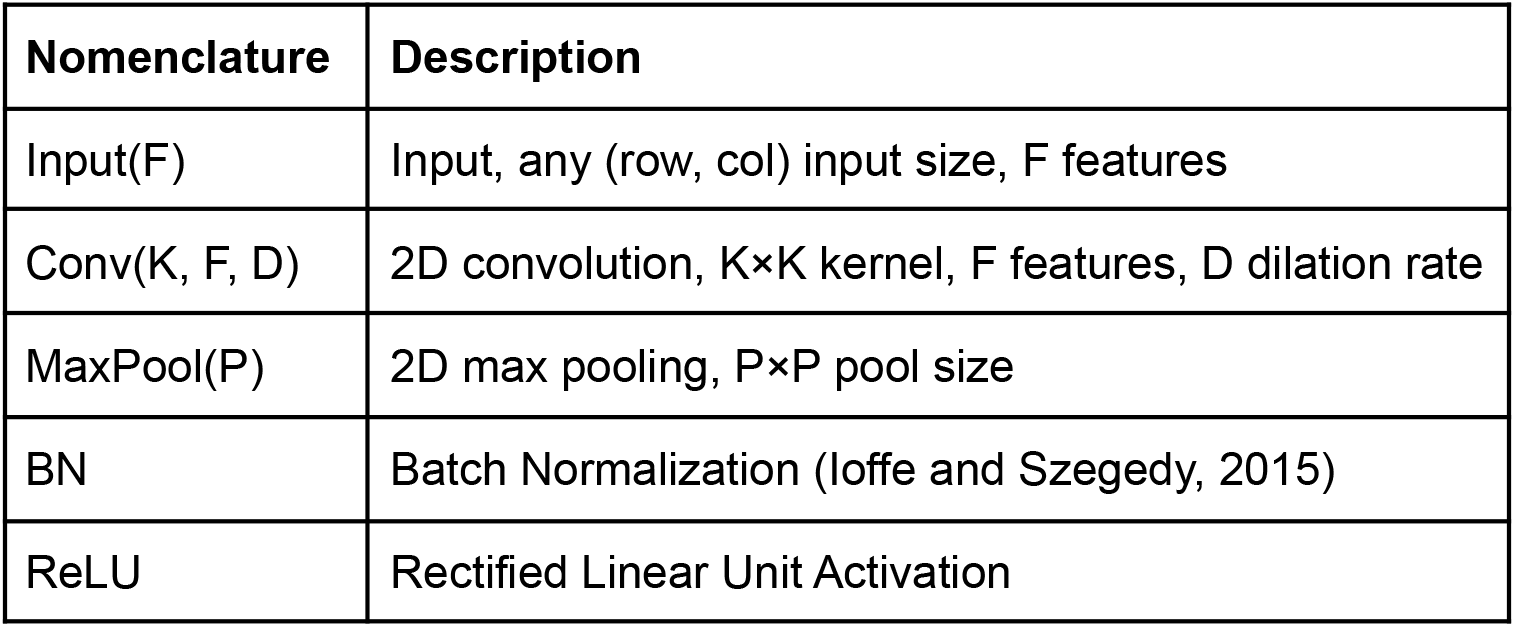
Nomenclature for layers of pipeline CNNs. All Conv and MaxPool layers have stride 1, and zero padding so that their output has the same (row, column) size as their input.

**Table 4.**
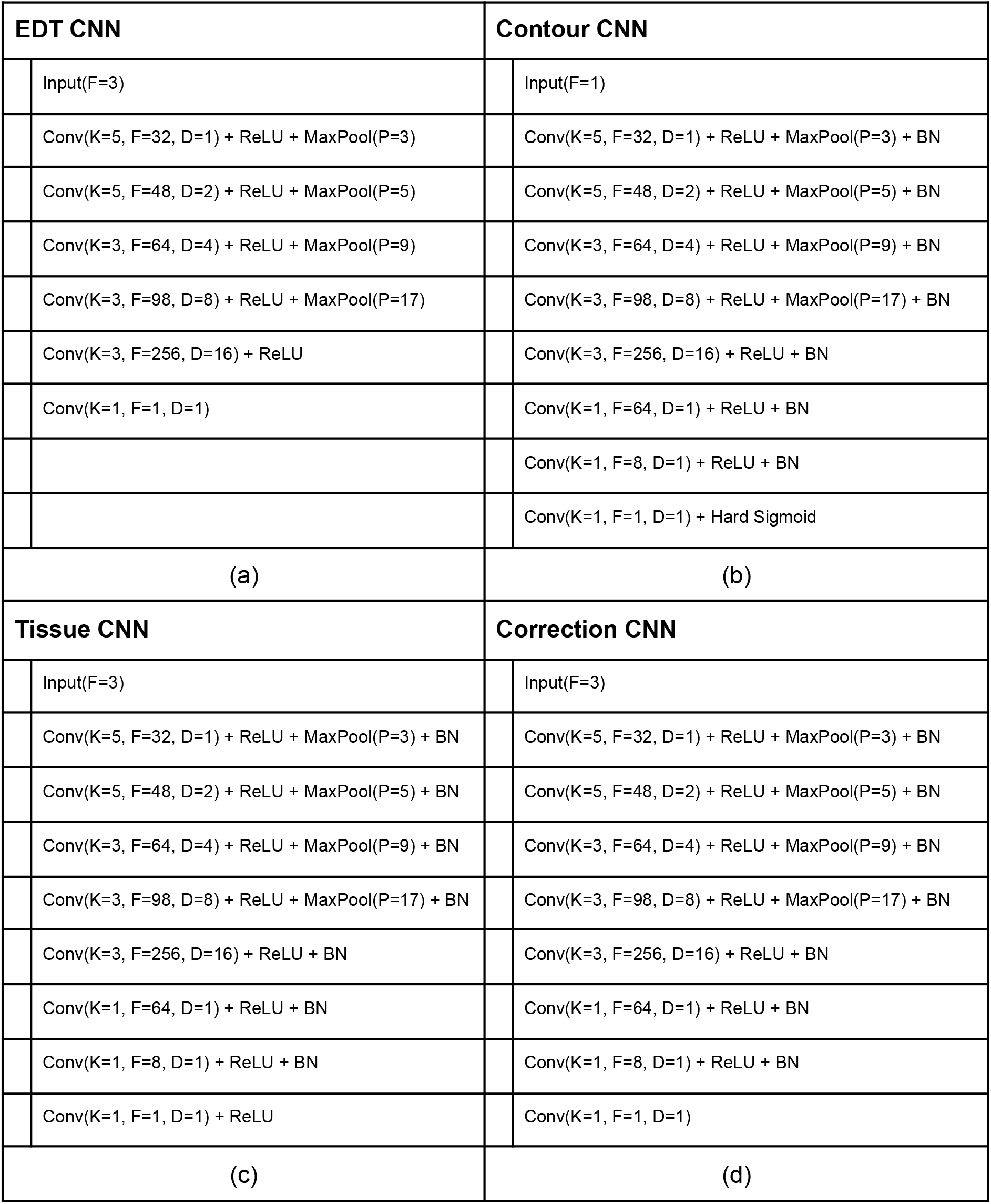
Description of the four CNN architectures used by the DeepCytometer pipeline. (See nomenclature in Table 3).

| EDT CNN |  | Contour CNN |  |
| --- | --- | --- | --- |
|  | Input(F=3) |  | Input(F=1) |
|  | Conv(K=5, F=32, D=1) + ReLU + MaxPool(P=3) |  | Conv(K=5, F=32, D=1) + ReLU + MaxPool(P=3) + BN |
|  | Conv(K=5, F=48, D=2) + ReLU + MaxPool(P=5) |  | Conv(K=5, F=48, D=2) + ReLU + MaxPool(P=5) + BN |
|  | Conv(K=3, F=64, D=4) + ReLU + MaxPool(P=9) |  | Conv(K=3, F=64, D=4) + ReLU + MaxPool(P=9) + BN |
|  | Conv(K=3, F=98, D=8) + ReLU + MaxPool(P=17) |  | Conv(K=3, F=98, D=8) + ReLU + MaxPool(P=17) + BN |
|  | Conv(K=3, F=256, D=16) + ReLU |  | Conv(K=3, F=256, D=16) + ReLU + BN |
|  | Conv(K=1, F=1, D=1) |  | Conv(K=1, F=64, D=1) + ReLU + BN |
|  |  |  | Conv(K=1, F=8, D=1) + ReLU + BN |
|  |  |  | Conv(K=1, F=1, D=1) + Hard Sigmoid |
| (a) |  | (b) |  |
| Tissue CNN |  | Correction CNN |  |
|  | Input(F=3) |  | Input(F=3) |
|  | Conv(K=5, F=32, D=1) + ReLU + MaxPool(P=3) + BN |  | Conv(K=5, F=32, D=1) + ReLU + MaxPool(P=3) + BN |
|  | Conv(K=5, F=48, D=2) + ReLU + MaxPool(P=5) + BN |  | Conv(K=5, F=48, D=2) + ReLU + MaxPool(P=5) + BN |
|  | Conv(K=3, F=64, D=4) + ReLU + MaxPool(P=9) + BN |  | Conv(K=3, F=64, D=4) + ReLU + MaxPool(P=9) + BN |
|  | Conv(K=3, F=98, D=8) + ReLU + MaxPool(P=17) + BN |  | Conv(K=3, F=98, D=8) + ReLU + MaxPool(P=17) + BN |
|  | Conv(K=3, F=256, D=16) + ReLU + BN |  | Conv(K=3, F=256, D=16) + ReLU + BN |
|  | Conv(K=1, F=64, D=1) + ReLU + BN |  | Conv(K=1, F=64, D=1) + ReLU + BN |
|  | Conv(K=1, F=8, D=1) + ReLU + BN |  | Conv(K=1, F=8, D=1) + ReLU + BN |
|  | Conv(K=1, F=1, D=1) + ReLU |  | Conv(K=1, F=1, D=1) |
| (c) |  | (d) |  |

#### 2.4.5. Effective receptive field (ERF)

The theoretical receptive field (span of input pixels that contribute to an output pixel) can be computed considering the properties of convolutions, downsampling and pooling. However, the weight of an input pixel’s contribution decreases quickly towards the edges of the span, and thus the effective receptive field (ERF) is much smaller than the theoretical one (Luo et al., 2016). To estimate the ERF, we used gradient backpropagation (Luo et al., 2016), but replaced ReLU activations by Linear activations and Max Pooling by Average Pooling to avoid numerical instabilities. The ERF was around 131 × 131 pixels (Table 9), or 37.1% of maximum cell diameter (160 μm or 353 pixels), causing the EDT CNN to clip the estimated distance to the membrane for distances larger than the ERF. However, the segmentation validation showed no performance drop for large cells. This is because distant points do not contribute essential information to the Contour CNN. This was convenient, as increasing the ERF is computationally expensive, generally requiring deeper networks.

### 2.5. Segmentation sub-pipeline

The segmentation sub-pipeline combines the EDT, Contour, Tissue and Correction CNNs with traditional image processing methods (Fig. 7b) to segment an input histology image tile, producing one label per white adipocyte.

#### 2.5.1. Histology colour correction

To estimate the typical background colour of the training data set, we computed density histograms with 51 bins between 0-255 for each RGB channel of the 126 training images (Table 1). The 50% HD quantile was computed for each bin, producing a median histogram for each channel. The modes in each median channel were taken as the typical background colour, R_target_ = 232.5, G_target_ = 217.5, B_target_ = 217.5. Colour correction was applied for inference but not for training, to reduce overfitting. To apply colour correction to a histology slide, we estimated the mode intensity of each channel R_mode_, G_mode_, B_mode_. Then, each colour channel was corrected as Î = I_R_ - R_mode_ + R_target_, Î = I_G_ - G_mode_ + G_target_, Î = I_B_ - B_mode_ + B_target_.

#### 2.5.2. White adipocyte label segmentation without overlap (Auto)

The colour-corrected histology image was used as input to the EDT CNN, and its output to the Contour CNN. To conservatively detect pixels inside objects, we thresholded the resulting image with a zero threshold (pixels on or near EDT troughs have values > 0). This produced one connected component inside each cell or object. We filled holes with fewer than 10,000 pixels, and each connected component was given a different label. Components with fewer than 400 pixels were removed. The colour-corrected histology image was also fed to the Tissue CNN, and pixels with score > 0.5 were labelled as white adipocyte pixels. All non white adipocyte pixels that do not already belong to a seed were labelled as a single new seed. Seeds were expanded using a watershed algorithm on the negative of the EDT surface (the negative sign turned hills into basins). Each watershed basin corresponded to a candidate object. Objects were rejected if any of the following were true: 1) were smaller than 1,500 pixel (308.9 μm^2^), 2) did not overlap at least 80% with the coarse tissue mask, 3) did not contain at least 50% white adipocyte pixels, 4) touched an edge, 5) were larger than 200,000 pixels (41,187.4 μm^2^) - the largest training cell was 92,544 pixel (19,058.2 μm^2^). 6) *compacteness*^−1^ = *perimeter*^2^ /4π *area* > 2 Objects that were inside another object were merged into the surrounding object. Each surviving label was considered to segment one white adipocyte, but without overlap.

#### 2.5.3. Segmentation label correction with overlap (Corrected)

The colour-corrected histology image was cropped and resized around each valid white adipocyte label, and passed through the Correction CNN as described above. Output pixels with scores > 0.5 were added to the label, and pixels with scores ≤ −0.5 were removed. Label holes were filled, and the connected component that had the best overlap with the input label was kept as the corrected label. Finally, the corrected label was smoothed using a closing operator with an 11×11 pixel square kernel. (See Fig. 6c).

**Fig. 6.**
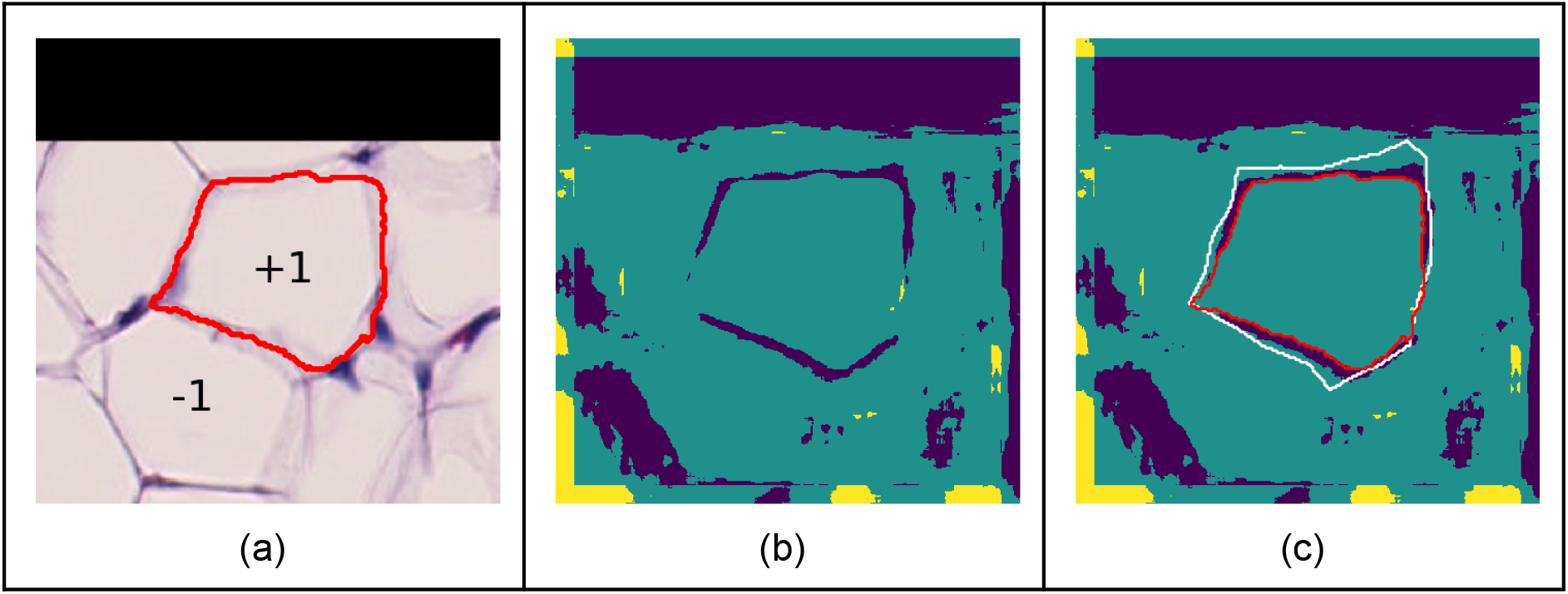
Illustration of Correction CNN. (a) Cropped and scaled histology region around Auto segmentation (red contour). Size 401×401 pixel. Intensity values multiplied by +1 inside segmentation, and by −1 outside segmentation. (b) Network estimation of whether pixels are correctly segmented: Oversegmented (yellow), correct (green), undersegmented (blue). (c) Correction overlayed with manual ground truth (white contour) and input segmentation (red contour).

**Fig. 7.**
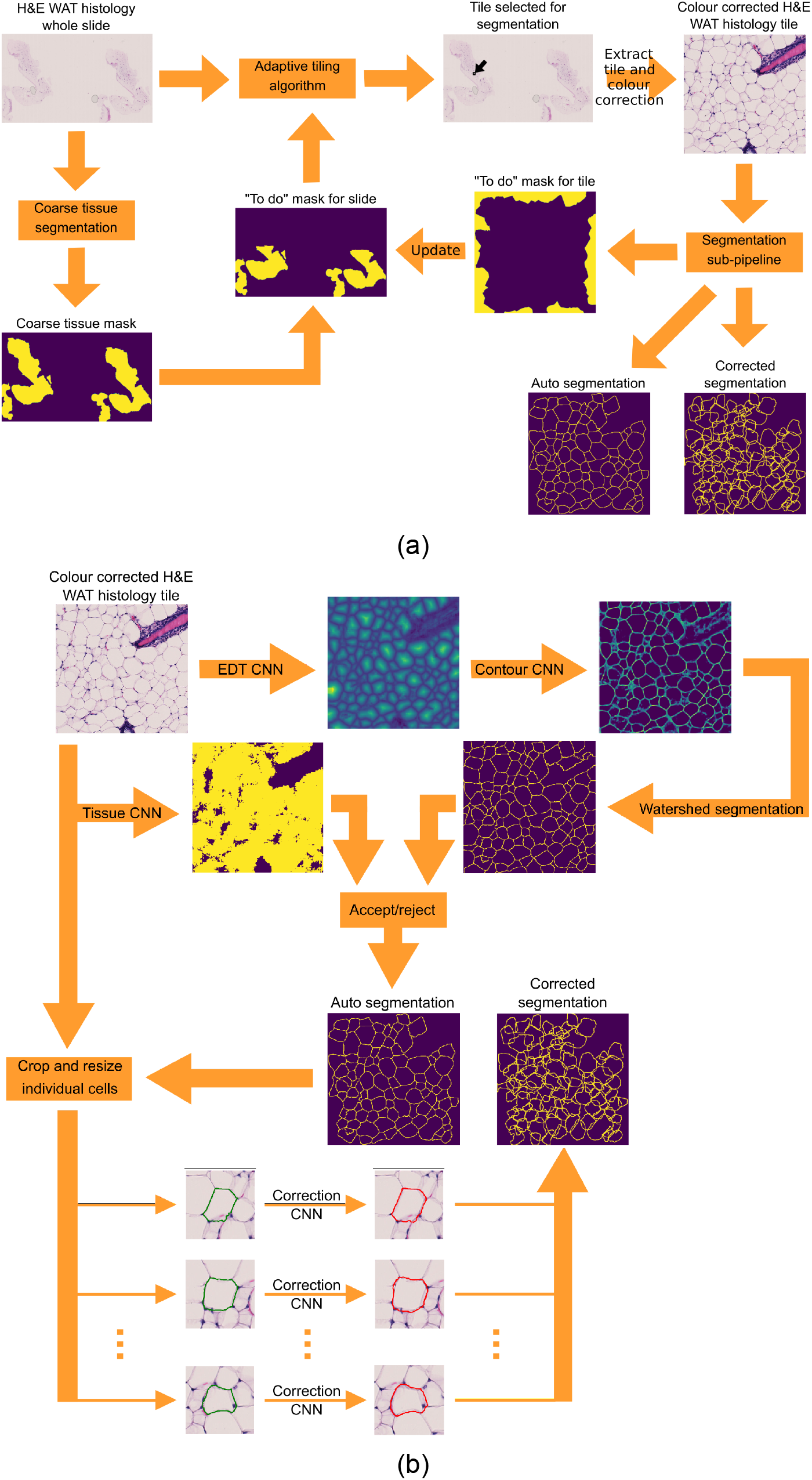
Pipeline diagrams. (a) DeepCytometer pipeline. (b) White adipocyte segmentation sub-pipeline.

#### 2.5.4. Segmentation output to AIDA user interface

Each corrected label was converted to a linear polygon with vertices *c* = {(*x*_0_, *y*_0_),…, (*x*_*P*−1_, *y*_*P*−1_)} using Marching Squares (Lorensen and Cline, 1987). Point coordinates were converted from the processing window to the whole histology image using 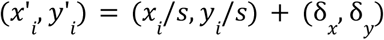, where *s* is the scaling factor and (*δ_x_, δ_y_*) are the coordinates of the processing window’s top-left pixel within the whole histology image. The contours were then written to a JSON file that can be read by the browser interface AIDA (Fig. 15).

#### 2.5.5. Tissue CNN validation

The Tissue CNN was applied to each of the 126 training images (see Table 1) according to the 10-fold split, and classification scores > 0.5 were labelled as white adipocyte pixels. Then, for each hand traced contour we computed the white adipocyte score *Z_obj_* = #*WA*/(#*WA* + #*NWA*) ∈ [0,1], where #*WA* stands for the number of white adipocyte pixels within the object, and #*NWA* for the number of non white adipocyte pixels. This *Z_obj_* was compared to the ground truth score *Z_gt_* = {0,1} to compute the receiver operating characteristic (ROC) curve, weighting each object by its number of pixels. This way, the ROC takes into account the fact that white adipocyte objects tend to be much smaller and more numerous than other objects. Namely, this allows us to interpret the classification error in terms of the more balanced tissue area classification error (our training contours contain ≈ 23.7 · 10^6^ white adipocyte pixels and ≈ 45.1 · 10^6^ non white adipocyte pixels).

#### 2.5.6. Segmentation sub-pipeline validation

We applied the segmentation sub-pipeline to each of the 55 colour-corrected images with hand traced cells (see Table 1), using 10-fold cross validation. This produced Auto and Corrected contours for each image that we compared to the hand traced ones for validation.

The literature relies on summary statistics for segmentation validation. For instance, per-image average diameter (Galarraga et al., 2012), mean Dice coefficient (DC) and mean Jaccard Index (Van Valen et al., 2016), median cell area/volume per subject (Small et al., 2018), IoU and F1-score (Caicedo et al., 2019) and mean cell area (Glastonbury et al., 2020). As summary statistics, we used the DC and relative area error between each pipeline-produced contour and the hand traced (ht) contours in the image. The *DC* = 2*a*_*ht ∩ pipeline*_/(*a_ht_* + *a_pipeline_*), where *a* are areas computed with polygon operations.

The highest DC was considered the best match. DC ≤ 0.5 were considered no match, as they are usually contours that segment an adjacent object. To compare cell populations, cell area distributions were computed using box-and-whisker plots with median notches for the hand traced, Auto and Corrected contours with a match (Fig. 8). The lower whisker was set at the lowest datum above Q1 - 1.5 (Q3 - Q1), and the upper whisker, at the highest datum below Q3 + 1.5 (Q3 - Q1). Data outside the whiskers was displayed as circles. The relative area error was computed as *ϵ_Auto_* = *a_Auto_*/*a_ht_* – 1 and *ϵ_Corrected_* = *a_Corrected_*/*a_ht_* – 1, respectively.

**Fig. 8.**
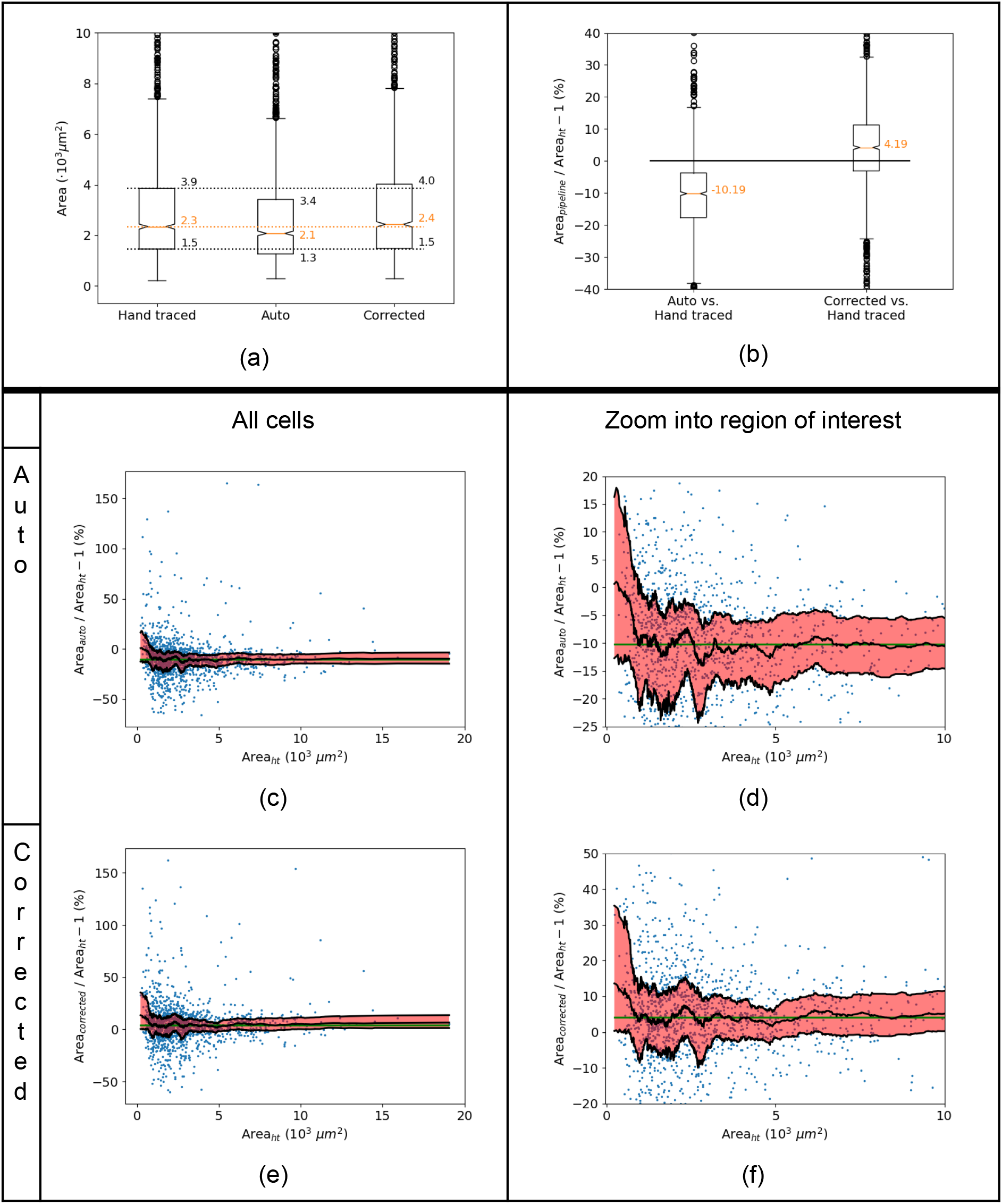
Validation of the segmentation sub-pipeline on the training data set. (a) Comparison of cell area distribution between the hand-traced data set, the Auto and Corrected automatically segmented labels. Only matches with *DC_Auto_* > 0.5 were accepted. (b) Relative area segmentation error with respect to hand traced cells. (c)-(f) Segmentation error as a function of hand traced cell area (c: Auto, e: Corrected). Zoom into regions of interest for Auto (d) and Corrected (f). Blue dots correspond to individual cells in the training data set. Black solid curves represent the HD quartiles (Q1, Q2, Q3) on points sorted by Area w, with region between the curves highlighted as a shaded red area. Green horizontal line represents the overall HD Q2 for all cells.

Although they are the standard, summary statistics hide important subpopulation (i.e. cells of a similar size) information. Ideally, we would like a constant relative error for all cell subpopulations; otherwise, segmentation errors that affect cells of a certain size but not others could be erroneously interpreted as phenotypes. To assess this for our algorithm, we plotted Auto and Corrected relative errors vs. hand traced cell area. We then sorted the errors by hand traced cell area and computed rolling Harrell-Davis (HD) estimates for quartiles={Q1, Q2, Q3} of the relative errors using a rolling window of 100 points that shrinks as it overflows at the extremes down to a minimum size of 20 points. The rolling median and interquartile range Q1-Q3 were plotted as a solid curve and red shaded area, respectively (Fig. 8c-f). Finally, we computed the global HD estimate for Q2 and plotted it as a horizontal green line.

### 2.6. Phenotype framework for WAT

We propose a novel phenotype framework for WAT based on linear models to relate body weight (BW), depot weight (DW) and cell area quartiles, described in this section. We also compare with two traditional approaches: cell area probability density functions (pdfs) to represent cell populations and Tukey’s HSD test for mean differences.

Previous *KLF14* phenotype studies stratified data sets by sex (Glastonbury et al., 2020; Small et al., 2018) and compared two groups (sex or risk allele vs. non-risk allele) with summary statistics, namely median adipocyte area with a Wilcoxon signed-rank test (Small et al., 2018), or mean area with inverse variance fixed effects meta-analysis (Glastonbury et al., 2020). Neither adjusted for BW (the Wilcoxon signed-rank test compared pairs of BMI-matched subjects and the meta-analysis study selected subjects within the normal BMI range), although BW is a known general phenotype confounder (Karp et al., 2012; Oellrich et al., 2016), in particular of mouse adipocyte diameter and DW (van Beek et al., 2015). In addition, summary statistics could miss changes in cell subpopulations that become apparent when comparing population quantiles (Rousselet et al., 2017).

To tackle equivalent issues in the mouse model, we propose a *Klf14* phenotype framework with three levels interconnected via linear models: the mouse level (body weight, BW), the depot level (depot weight, DW) and the cell level (quartile cell area). We study cell quartile sizes, but other quantiles could be used too. In each level, the linear model quantifies a trait (BW, DW or cell area) vs. categorical effect (genotype) and continuous covariate (BW or DW). This builds upon the mixed model proposed by (Karp et al., 2012; Oellrich et al., 2016) (trait ~ genotype*sex + body_weight + (1|batch)) and adipocyte size or DW vs. BW non-linear models by (van Beek et al., 2015). Our approach is different in six ways: 1) as we have a smaller number of animals per stratum, we did not consider “batch”, and thus could use simpler Ordinary Least Squares (OLS) models. In larger studies, batch and other random effects (litter, mother) could be considered. 2) As sexual dimorphism would dominate other effects, we used separate models for females and males. 3) We combined the heterozygous parent of origin for the KO allele (father, PAT or mother, MAT) and genotype (wildtype, WT or heterozygous, Het) into a categorical effect, referred to as “genotype” for simplicity. We performed exploratory analysis of three genotype categories: Control (PAT WT and PAT Het), MAT WT and functional knockout or FKO (MAT Het) —we pooled PAT WT and PAT Het mice because *KLF14* is only expressed from the maternally inherited allele (Parker-Katiraee et al., 2007) and this reduces the number of tests. 4) We considered two covariates to adjust for, BW and DW, instead of just BW. 5) We added an interaction term between genotype and covariate to allow for different slopes for different genotypes. 6) We used adipocyte area instead of adipocyte diameter (van Beek et al., 2015) to make the relationship with DW more linear.

Accordingly, the three-level linear models we propose are: (BW ~ genotype), (DW ~ genotype * BW/<u>BW</u>) and (area_q_ ~ genotype * DW), where 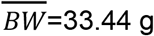 is the mean BW of all animals and q={Q1, Q2, Q3} are quartiles. BW, DWand area_q_ are continuous variables, and genotype={Control, MAT WT, FKO} is a categorical variable. In addition, a constant term is included in all models, but omitted in the formula for simplicity. The main difference with studies that adjust additively for BMI = BW/ height^2^ (Glastonbury et al., 2020; Honecker et al., 2021), with models like (cell size ~ variables + BW/ height^2^) is our interaction term and that our intermediate DW level between cell area and BW. BWand DWwere measured with Sartorius BAL7000 scales. For cell area quantification, we applied DeepCytometer to 75 inguinal subcutaneous and 72 gonadal full histology slides (with one or two tissue slices each), including the 20 slides sampled for the hand-traced data set. Each slide belongs to an animal and depot (gonadal or subcutaneous), stratified by sex —female (f) or male (m)—, and genotype (PAT WT, PAT Het, MAT WT, MAT Het). For segmentation of training slides, we used the pipeline instance that did not see any part of it in training. Slides not used for training were randomly assigned to one of the 10 pipeline instances.

We extract two results from the linear models. First, their slopes as ratios between a trait and BW or DW, and slope two-tailed t-tests for significance of association or correlation (due to the equivalence with Pearson’s coefficient t-test). Second, Likelihood Ratio Tests (LRTs) assess whether the fit with three genotype categories (Control, MAT WT, FKO) is significantly better than with two categories pooled together (e.g. Control + MAT WT, FKO). If so, the pooled models are considered significantly different as a whole. When fitting multiple linear models, e.g. 12 models for 2 sexes, 2 depots and 3 genotypes, the p-values of all slopes where corrected using the FDR two-stage step-up Benjamini-Krieger-Yekutieli method (Benjamini et al., 2006) for a significance level α=0.05.

To check whether linear models offer more insight into the data than mean differences tests, we also applied Tukey’s HSD test to the three genotype={Control, MAT WT, FKO} categories for a family-wise error rate FWER=0.05 to test for differences in BW, DW and cell size.

In addition, we computed cell area pdfs, by applying Kernel Density Estimation with a Gaussian kernel and bandwidth = 100 μm to preserve sharp peaks in the distributions. Quartiles (Q1, Q2, Q3) or other quantiles were computed with Harrell-Davis (HD) quantile estimation (Harrell and Davis 1982). For a weighted average of quantiles from multiple mice, for each p-quantile for the i-th mouse q_i_(p) we computed the corresponding HD standard error se_i_(p), using jackknife and applied the meta-analysis inverse-variance method (Cochran, 1954, 1937). The combined quantile estimate is

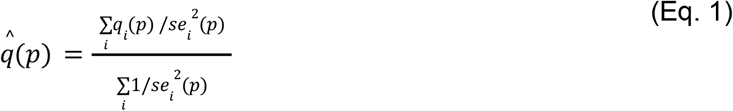

and the combined standard error is

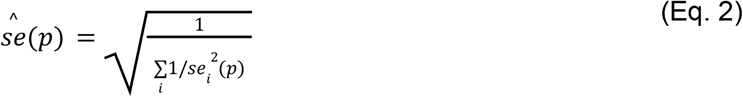

The 95%-CI for the combined quantile estimate is

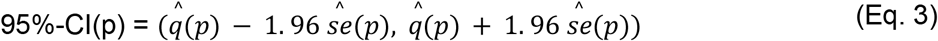

### 2.7. Quantile colour map plots to assess cell size heterogeneity

Because of skewness and long tails in the histogram (Suppl. Materials Fig. S1a and S3a), overlaying a colour map proportional to cell area *a_i_* on the tissue sample produces a low contrast image that offers very little visual information about spatial patterns. To overcome this, we propose a colour map in Hue/Saturation/Lightness (HSL) mode, where hue is proportional to area quantile *q_i_* computed from an ECDF of all segmented cells

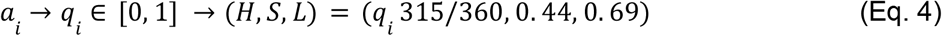

We create a the scatter map (*x_i_*, *y_i_*) → *a_i_*, where (*x_i_*, *y_i_*) is the *i*-th white adipocyte centroid, and use Delaunay triangulation and linear interpolation to rasterise the scatter map to all pixels that belong to white adipocytes. Because of sexual dimorphism, we created a colour map for each sex.

## 3. Results

In this section we present the DeepCytometer pipeline and its validation, followed by a novel analysis of *Klf14^tm1(KOMP)Vlcg^* C57BL/6NTac mouse WAT from the (Small et al., 2018) study, as described in Section 2.1. This analysis is based on body and depot weights, and cell areas derived from automatic segmentations by our pipeline.

### 3.1. DeepCytometer pipeline

A main contribution of this paper is our DeepCytometer pipeline to segment white adipocytes from whole H&E histology slides. The pipeline (Fig. 7a) performs a coarse segmentation of the tissue, uses an adaptive tiling algorithm to select image blocks that fit in GPU memory and processes each block with a white adipocyte segmentation sub-pipeline (Fig. 7b) based on DCNNs. The sub-pipeline works by segmenting all objects in the image block, then classifying which ones are white adipocytes, and correcting their outlines to account for cell overlap. Pixels that belong to cropped cells on the edges are flagged to be processed in neighboring image blocks. A detailed description of slide preprocessing can be found in Section 2.3. The segmentation sub-pipeline is described in Section 2.5, with details of its constituent DCNNs, in Section 2.4.

The pipeline experiments we present below quantify the reduction in the number of pixels that need to be processed with our adaptive tiling algorithm compared to uniform tiling; and validate the Tissue CNN and segmentation sub-pipeline.

#### 3.1.1. Adaptive tiling computational load reduction

We measured the reduction of computational load provided by our adaptive tiling compared to uniform tiling of the tissue region with overlapping blocks, by comparing the number of pixels each approach needs to process. Uniform tiling was produced by splitting the image into (L_max_, L_max_) square blocks, where L_max_=2,751 pixels is the maximum tile length allowed in our adaptive algorithm. Blocks overlapped by 2 *R_max_* + *ERF* on each side, where *R_max_* = 179 pixels is the radius of the largest circular cell accepted by the pipeline, and *ERF* = 131pixels is the maximum Effective Receptive Field of the CNNs. The sum *A_total,uniform_* of the areas of all the uniform blocks containing tissue in an image were compared to the sum *A_total,adaptive_* of the areas of the adaptive blocks (Fig. 9). The average ratio from the 147 whole slides used in the phenotyping experiments was *A_total,adaptive_/A_total,uniform_* = 0.86 ± 0.13, corresponding to a reduction of 16.59% in the number of processed pixels (and correspondingly, time), from an average of 2.11 · 10^-9^ pixel (uniform) to 1.81 · 10^-9^ (adaptive) per slide.

**Fig. 9.**
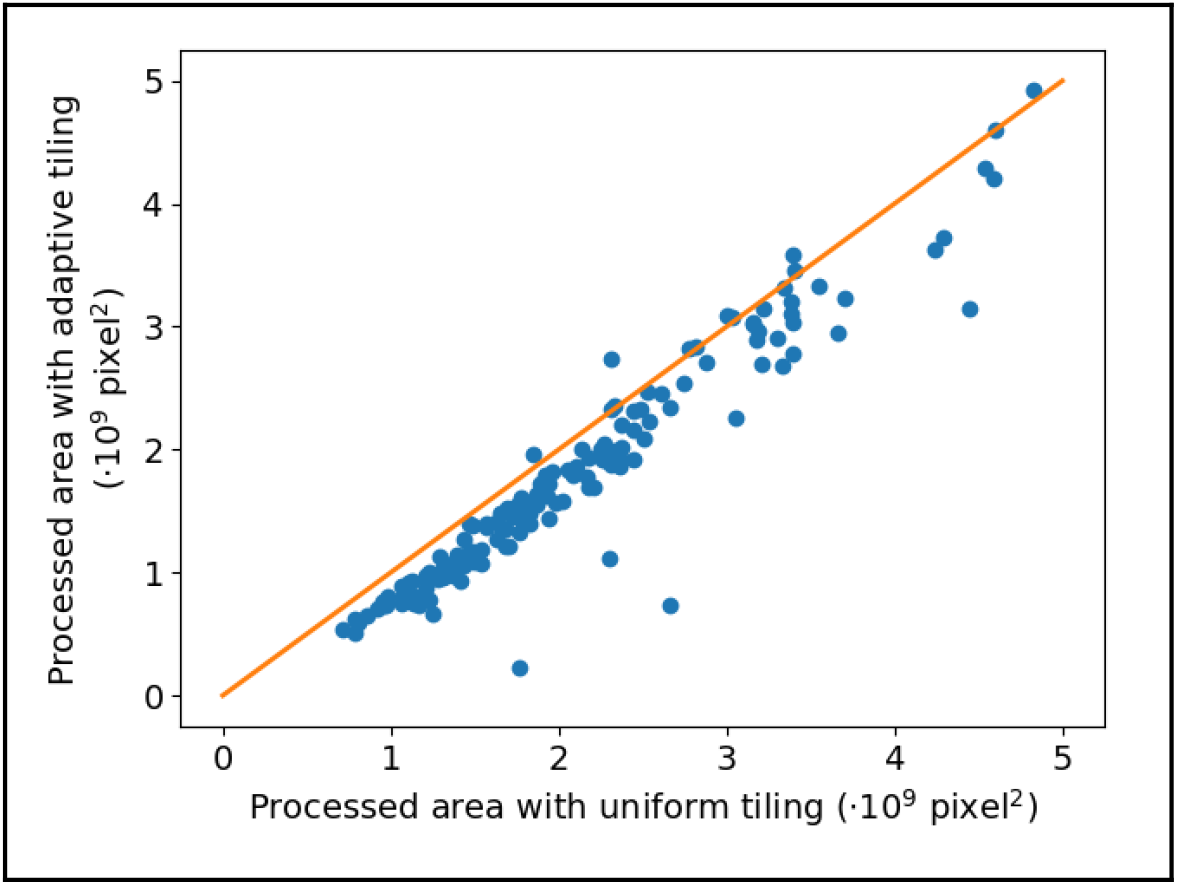
Comparison of total area processed by uniform tiling of histology images vs. our adaptive tiling. Each point corresponds to one histology image. The orange line is the identity line.

#### 3.1.2. Tissue CNN validation

The Tissue CNN was validated on 126 training images using 10-fold cross validation. We calculated the receiver operating characteristic (ROC) curve for the classification of white adipocytes vs. “other” objects, weighted by the number of pixels in each object (experiment details in Section 2.5, curve in Fig. 10a). The classifier performs very well, with area under the curve = 99.59%, and pixel-wise false positive rate (FPR) = 1.80% and true positive rate (TPR) = 97.71% for a white-adipocyte classification threshold *Z_obj_* ≥ 0.5. The low FPR means that cell population studies will contain a negligible number of false objects, and the high TPR indicates that the vast majority of white adipocytes will be detected in the slides. We also provide TPR and FPR values for other thresholds in Fig. 10b.

**Fig. 10.**
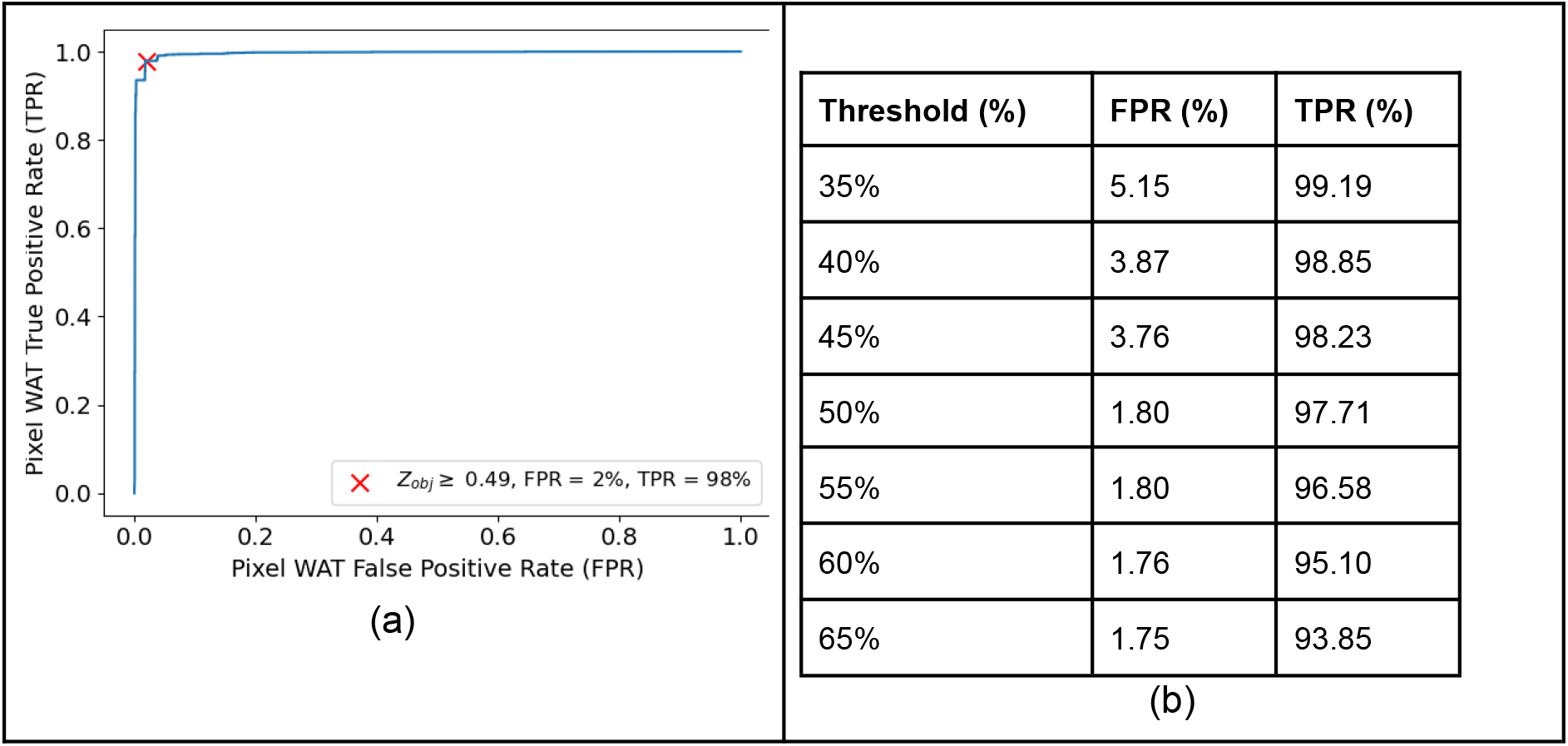
Tissue CNN classification validation. (a) Receiver Operating Characteristic (ROC) curve. (b) Some ROC curve numerical values. Object classification error (white adipocyte vs. non white adipocyte) weighted by number of pixels. (Weighting used as the hand traced data set contains many more white adipocyte objects, but non white adipocyte objects can be very large.)

#### 3.1.3. Segmentation sub-pipeline validation

We validated the segmentation sub-pipeline on 55 hand-segmented images using 10-fold cross validation, both for the Auto and Corrected methods (experiment details in Section 2.5). We computed the Dice Coefficient (DC) between pairs of DeepCytometer and hand-traced (ht) cell contours, matched as described in Section 2.5.6. For the Auto method we obtained DC_Auto_ = 0.89 (median), 0.85 (mean), 0.10 (std), and for the Corrected method, DC_Corrected_ = 0.91 (median), 0.87 (mean), 0.10 (std). We also computed the median relative errors, which were −10.19% (Auto) and 4.19% (Corrected) (Fig. 8b). Both DC and relative error suggest that DeepCytometer segments white adipocytes with an acceptable area error, and that the Corrected method performs better than the Auto method.

To evaluate whether DeepCytometer segmentations represent the training cell area population, we compared box-and-whisker plots of the hand traced, Auto and Corrected white adipocyte areas (Fig. 8a). The hand traced population had quartiles (Q1, Q2, Q3) = (1.5, 2.3, 3.9) 10^3^ μm^2^. The Auto method moderately underestimated cell areas, as expected due to the lack of cell overlap, (Q1, Q2, Q3) = (1.3, 2.1, 3.4) 10^3^ μm^2^. The Corrected method approximated the hand traced population better, with just a slight overestimation (Q1, Q2, Q3) = (1.5, 2.4, 4.0) 10^3^ μm^2^.

Furthermore, as discussed in *Methods*, we went beyond summary statistics commonly found in the current literature to evaluate whether segmentation errors remain constant across the cell population. For this, we plotted Auto and Corrected relative errors vs. hand traced cell area, the rolling median and interquartile range curves and the global median relative error (Fig. 8c-f). The curves suggest that relative errors remain constant for Area_ht_ ≥ 780 μm^2^, but shift towards more positive values for smaller cells. This would suggest less reliable phenotyping results for cells with Area_ht_ < 780 μm^2^, which comprise the bottom 15.9% of the population. This highlights the need to report segmentation errors by cell size in future literature.

### 3.2. White adipocyte population study using hand traced and DeepCytometer segmentations

In this section, we calculate probability distribution functions (pdfs) from the hand traced data set and the DeepCytometer segmented whole slides to compare hand traced populations to DeepCytometer whole slide ones. In addition, we present heatmaps of cell areas computed from the DeepCytometer segmentations to visually assess local correlations and whether slides from the same stratum have similar spatial distributions. We stratify the data into three genotype categories: Control (PAT WT + PAT Het), MAT WT and functional knockout or FKO (MAT Het).

#### 3.2.1. Area population distributions of hand traced cells

The hand traced data set consists of 1,903 cells pooled from 60 subcutaneous windows and 20 mice (see Section 2.1). (Note that the hand-traced data set only included subcutaneous slides from Control and FKO mice.) The cells measured between 66 μm^2^ (321 pixel) and 19,058 μm^2^ (92,544 pixel). To represent cell populations, we estimated probability density functions (pdfs) of the areas of hand traced cells (Fig. 11a). We also calculated the cell area Harrell-Davis (HD) quartiles (Q1, Q2, Q3) with 95%-CIs (Fig. 11b). Male cells were notably larger than female cells for each quartile. However, there were no significant differences between Control and FKO area quartiles, with overlapping 95%-CIs (although there is a trend for smaller FKO cells).

**Fig. 11.**
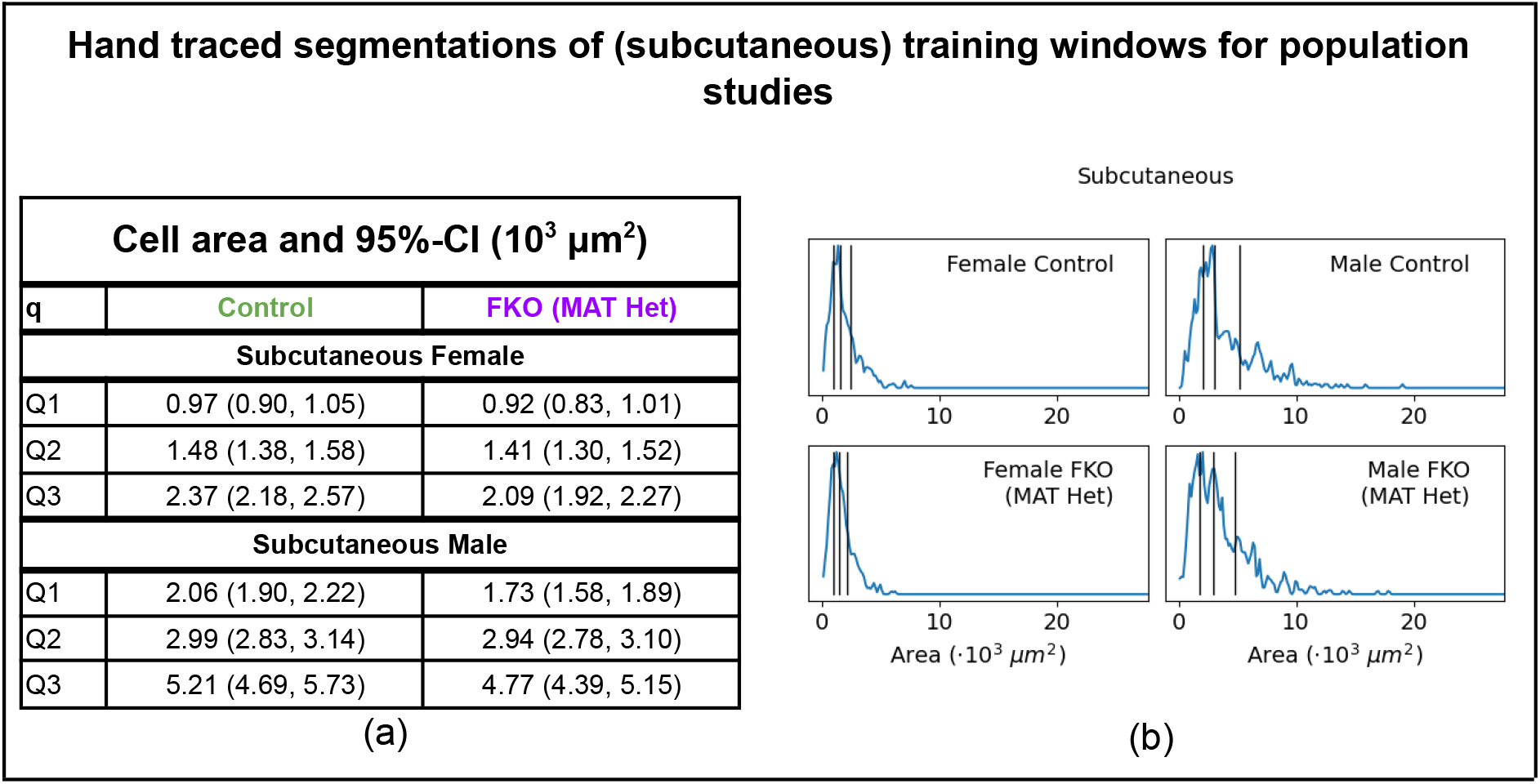
Cell populations of the hand traced data set stratified by sex, depot and genotype. (a) Harrell-Davis (HD) quartile estimates and their 95%-CIs. (b) Kernel Density estimation of cell probability distribution function (pdf, blue curve) and HD quartiles q={Q1, Q2, Q3} (vertical black lines).

#### 3.2.2. Comparison of hand traced vs. DeepCytometer segmented cell populations

In Section 2.5, we showed that DeepCytometer automatic segmentation accurately approximates hand traced contours. In this section, we test whether whole slide segmentation and segmenting more slides also add valuable population information to the hand traced data set. For that, we segmented 75 inguinal subcutaneous and 72 gonadal whole histology slides using DeepCytometer (the Corrected method), corresponding to 73 females and 74 males, to produce 2,560,067 subcutaneous and 2,467,686 gonadal cells. We estimated one pdf per slide using Kernel Density Estimation (Fig. 12a-b), and combined all pdfs in a stratum by computing pdf HD quantiles 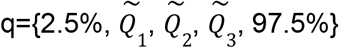 (i.e. density quantiles instead of area quantiles) to summarise the pdf information. We display 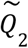 as a solid curve in Fig. 12c-d, the 2.5%-97.5% interval as a light shaded area and the interquartile range 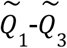 as a dark shaded area. In addition, we computed the cell area HD quartiles Q1, Q2, Q3 and their standard errors for each pdf. We combined those estimates using the inverse-variance meta-analysis method to produce combined quartiles, 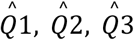, and their 95%-CIs per stratum. The combined 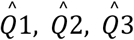 are displayed as vertical black lines in Fig. 12c-d, and their numerical values and 95%-CIs are provided in Fig. 13. These CIs are very narrow due to the large number of correlated size values. We also computed similar combined pdfs and quartiles from the subset of 20 subcutaneous Control and FKO whole slides that training windows were extracted from (Fig. 14).

**Fig. 12.**
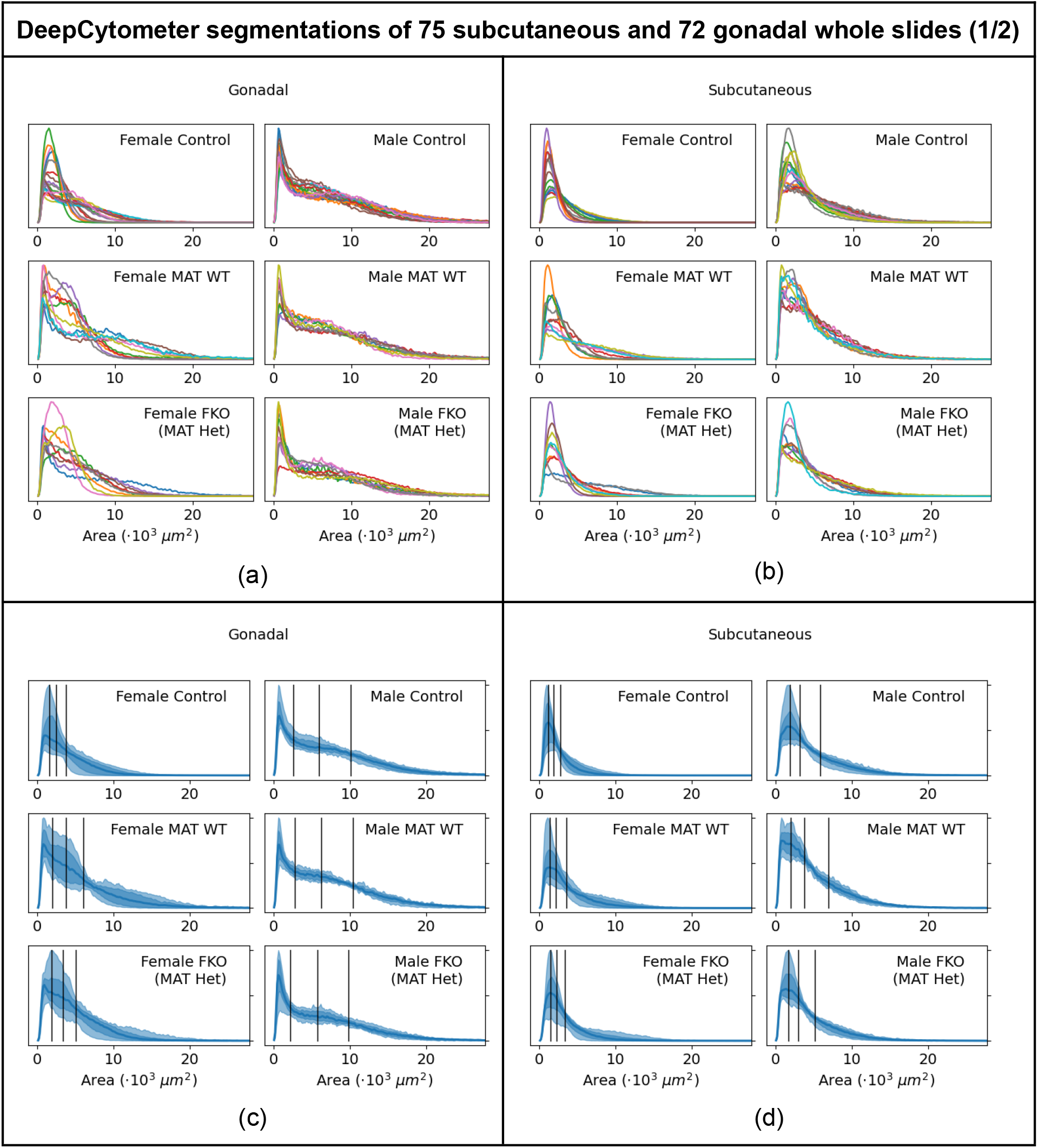
Cell populations of DeepCytometer segmentations of 75 subcutaneous and 72 gonadal whole slides, stratified by sex, depot and genotype. (a)-(b) Pdfs from Kernel Density Estimation, one curve per mouse. (c)-(d) Pdf quantiles (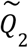 as solid blue curve, 2.5%-97.5% interval as a light shaded area, 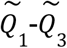 interval as a dark shaded area) and cell area quartiles (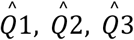 as vertical black lines).

**Fig. 13.**
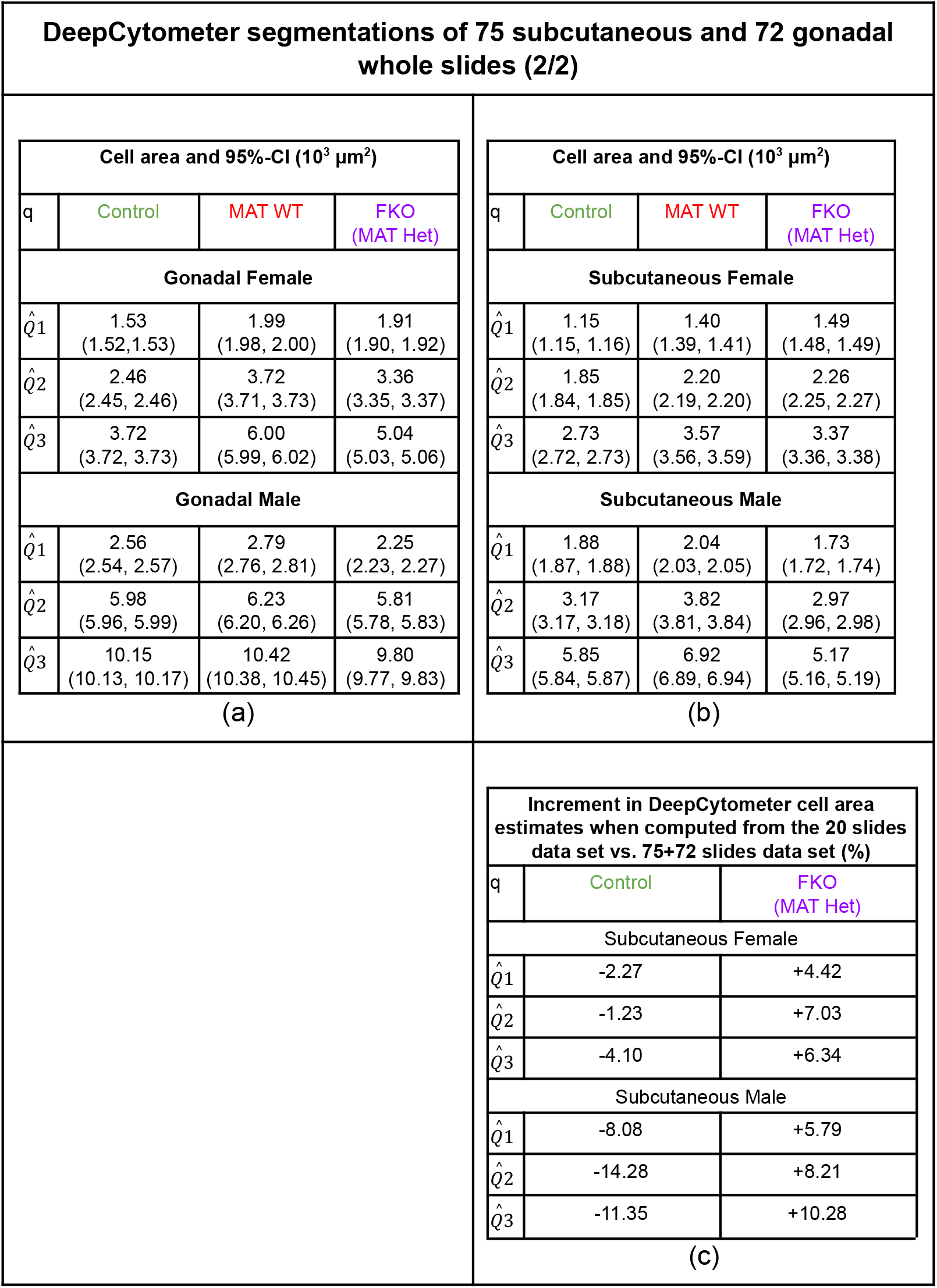
Cell populations of DeepCytometer segmentations of 75 subcutaneous and 72 gonadal whole slides, stratified by sex, depot and genotype. (a)-(b) Combined cell area Harrell-Davis (HD) quartiles 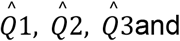 their 95%-CIs. (c) Increment in cell area estimates from the 20 slides data set in Fig. 14c vs. 75+72 slides data set in this figure’s (a)-(b).

**Fig. 14.**
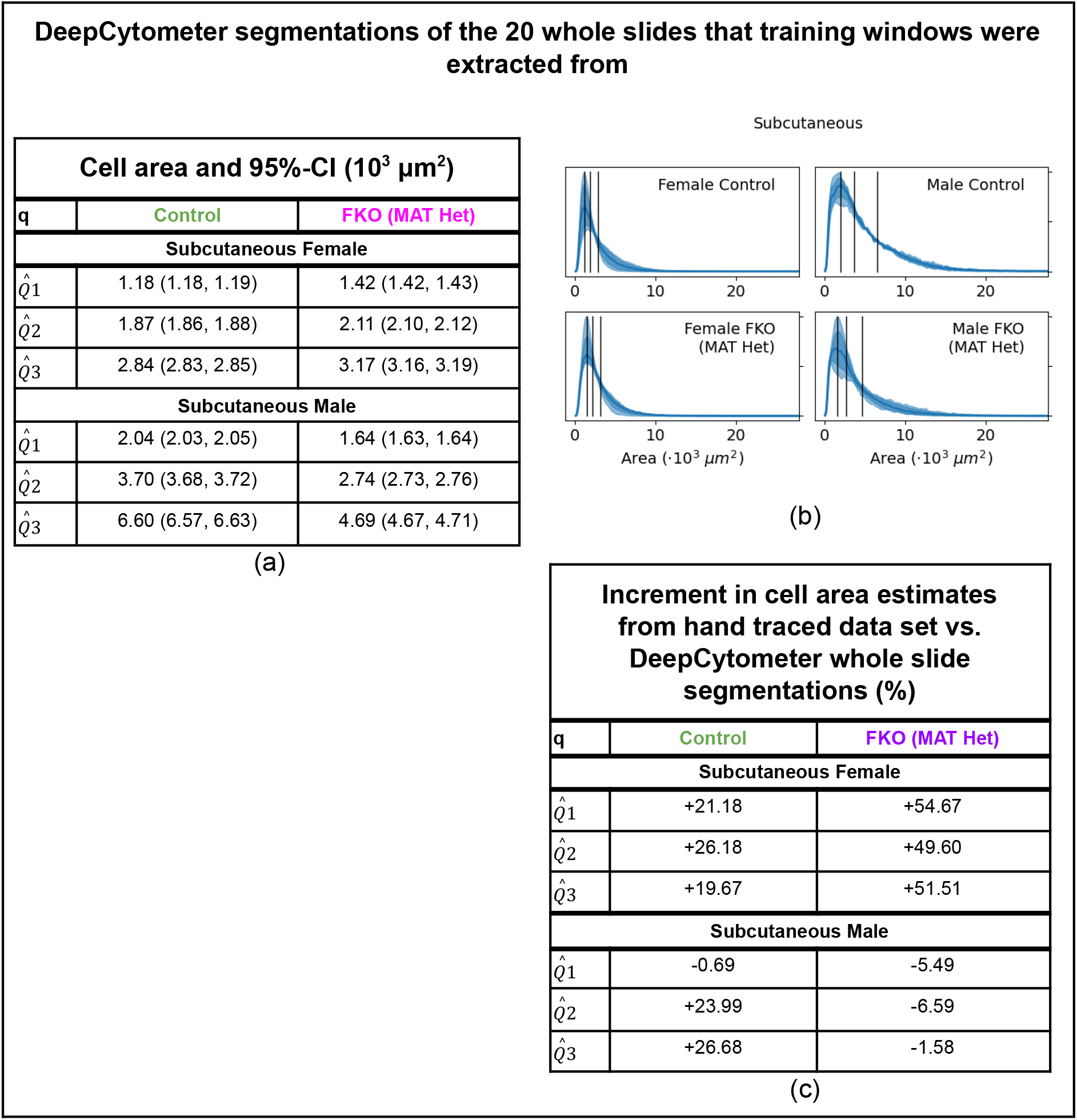
Cell populations of DeepCytometer segmentations of the 20 whole slides that training windows were extracted from, stratified by sex, depot and genotype. (a) Combined cell area Harrell-Davis (HD) quartiles 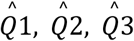 and their 95%-CIs. (b) Pdf quantiles (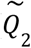 as solid blue curve, 2.5%-97.5% interval as a light shaded area, 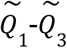 interval as a dark shaded area) and combined cell area quartiles (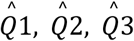 as vertical black lines). (c) Increment in cell area estimates from hand traced data set in Fig. 11b to DeepCytometer whole slide segmentations in (a).

We used this latter subset to compare populations estimated from whole slides (Fig. 14) and from a sample of windows (Fig. 11). The area quartile difference (%) between both analyses is tabulated in Fig. 14c. Although the area quartile difference between data sets is small for male FKOs (between −1.58% and −6.59%), in female FKOs it varies between +49.60% and +54.67%. In the Control strata, the difference is up to +26.68%. This suggests that our 1,903 hand traced cells from 60 windows and 20 mice, despite being a rather large data set, misrepresents the whole slide cell area populations, by undersampling the long tails on the right-hand side of the pdfs, i.e. the larger cells in the population. These differences are not systematic, and vary between strata, which would cause Type I and Type II errors in phenotype assessment. This could be due to the fact that the hand traced data set samples a relatively small number of cells per mouse and pools them, and highlights the need for whole slide analysis.

Furthermore, we compared the pdfs from the 20 subcutaneous slides to the pdfs of the full 75 slides data set (Fig. 13). The area quartile differences between both subcutaneous data sets is shown in Fig. 13c. These differences are not negligible, as the area quartile estimation error is negative for Controls and positive for FKOs. For instance, the 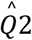 estimate in females changes by −1.23% for Controls and +7.03 for FKOs, and in males, −14.28% for Controls and +8.21 for FKOs. This would increase the chances of Type II errors or failing to detect phenotypes.

Therefore, both whole slide analysis and the analysis of more mice significantly changed the cell population pdfs. Both increases are enabled by DeepCytometer’s segmentation.

#### 3.2.3. Quantile colour map plots to assess cell size heterogeneity

In order to gain insight into the spatial distribution of adipocyte populations, we used the area-to-colour map described in Section 3.2.3 to visualise cell area distribution in all whole slides processed by DeepCytometer, both in AIDA to visually assess the segmentation (Fig. 15), and to generate figures for this paper (we show some heatmap examples in Fig. 16a-e; refer to Appendix A for all slides). The colour map is linear with area quantile, rather than area, and we use separate colour maps for females and males, due to sexual dimorphism. The results clearly show local subpopulations or clusters of white adipocytes. These clusters are irregular in shape, and present high inter- and intra-slice variability, even within the same sex and depot stratum. They illustrate the challenges for statistical studies of cell populations performed on subsamples of whole slides, such as our hand traced data set. Namely, cells within clusters are correlated observations, so although spatial analysis is without the scope of this paper, we conjecture that an apparently large number of cells (~2,000 in our hand traced data set) may not properly represent the mixture of subpopulations in the original whole slides.

**Fig. 15.**
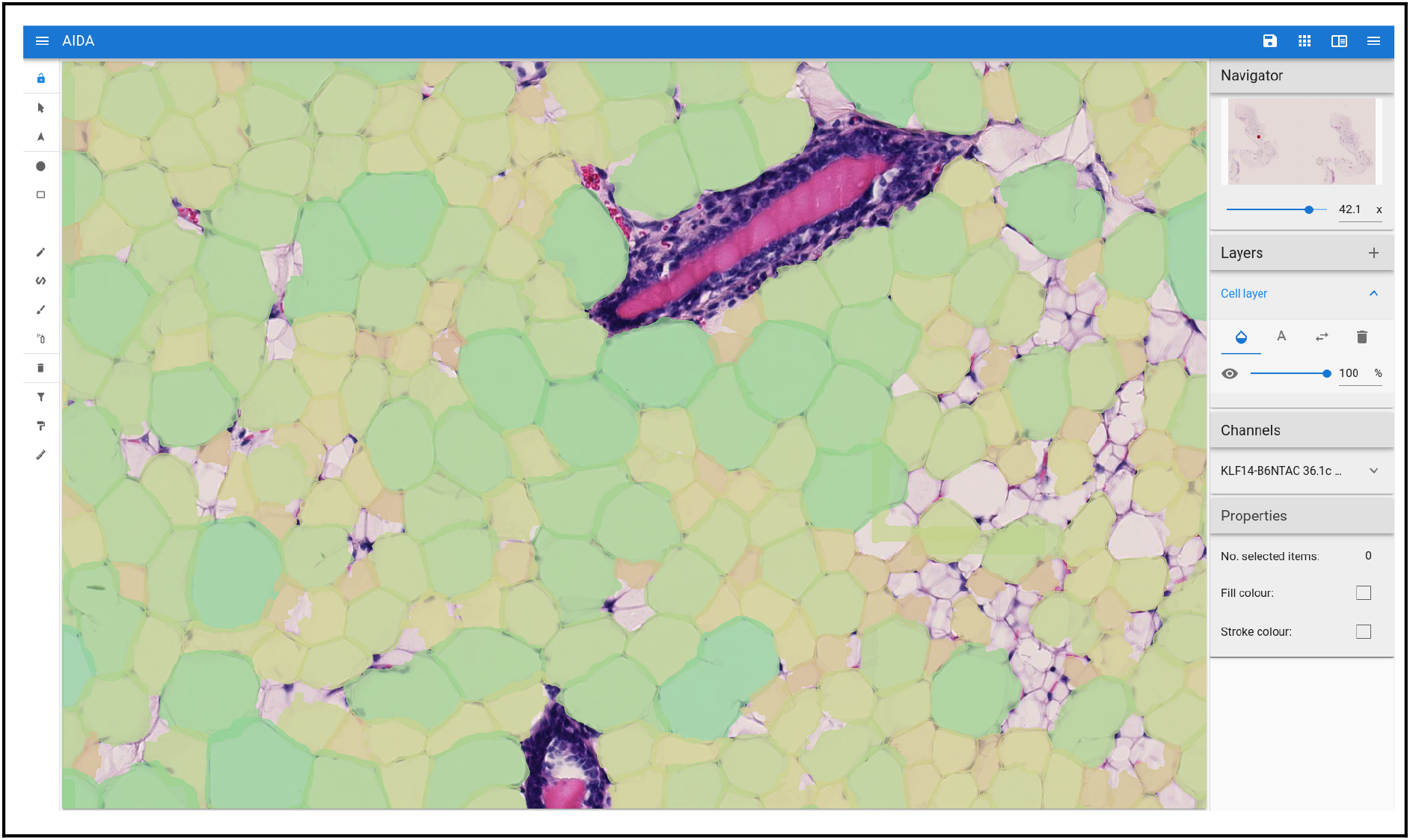
AIDA web interface with DeepZoom navigation of histology file displaying overlaic DeepCytometer segmentations, coloured according to quantile colour map.

#### 3.2.4. Application to human WAT

As a proof-of-concept of whether the pipeline can be applied to other data sets, we applied the pipeline to 347 human WAT histology slides from the GTEx project (Lonsdale et al., 2013). We filtered out segmented objects smaller than 200 μm^2^ or larger than 160,000 μm^2^, with a white adipocyte classifier score <0.5 or inverse compactness<2.0. Visual inspection of the results suggests that the pipeline can segment them similarly to mouse histology. We display four illustrative examples in Fig. 16f.

**Fig. 16.**
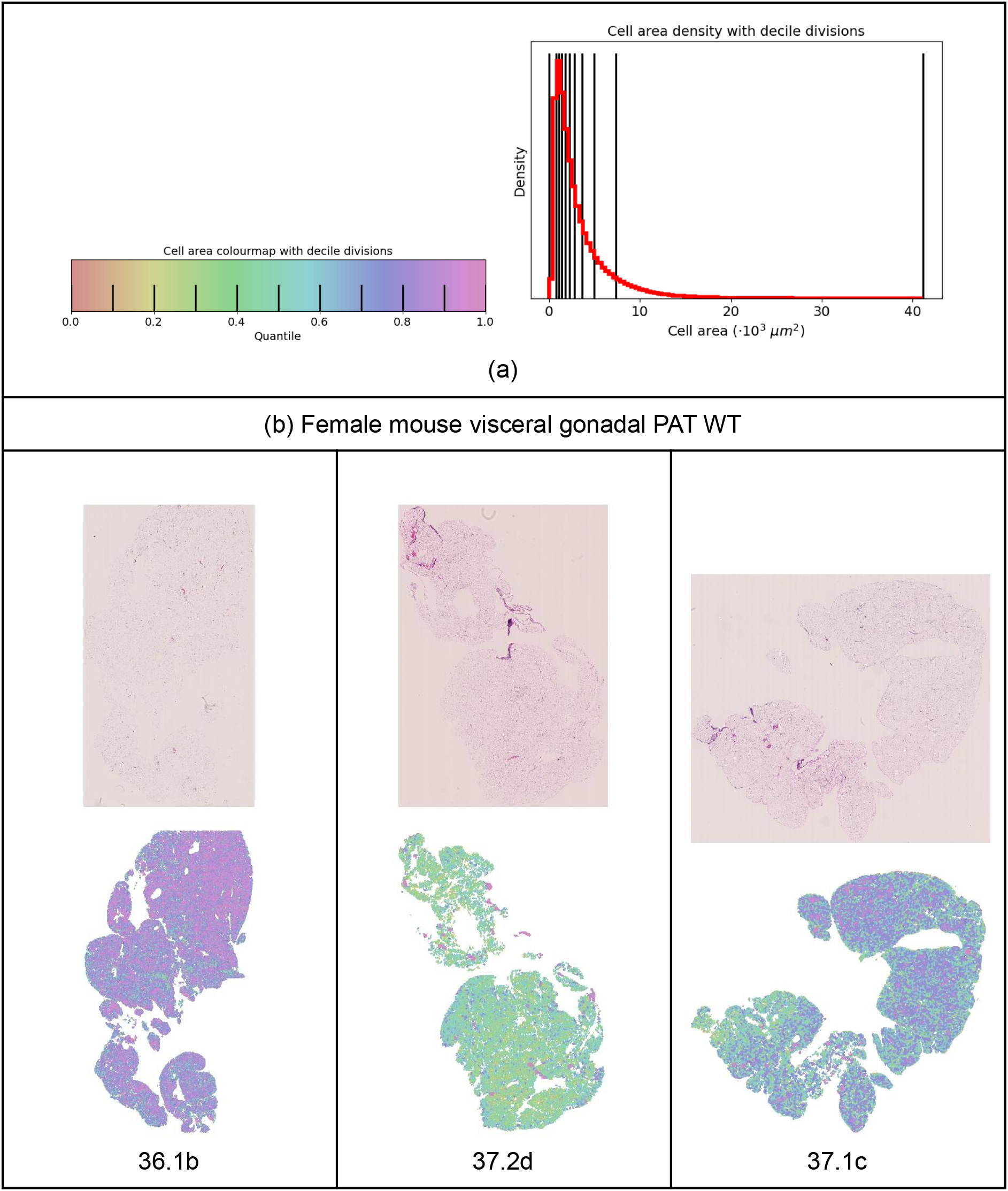
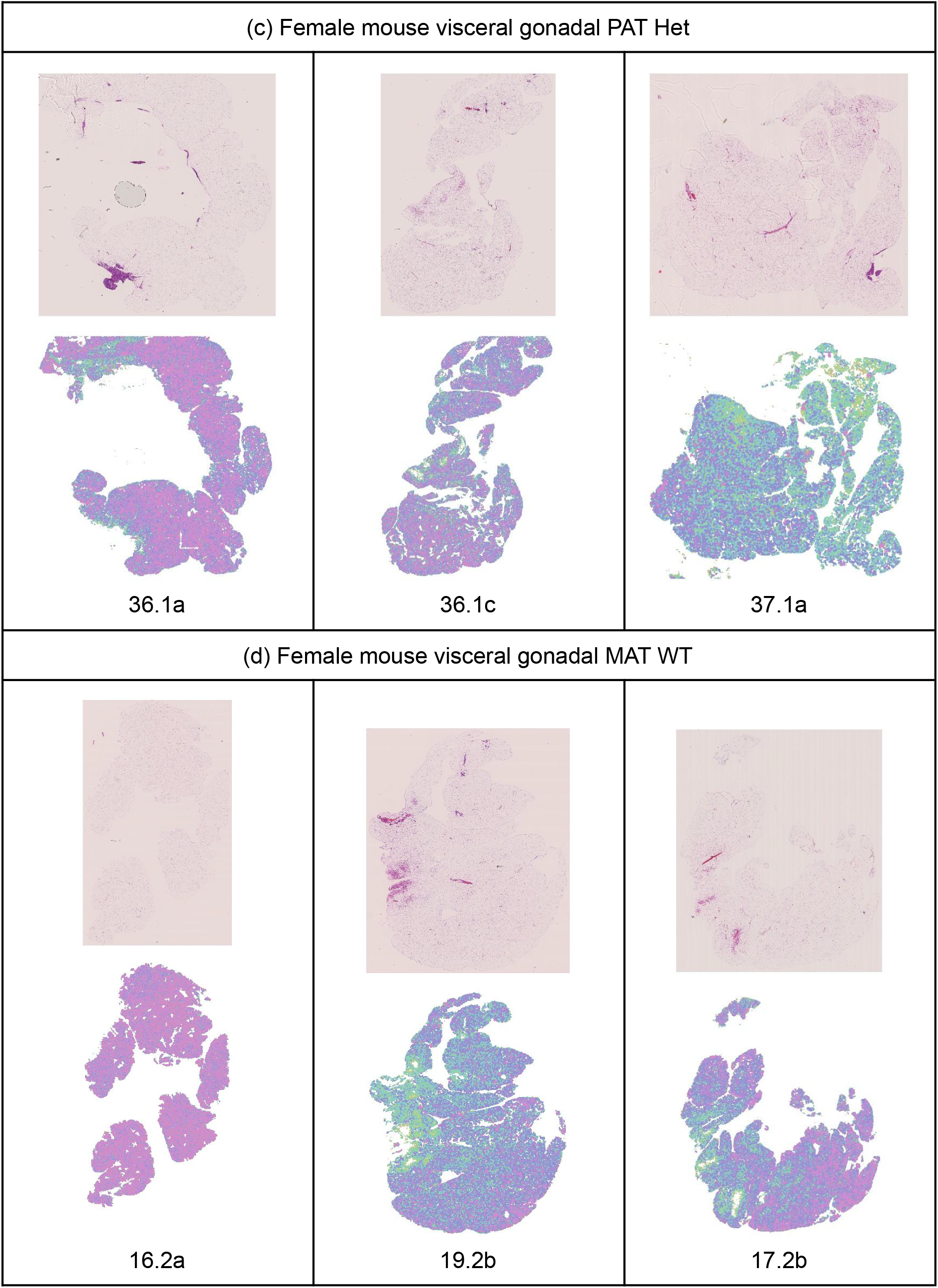
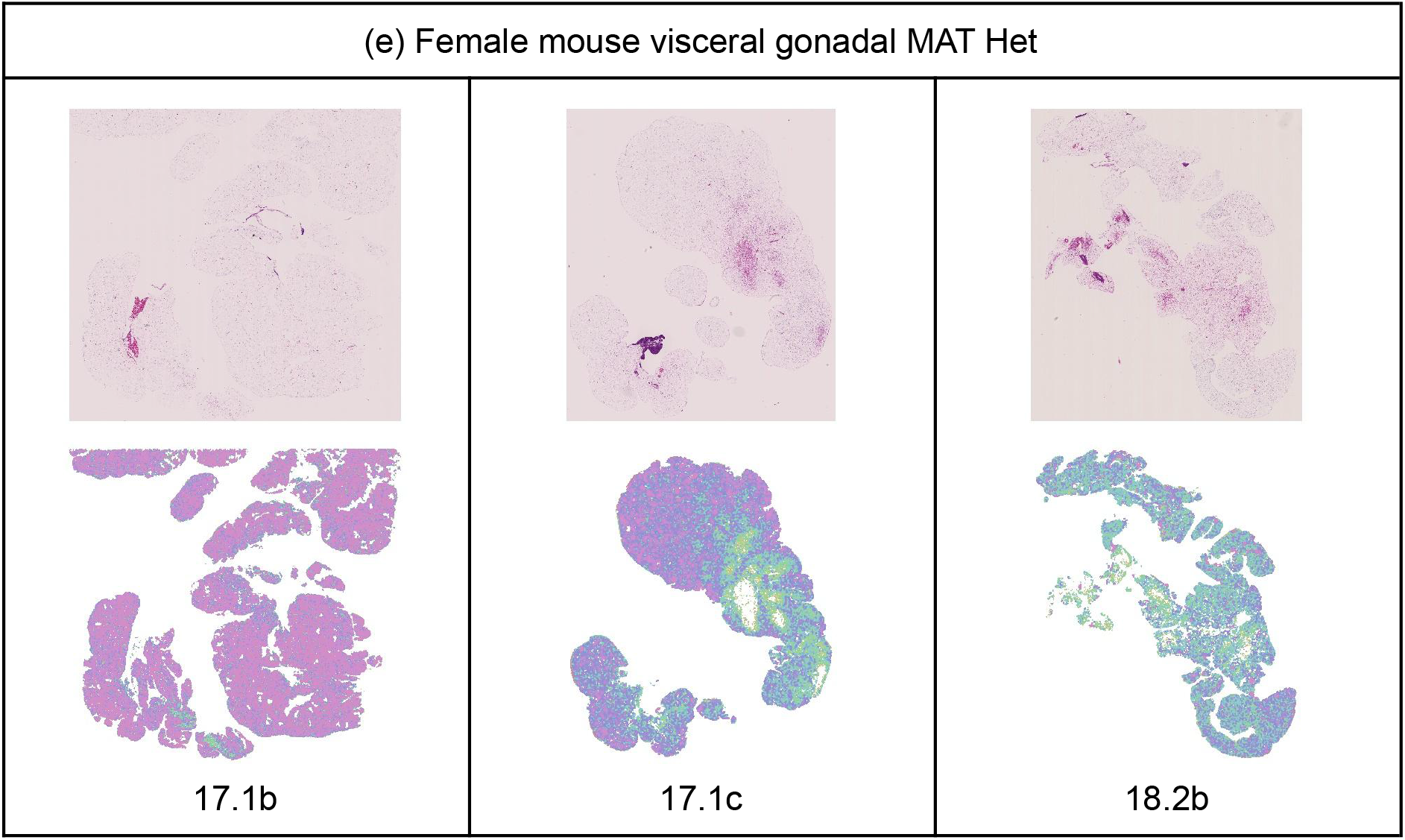
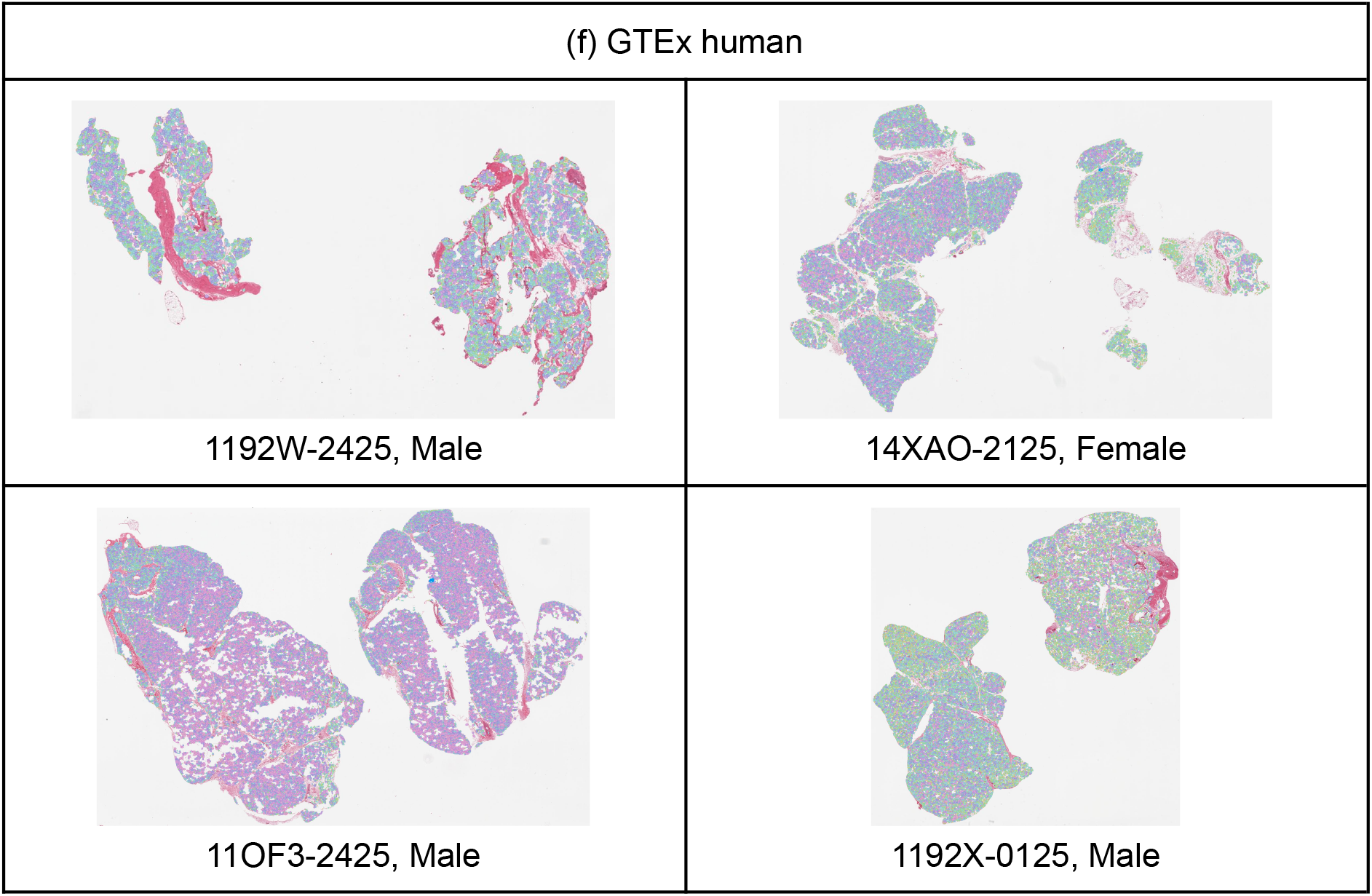
Illustrative examples of white adipocyte tissue histology and area quantile heatmaps (for female mouse visceral gonadal depot). Heatmaps for all processed slides can be found in Appendix A. (a): Quantile colour map and cell area density for female Corrected segmentation with deciles plotted for reference (vertical black lines). (b): PAT WT. Illustrative examples of white adipocyte tissue histology and area quantile heatmaps (for female mouse visceral gonadal depot). (c): PAT Het. (d): MAT WT. (e): MAT Het. Illustrative examples of white adipocyte tissue histology and area quantile heatmaps (for female mouse visceral gonadal depot). (e): MAT Het. Illustrative examples of white adipocyte tissue histology and area quantile heatmaps (for human GTEx histology). (f) Three males and one female. Images taken from the AIDA interface, with cell segmentations overlaid on histology. Colour map: male mouse from Appendix A.

### 3.3. Phenotype study of WAT using BW, DW and DeepCytometer adipocyte segmentations

We broke down the study of *Klf14* phenotypes into three interconnected levels: the mouse level (body weight, BW), the depot level (depot weight, DW) and the cell level (quartile cell area). The mice were stratified by heterozygous parent of origin (*Klf14* is only expressed from the maternally inherited allele) for the KO allele (father, PAT or mother, MAT) and genotype (WT or Het), jointly referred as genotype for simplicity. We performed exploratory analysis of three genotype categories: Control (PAT WT + PAT Het), MAT WT and functional knockout or FKO (MAT Het). BW and DW were measured in the laboratory, and cell areas were computed both for hand traced and DeepCytometer segmentations. (Details are provided in Section 2.6).

#### 3.3.1. Mouse level

At this level, we studied the sex and genotype effects on BW.

##### 3.3.1.1. Sex effect on BW

To assess the effect of sex on BW, we fitted a Robust Linear Model (RLM) (BW ~ sex) to the Control mice (n_female_=n_male_=18). The RLM was preferred to an Ordinary Least Squares (OLS) model to reduce the leverage of a large male outlier. The model calculated a mean BW for females of 25.06 g and 38.30 g for males (males 52.8% larger, β=13.24 g, p=6.00e-20). This sexual dimorphism is larger than genotype effects that we found in the next sections at the BW, DW or cell level, so we stratified the data by sex for the rest of the study.

##### 3.3.1.2. Genotype effect on BW

We computed Tukey’s HSD tests to compare mean BW among the three genotype groups (Control, MAT WT, FKO), stratified by sex (n_female_=n_male_=38). As shown in Fig. 17, female and male BWs share the same effect direction: MAT WTs are larger than Controls, and FKOs are smaller than MAT WTs but larger than Controls. However, the only statistically significant difference is for female MAT mean BW being 22.9% larger than Control’s (β=5.76 g, p=0.010).

**Fig. 17.**
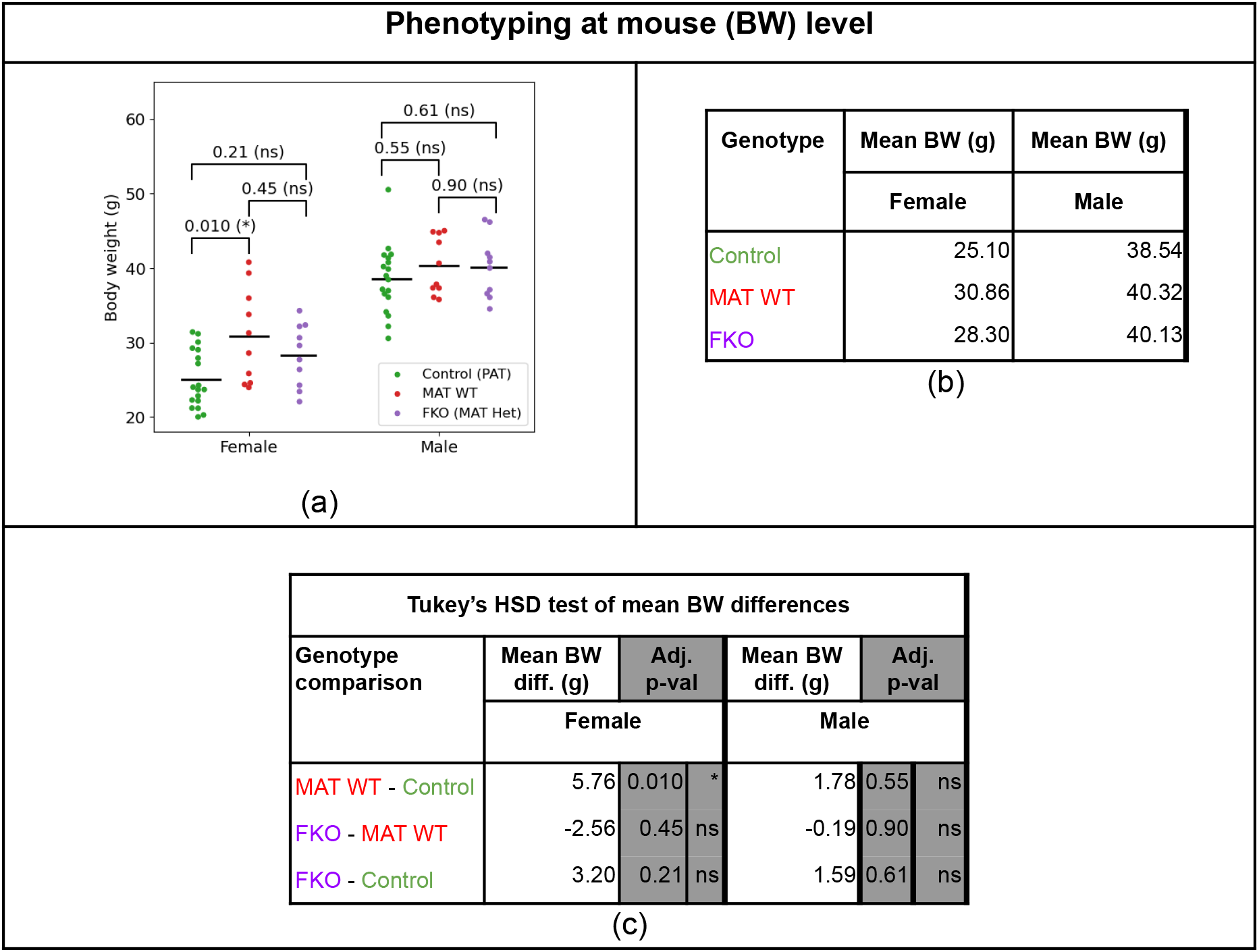
Mouse body weight (BW) phenotyping. (a) BW swarm plots stratified by sex and genotype. Each point corresponds to a mouse. Mean BW for each group provided as horizontal black line. P-values of mean differences computed with Tukey’s HSD test for each sex. (b) Table summarising mean BW stratified by sex and genotype. (c) Table summarising Tukey’s HSD test mean BW differences and adjusted p-values.

#### 3.3.2. Depot level

In this section we study genotype effects on inguinal subcutaneous (for SAT) and gonadal (for VAT) DW adjusting for BW, as well as DW correlation with BW.

##### 3.3.2.1. Linear models of DW as a function of BW and genotype

We fitted OLS models 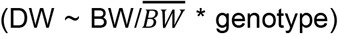 to the same 76 mice stratified by sex and depot (plots in Fig. 18a, intercept and slope values in Table 5), where 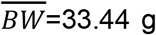 is the mean BW of all animals used as a normalisation factor to lower the condition number of the linear model, and genotype={Control, MAT WT, FKO}. We then used Likelihood ratio tests (LRTs) to test for pairwise differences between the three OLS models (Fig. 18b), and t-tests of the slopes to evaluate correlation between DW and BW (p-values from the 12 slope t-tests corresponding to 2 sexes, 2 depots and 3 genotypes, before and after Benjamini-Krieger-Yekutieli (Benjamini et al., 2006) correction are shown in Table 5; intercept p-values were not used in tests, so they did not need to be corrected).

**Fig. 18.**
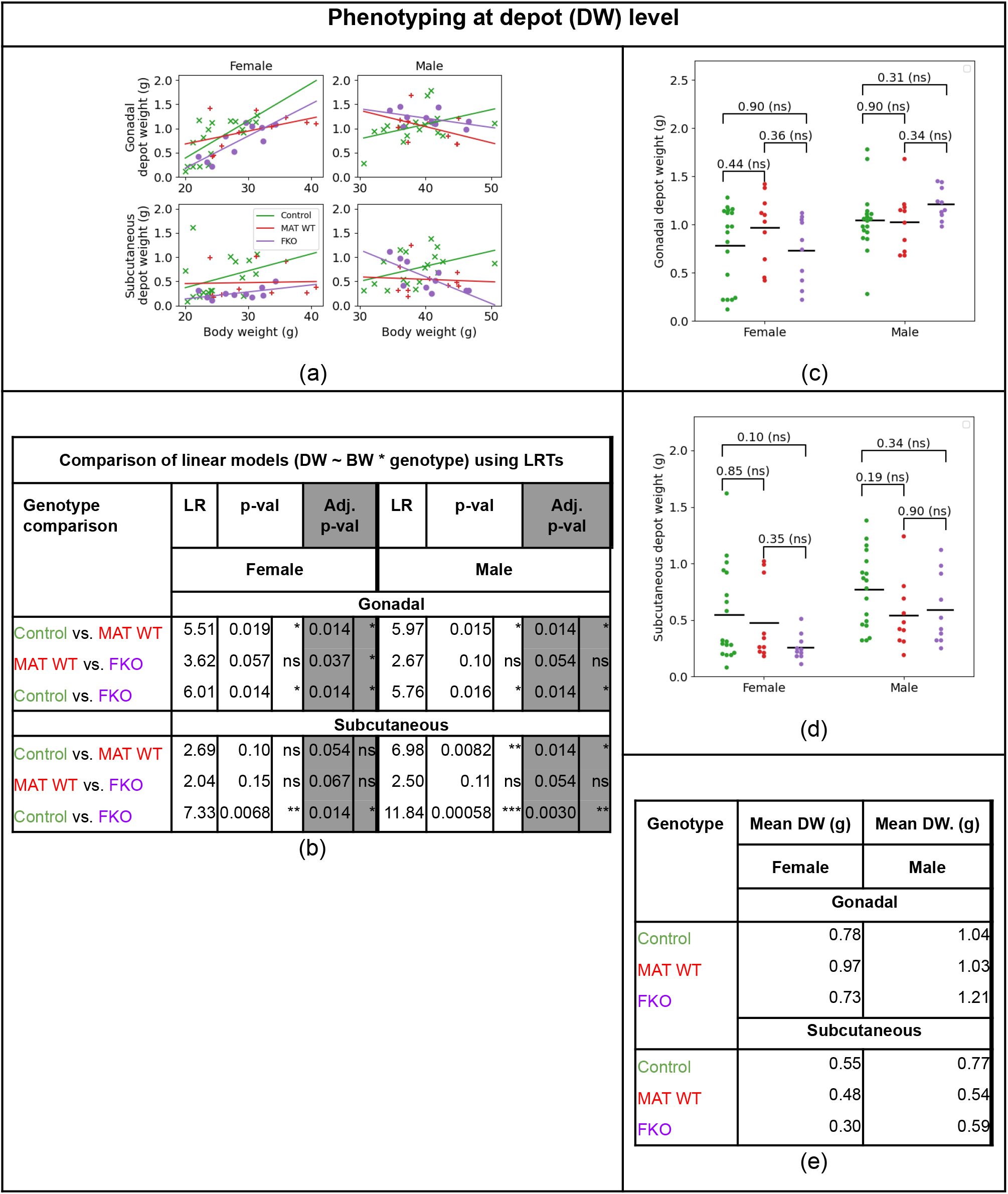
Mouse depot weight (DW) phenotyping. (a) DW OLS models 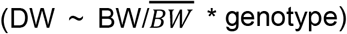, where genotype={Control, MAT WT, FKO}, stratified by sex and depot. Each point corresponds to a mouse. Intercept, slope values and t-test p-values are provided in Table 5. (b) LRT comparisons between Control, MAT WT, FKO linear models, without and with Benjamini-Krieger-Yekutieli multitesting adjustment. (c)-(d) DW swarm plots stratified by sex, depot and genotype. Mean DW for each group provided as horizontal black line. P-values of mean differences computed with Tukey’s HSD test for each sex. (e) Mean DW values stratified by genotype, sex and depot.

**Table 5.**
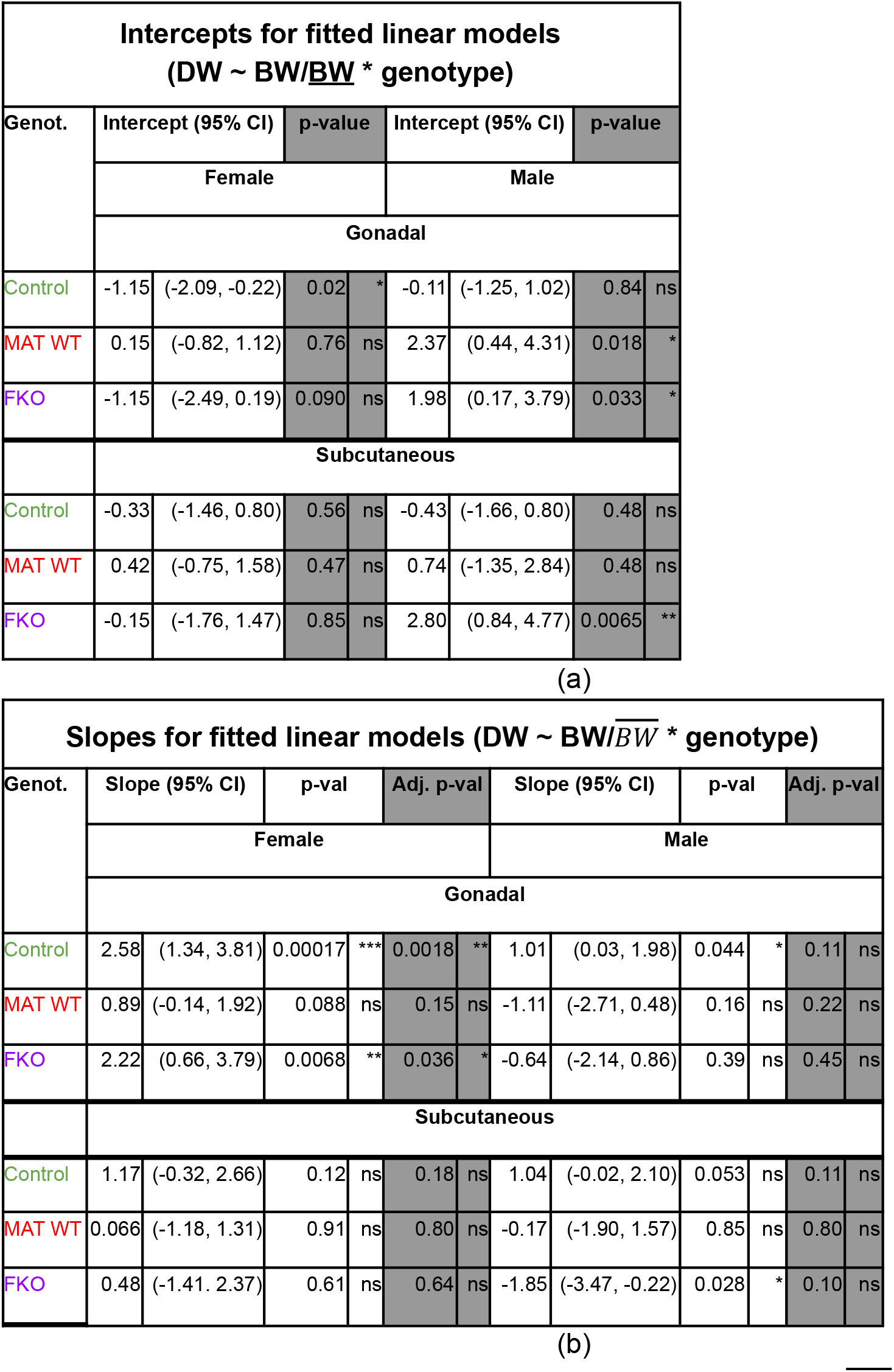
Coefficients and p-values from OLS models 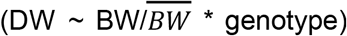, where genotype={Control, MAT WT, FKO}, stratified by sex and depot, as shown in Fig. 18a. (a) Intercept. (b) 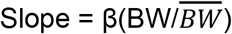.

The trend for the Control groups (both sexes and both depots) is that DW increases with BW, as expected, although the slopes are only statistically significantly ≠0 for the female gonadal stratum after correction (β=2.58, p=0.0018).

The MAT WT group, by contrast, displays flatter slopes suggesting that MAT WT DW is less correlated with BW (all slope p-values are n.s.). The difference between MAT WT and Control linear models is significant for female gonadal and subcutaneous and male subcutaneous depots (LRTs: p_female gonadal_=p_male gonadal_=p_male subcut_ =0.014), and almost significant for female subcutaneous depots (p_female subcut_ =0.054).

Finally, the FKOs show differences with Controls by sex and depot. In FKO females, the depots are smaller than for Controls (LRT Control vs. FKO: p_gonadal_=p_subcutaneous_=0.014). For instance, plugging the mean female BW, 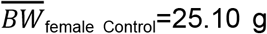 (Fig. 17b) into the Control and FKO linear models (Table 5), we obtain a predicted DW_female gonadal Control_=0.79 g vs. DW_female gonadal FKO_=0.52 g (Δ=-34.2%) and DW_female subcut. Control_=0.55 g vs DW_female subcut. FKO_=0.21 g (Δ=-61.8%). Another difference is that the FKO slopes are smaller compared to Controls, β_female gonadal Control_=2.58 vs. β_female gonadal FKO_=2.22 (Δ 14.0%), β_female subcut. Control_=1.17 vs. β_female subcut. FKO_=0.48 (Δ=-59.0%). Thus, FKO females have lower fat percentage than Control ones, and this effect is more pronounced with larger BW, and more pronounced in subcutaneous than gonadal depots. FKO males present a very different behaviour, with a trend for negative slopes (i.e. DW *decreases* with BW). But, although Control vs. FKO model differences are significant (LRT: p_male gonadal_=0.014, p_male subcut.I_=0.0030), it should be noted that all male Control and FKO slopes are non significant. Thus, more data would be necessary to decide whether the crossing of male Control and FKO linear models means that smaller FKOs actually tend to have more fat percentage than Controls, and larger FKOs tend to have less, or the negative slope is a small data set artifact.

##### 3.3.2.2. Tukey’s HSD test comparison of mean DW values

To highlight the importance of analysing DW in the context of BW, we also performed Tukey’s HSD test to compare mean DW among the three genotype groups. As shown in Fig. 18c-d, no differences are significant by this approach. Thus, using linear models with a BW covariate enables much richer analysis of the data, revealing phenotypes that would otherwise go unnoticed.

#### 3.3.3. Cell level

We study genotype effects on cell area adjusting for DW, as well as cell area correlation with DW, in both gonadal and subcutaneous depots. To provide a more comprehensive view of the cell population, we use the three cell area quartiles q={Q1, Q2, Q3} from each slide.

##### 3.3.3.1. Linear models of cell area quartiles as a function of DW and genotype

We segmented 75 inguinal subcutaneous and 72 gonadal whole histology slides with DeepCytometer (with the Corrected method), corresponding to 73 females and 74 males, to produce 2,560,067 subcutaneous and 2,467,686 gonadal cells (on average, 34,134 and 34,273 cells per slide, respectively). (We then removed 2 gonadal and 1 subcutaneous slides due to lack of BW and DW records of two mice.)

We calculated the polygonal area of each white adipocyte contour returned by the segmentation, and pooled the cells in each slide to compute Harrell-Davis (HD) area quartiles, area_q_, q={Q1, Q2, Q3}. We performed an exploratory analysis analogous to the depot level one. Here, we fitted OLS models (area_q_ ~ DW * genotype), genotype={Control, MAT WT, FKO}, stratified by sex, depot and quartiles (plots in Fig. 19a-b, intercept and slope values in Table 6). We used LRTs to compare the OLS models of the three genotypes to each other (Fig. 19c), and t-tests of the slopes to evaluate correlation between area_q_ and DW (p-values from the 36 slope t-tests corresponding to 2 sexes, 2 depots, 3 genotypes and 3 quartiles, before and after Benjamini-Krieger-Yekutieli correction are shown in Table 6).

**Fig. 19.**
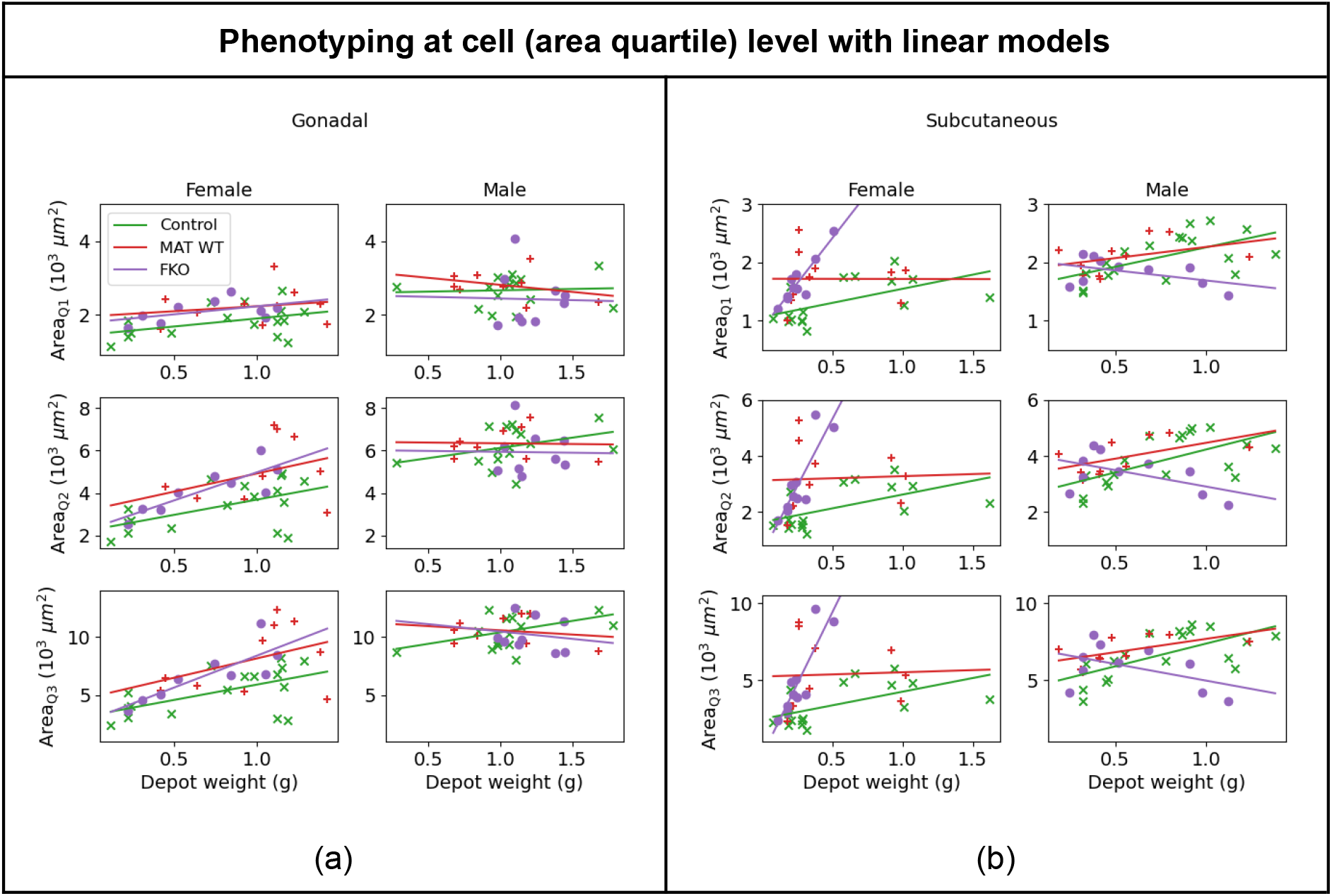
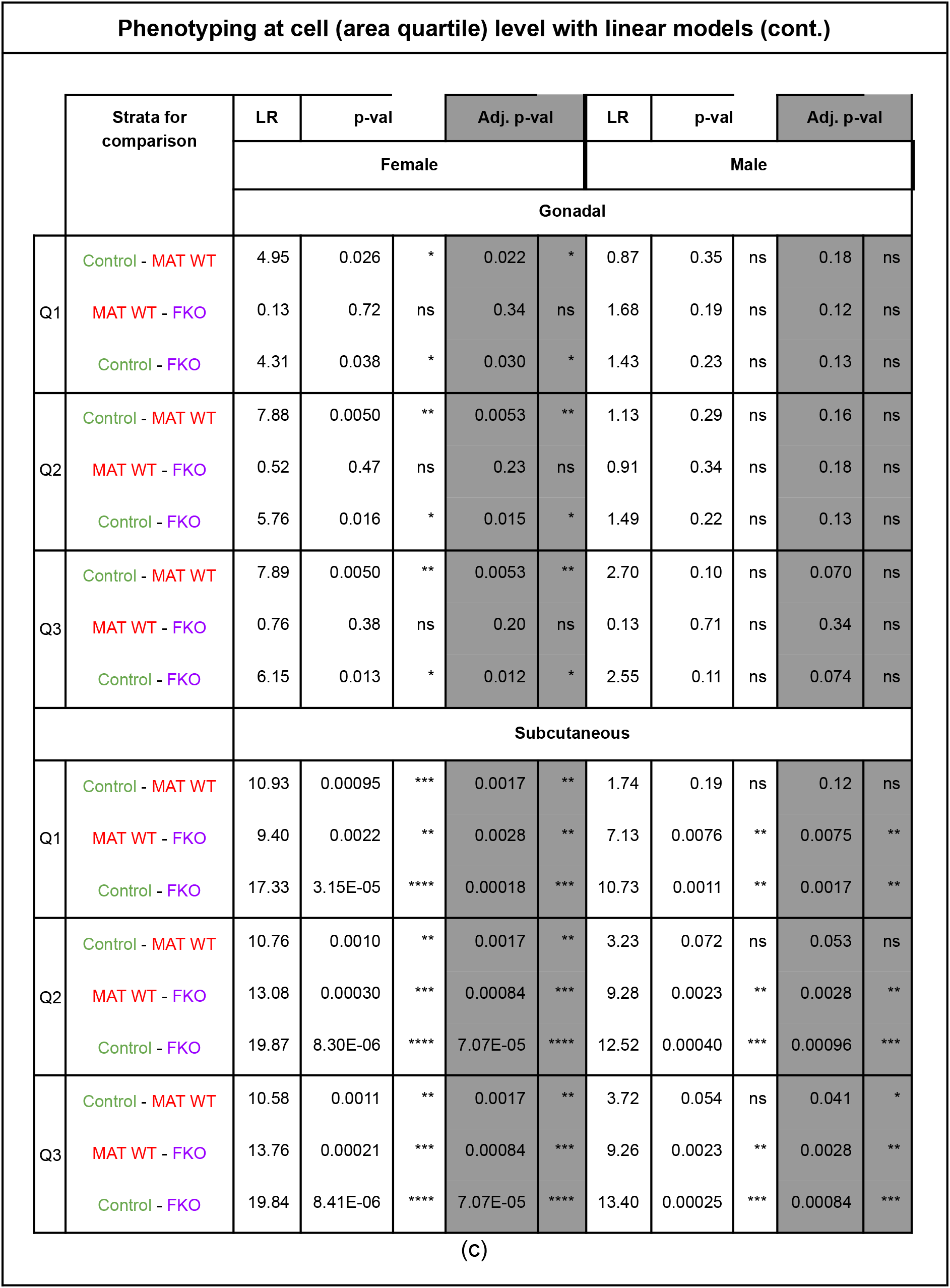
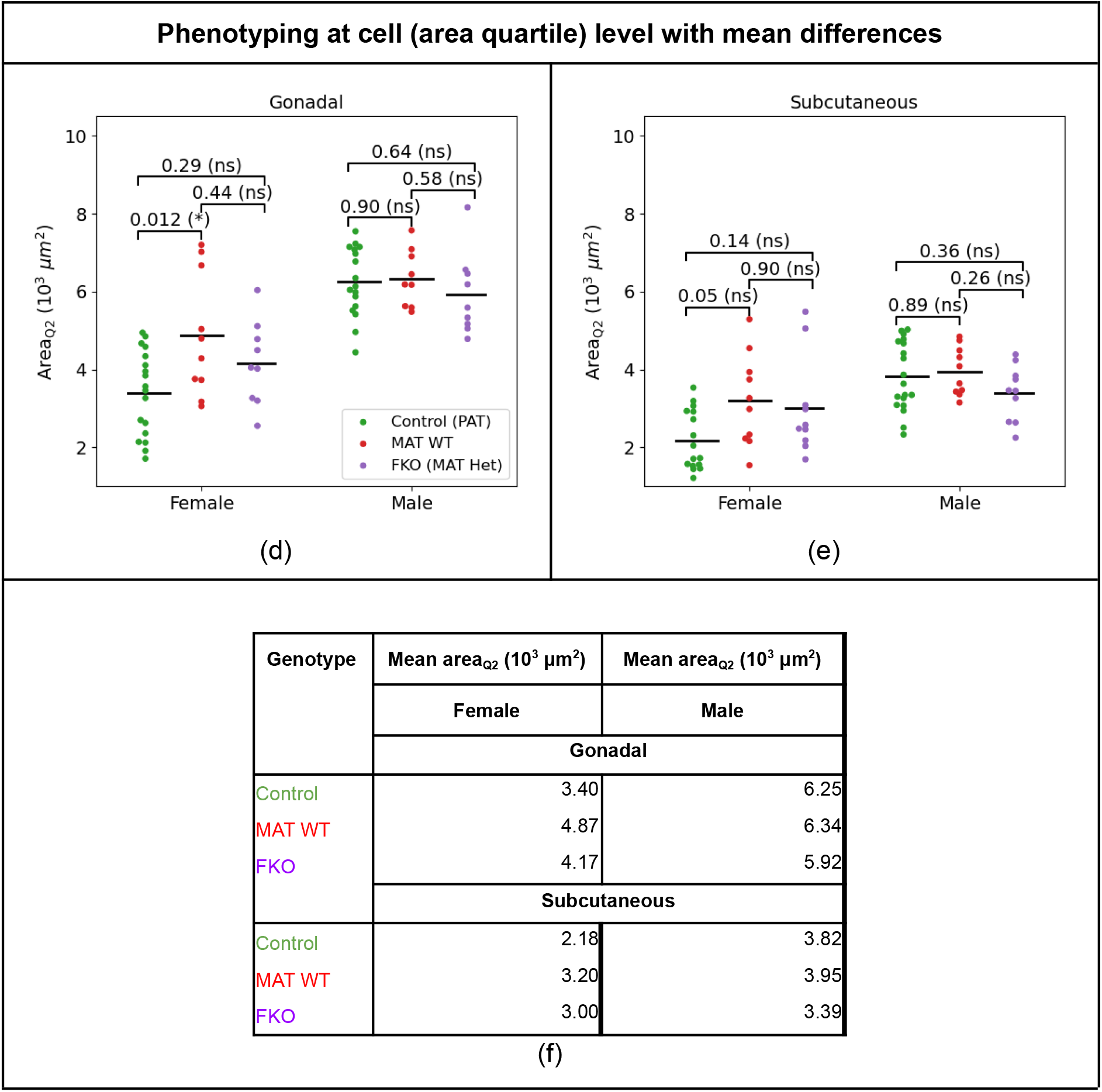
Cell area quartile analysis. (a)-(b) OLS models (area_q_ ~ DW* genotype) for quartiles q={Q1, Q2, Q3}, stratified by sex, depot and genotype, where genotype={Control, MAT WT, FKO (MAT Het)}. Each point corresponds to a mouse. Cell area quartile analysis. (c) P-values of LRT comparisons between Control, MAT WT, FKO linear models, without and with Benjamini-Krieger-Yekutieli multitesting correction. Cell area quartile analysis. (d)-(e) Median (Q2) area swarm plots stratified by sex and genotype. Mean area_Q2_ for each group provided as horizontal black line. P-values of mean differences computed with Tukey’s HSD test for each sex. (f) Mean area_Q2_ values stratified by genotype, sex and depot.

**Table 6.**
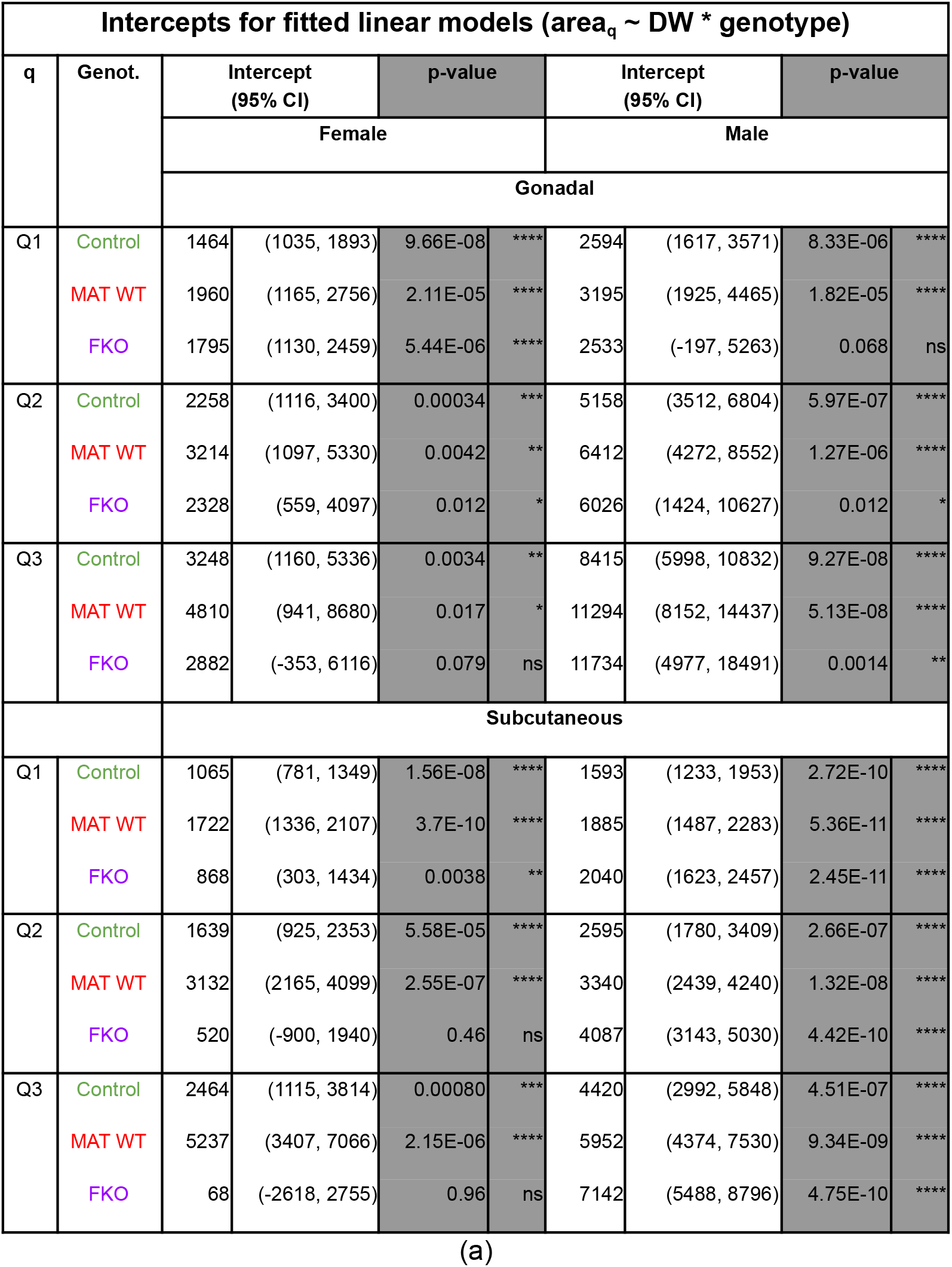
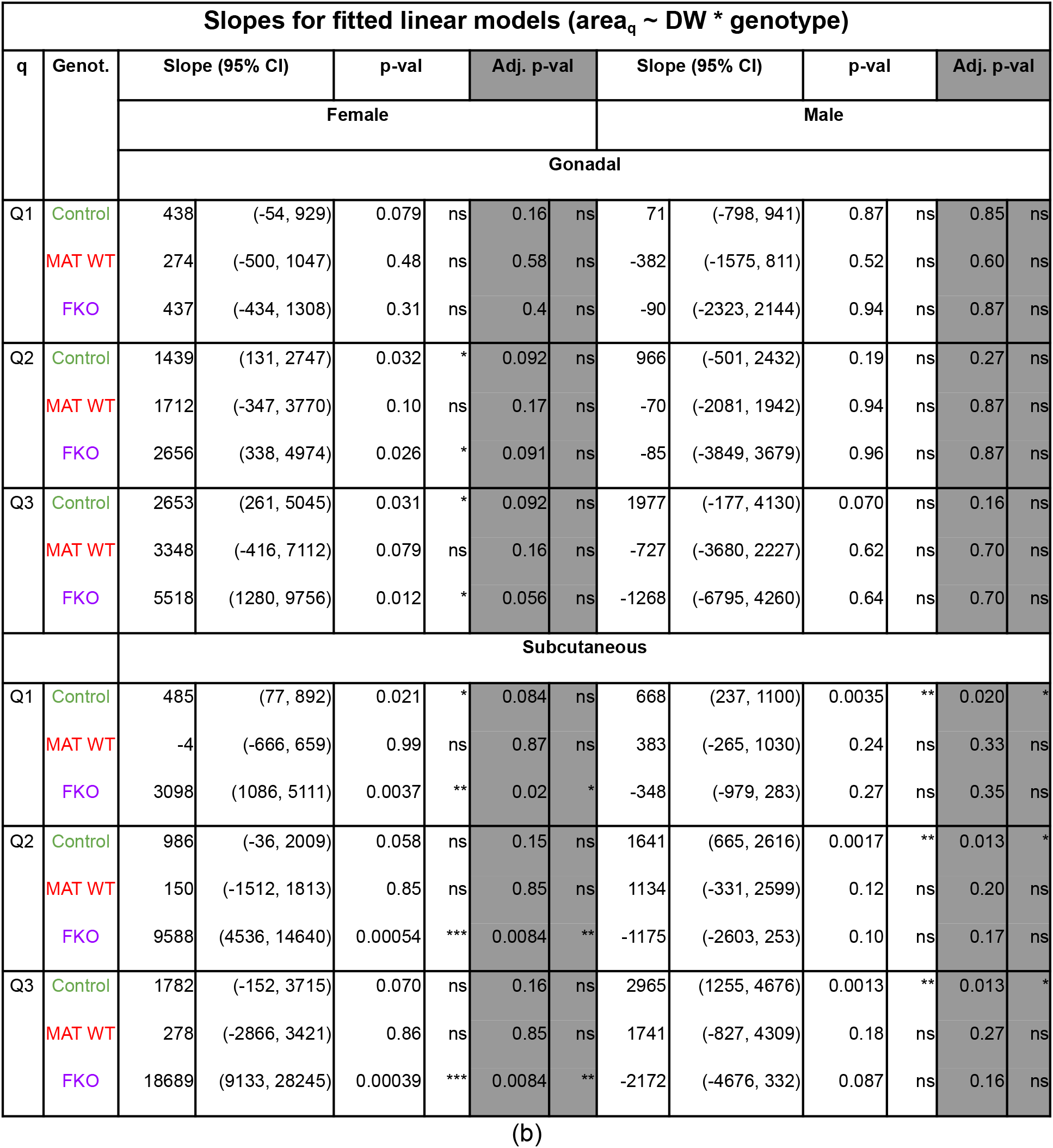
Intercepts and slopes for cell area quartile OLS models (area_q_ ~ DW* genotype) for quartiles q={Q1, Q2, Q3}, stratified by sex, depot and genotype={Control, MAT WT, FKO (MAT Het)}. Plots of the models are shown in Fig. 19. (a) Intercepts. Intercepts and slopes for cell area quartile OLS models (area_q_ ~ DW * genotype) for quartiles q={Q1, Q2, Q3}, stratified by sex, depot and genotype={Control, MAT WT, FKO (MAT Het)}. Plots of the models are shown in Fig. 19. (b) Slopes.

We use the Control group to obtain a baseline of normal physiology. First, Control male white adipocytes are larger than female ones for the same DW, across depots and quartiles. Second, area_q_ increases with DW, although with differences between sexes and depots: the slopes are steeper for females than males in gonadal depots, and vice versa in subcutaneous ones. This is also shown by significant slope p-values for male subcutaneous depots after multitesting correction (and for female gonadal Q2 and Q3 ones before correction).

Next, we explore phenotypes in the MAT WT and FKO groups. In female gonadal depots, differences between MAT WT and FKO are n.s. (according to LRT p-values), but both are significantly different from Controls. In particular, female gonadal MAT WT and FKO cells are larger for any given DW compared to Controls. For instance, plugging the mean female gonadal DW, 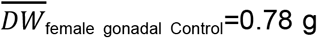 (Fig. 18e), into the Control and FKO area_Q2_ linear models (Table 6), we obtain a predicted area_Q2 female gonadal Control_=3,380 μm^2^ vs. area_Q2 female gonadal FKO_=4,400 μm^2^ (Δ=+30.1%). Male gonadal depots behave differently than female ones, as there are no significant differences between the three genotypes (LRT p-values are n.s.). All male gonadal slopes are n.s. too, before and after multitest correction. This suggests that male gonadal adipocytes maintain roughly a constant size across quartiles regardless of DW or genotype.

By contrast, subcutaneous depots present very strong phenotypes. In female subcutaneous depots, linear models from the three genotypes are very significantly different between them, according to the LRTs: controls feature a positive correlation between DW and cell area; MAT WT slopes are non significantly different from 0, and thus, adipocyte area remains roughly constant regardless of depot size; and FKO models present a very significantly steep slope, suggesting that small increases in depot size correspond to large increases in cell area. However, it should also be noted that the range of FKO subcutaneous DWs is very restricted, up to approximately 0.5 g, so this could be a small sample artifact. Male subcutaneous depots behave differently to female ones. Male subcutaneous Control linear models are similar to MAT WTs (LRT p-values are n.s.), but are very different from FKO models (LRT p_Q1_=0.0017, p_Q2_=0.00096, p_Q3_=0.00084). Consistently with those comparisons, male subcutaneous MAT WT and FKO models are also very different from each other (LRT p_Q1_=0.0075, p_Q2_=p_Q3_=0.0028). Visual inspection of Fig. 19b suggests that male subcutaneous adipocytes have a similar area for DW < 0.5 g in the three genotype groups. However, adipocyte area increases with DW for Controls (positive slopes p_Q1_=0.020, p_Q2_=p_Q3_=0.013) and MAT WTs (positive slopes, p-values n.s.), but decreases or remains constant for FKOs (negative slopes, p-values n.s.). This would suggest a male subcutaneous FKO phenotype where DW increases are due to cell count rather than cell size.

##### 3.3.3.2. Tukey’s HSD test comparison of mean area_Q2_ values

As above, to highlight the importance of analysing cell area in the context of DW, we also performed Tukey’s HSD test to compare mean area_Q2_ among the three genotype groups. As shown in Fig. 19a-b, no differences are significant by this approach, except for area_Q2_ being significantly larger for female gonadal MAT WTs than for Controls (this phenotype was also found by the linear model approach).

#### 3.3.4. Summary of phenotypic findings

In this section we summarise all phenotypic findings, both for differences between linear models that correspond to different genotypes (Table 7), and for slopes (correlation) significantly different from zero (Table 8).

**Table 7.**
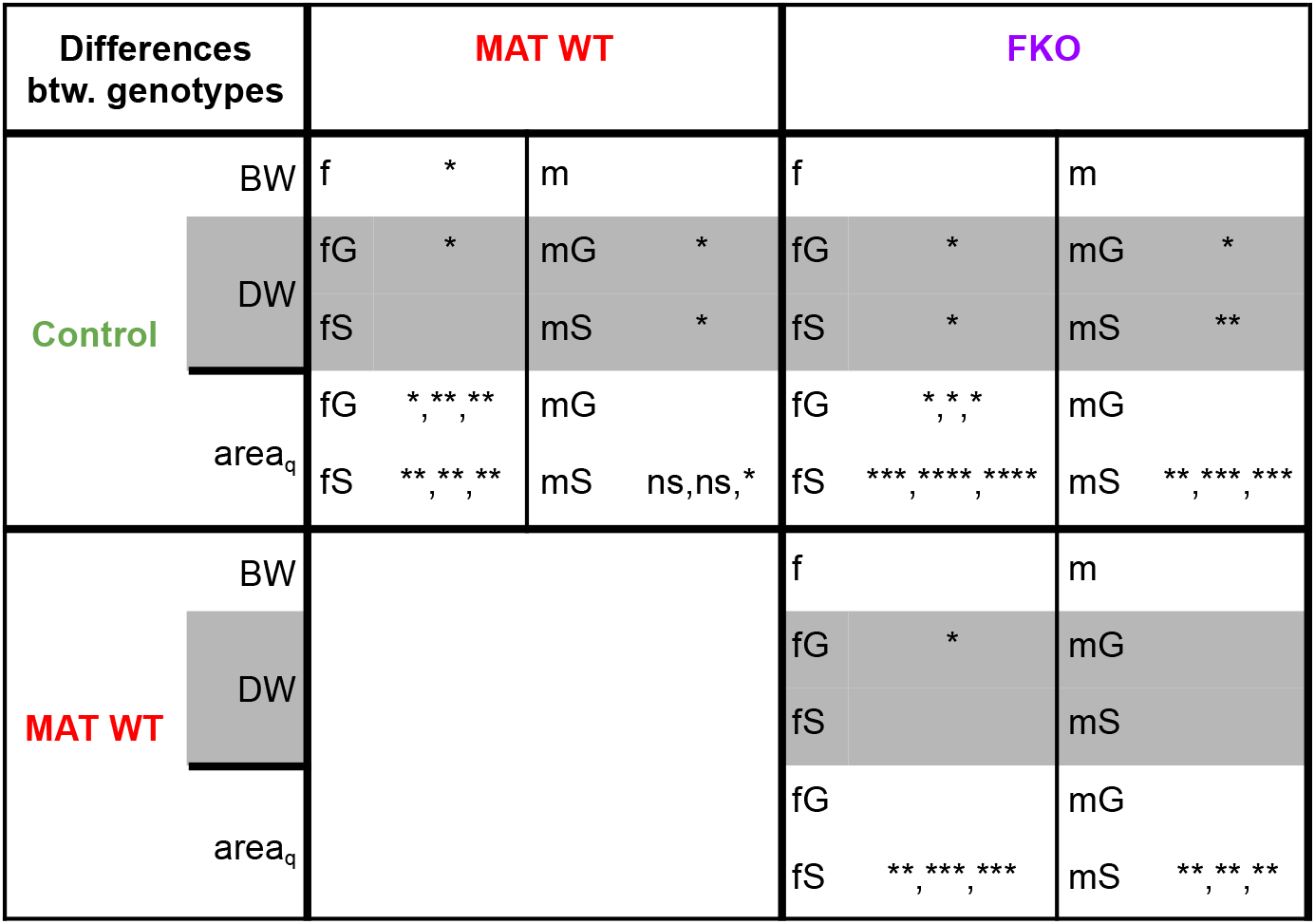
Summary of differences between genotype strata (Control, MAT WT and FKO). f/m/G/S: female/male/gonadal/subcutaneous. Empty cells correspond to n.s. p-values; otherwise, asterisks represent the adjusted p-value. Three p-values are provided for area_q_, corresponding to quartiles={Q1, Q2, Q3}. Significance means that two genotype strata are statistically significantly different under the corresponding model: Tukey HSD’s tests for BW and LRTs for DW, area_q_.

**Table 8.**
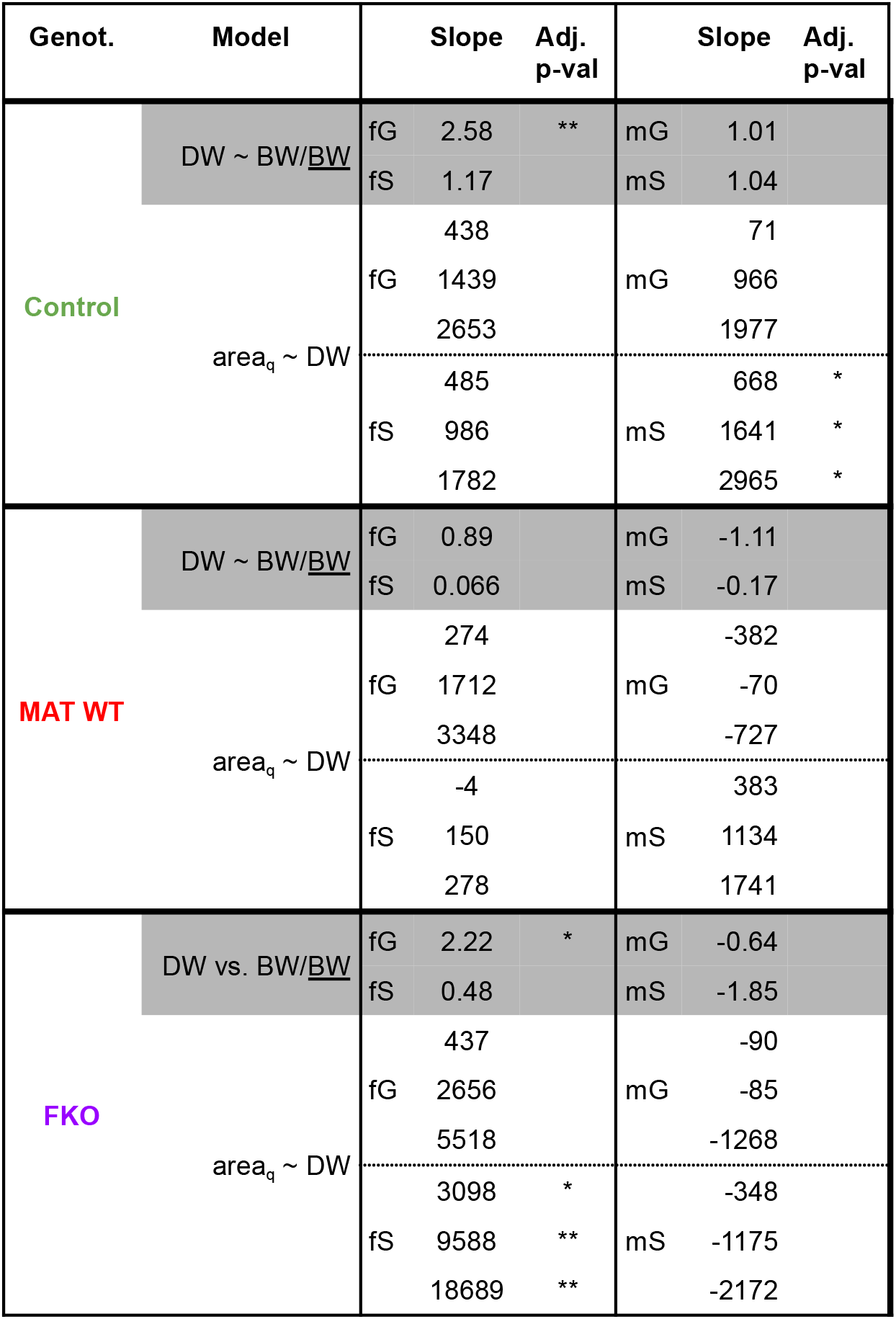
Summary of slopes (and significant correlations) for DW and area_q_ models of genotype strata (Control, MAT WT and FKO). f/m/G/S: female/male/gonadal/subcutaneous. Empty cells correspond to n.s. p-values; otherwise, asterisks represent the adjusted p-value. Three p-values are provided for area_q_, corresponding to quartiles={Q1, Q2, Q3}. Significance means t-test evidence that the slope is ≠ 0 and, equivalently, that Pearson’s correlation coefficient ≠ 0.

## 4. Discussion

This paper presents two parts: first, we proposed DeepCytometer, a pipeline to segment white adipocytes from high resolution H&E histology, together with visualisation tools for the results. Second, we introduced a novel phenotype framework for white adipose tissue that we applied to the *Klf14* mouse data. Unlike previous methods that have been limited to processing small windows containing mostly white adipocytes with good image quality, DeepCytometer can process whole slides containing cell overlaps, other types of tissue, image artifacts and variations in image quality. In the first part of the paper, we addressed several problems that naturally arise in whole slide segmentation: **1) Coarse tissue mask**. By applying traditional image processing techniques to segment tissue areas on the 16× downsampled slide. This resolution was enough to obtain a tissue mask, avoided processing large empty background areas and its computation time was negligible compared to segmentation. **2) Adaptive tiling for whole slide processing**. We improved on previous tiling approaches by proposing a method that does not discard cells cropped by tile edges, chooses each tile’s location and size to reduce overlap and redundant computations. We found a 16.59% reduction in the number of processed pixels compared to uniform tiling. As processing time is linear with the number of pixels, this resulted in an overall speedup of whole slide segmentation.

**3) White adipocyte segmentation**. We built on previous cell segmentation work, with a combination of FCNs and post-processing methods, in particular watershed and mathematical morphology (*Auto* method). As in previous work, we estimated an EDT from adipocyte membranes with a CNN, but followed it by a Contour CNN to find the EDT troughs, because we found that in practice, previously proposed simple post-processing methods did not work consistently in whole slides. In addition, we proposed the Correction CNN to account for cell overlap (*Corrected* method). Our median Dice coefficients were 0.89 (Auto) and 0.91 (Corrected), showing good agreement with the hand traced segmentation. The median relative area errors were −10.19% (Auto) and 4.19% (Corrected). Following previous practice in the literature, these measures would validate our segmentation method.

However, in this paper we also proposed looking at segmentation errors over the population distribution. We found that the segmentation area error was roughly proportional to cell area for cells ≥ 780 μm^2^, which is desirable, but shifts towards more positive values for smaller cells, the bottom 15.9% of the population. This illustrated how segmentation errors in subpopulations could go undetected using summary statistics, and highlighted the need for validating segmentation errors as a function of cell size. This did not affect our phenotyping evaluation, as we used the cell area population quartiles (25%, 50% and 75%) and not the smallest cells. Furthermore, DeepCytometer could successfully segment 347 whole slides of human WAT from the GTEx data set. Running time of the pipeline increased linearly with tissue area, with 5.5 h for a median size slide with 174.7 mm^2^ (848.2 Mpixel) of tissue. Most of our slides contained two slices, so the median time to analyse a slice would be 2.75 h. Of the processing time, 43.9% corresponded to the Auto method, and 56.1% to the Corrected method. Thus, the Corrected method increases computation time by ×2.28 over using only the Auto method. Whether this trade off is acceptable depends on the available resources. In our quantitative experiments, we used Corrected results. **4) Tissue and object classifier**. To determine which segmented objects are white adipocytes, first we classify each histology pixel, and then accept objects that contain ≥ 50% white adipocyte pixels. We weighed the object classifier’s ROC by the number of pixels per object, to balance the contribution to classification errors of white adipocytes (small but numerous) with other objects (scarcer but large). The 0.97 area under the ROC indicated overall good performance of the classifier. With the 50% acceptance threshold, the false positive rate (FPR) = 15% and true positive rate (TPR) = 95%, which we considered acceptable for phenotyping. If a lower FPR was required in other work, the 50% threshold could be raised. Given the large cell counts in the slides, the resulting lower TPR may be an acceptable trade-off. **5) Ground truth / training data**. We created a ground truth / training data set for the EDT, Contour and Correction CNNs with 55 random windows from 20 mice, totalling 2,117 white adipocyte hand traced contours, for 10-fold cross validation. For the Tissue CNN, we added another 71 windows containing only non-white adipocyte regions. This was necessary to create a balanced set of ≈ 23.7 · 10^6^ white adipocyte pixels and ≈ 45.1 · 10^6^ non white adipocyte pixels. We hope that the data set (Casero et al., 2021a) will be a useful resource in the field. **6) Segmentation results visualisation**. Integration of our pipeline with AIDA allowed us to review whole slide segmentation results in real-time. AIDA was launched on the same GPU server as the pipeline, and enabled real-time review of the results from a desktop or laptop using a regular browser. The pipeline features saving each block as a separate layer or all in the same layer for display. AIDA also allows manual correction of labels. **7) Cell size heterogeneity visualisation**. To highlight spatial size heterogeneity, we proposed a colour scale proportional to the cell area’s quantile. This scheme produced highly contrasted images readily showing cell area heterogeneity across tissue samples, with subpopulations of different sizes grouped in clusters (Section S1). **8) Suitability of hand traced data set for cell population study**. Section 3.2.2 showed very significant differences between the distributions obtained from the hand traced data set and whole slide automatic segmentations used as ground truth. This suggests that even though the hand traced data set contained 1,903 cells from 60 random windows, it failed to represent the true distribution of white adipocyte areas. A discrepancy between cell size estimates from sampled windows and whole slides was also observed by (Maguire et al., 2020). Although spatial analysis is without the scope of this work, we conjecture that the intra-slice clustering observed in

Section S1 introduces strong spatial local correlations in cell size, thus reducing the effective size of the training data set. Such difference highlights the need for whole slide segmentation and analysis, with pipelines that can run on dozens or hundreds of slides.

In the second part of this paper, we performed an exploratory study of *Klf14* phenotypes in B6NTac mice at three nested levels-animal (body weight, BW), depot (depot weight, DW) and cell (cell area quartile)-in three strata (referred to as “genotype” for simplicity): Control (PAT WT and PAT Het), MAT WT and FKO (Mat Het). Cell areas were obtained by DeepCytometer from 75 inguinal subcutaneous and 72 gonadal full histology slides. At the DWand cell levels, we considered two phenotype assessments: 1) association between variables (e.g. BW vs DW) via t-tests of linear model slopes and 2) differences between genotypes using LRTs. In addition, at the cell level we computed pdfs of cell areas.

Control mice presented marked BW sexual dimorphism, with males weighing 52.8% more than females on average. DW increased with BW in the four Control sex/depot strata, although the correlation was only significant for female gonadal depots (for male gonadal ones, it is significant only before multitesting correction). It is unclear whether the lack of correlation in the other strata is physiological, or an artifact of the limited number of data points. Cell area tends to increase with DW in the four Control strata, but in this case the correlation is only significant for male subcutaneous depots (female gonadal and subcutaneous correlation is significant before multitesting correction).

The MAT WT and FKO groups present differences from Controls and between them. Female MAT WTs have significantly larger mean BW than Controls (22.9%), but both gonadal and subcutaneous DW/BW slopes are smaller than in Controls. Female MAT WT gonadal cell area increases with DW at a similar rate (slope) as Control’s, but for any DW value, they are ~30% larger. Meanwhile, female MAT WT subcutaneous cell area is uncorrelated with DW; its values remain relatively constant, comparable to the largest Control cell area values. In summary, female MAT WTs are on average larger than Controls, and the larger they are, the lower fat percentage they have compared to similarly sized Controls. As they also tend to have larger adipocytes than Controls, this suggests lower adipocyte count per depot.

Male MAT WTs have similar mean BW as Controls. However, their depot physiology appears different, as MAT WT DW/BW slopes are negative, whereas Control ones are positive. Because the MAT WT slope p-values are n.s., this could be a chance observation, but as with females, the trend is that larger male MAT WTs have lower fat percentage than their Control counterparts. Male MAT WT cell area models, however, are not significantly different from Control ones. To sum up, male MAT WTs are similarly sized as Controls, the larger they are, the lower fat percentage they have (although this could just be a chance observation), and differences in DW seem due only to cell counts.

Female FKO mean BW is slightly larger than Controls (p-value n.s.). Their DW increases at a lower rate than Controls’ for both depots (slopes are 14.0% smaller for gonadal depots and 59.0% smaller for subcutaneous depots), but more noticeably, their DW values are consistently smaller for any BW (−34.2% gonadal and −61.8% subcutaneous for 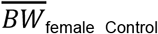). Female FKO gonadal cell area models (like MAT WT ones), have similar slopes but cell areas are ~30% larger than Controls’. Female FKO subcutaneous cell models, by contrast, are very different from both Controls’ and MAT Ws’, with very steep slopes, suggesting a fast increase of cell area with DW. However, due to the very limited range of DW values, this result could be a small data set artifact (the cells measured are within the interval of cell areas for MAT WTs, so perhaps larger FKO subcutaneous depots would not present adipocytes as large as those predicted by the models). Altogether, female FKOs are barely larger mice than Controls, but they have noticeably lower fat percentage, especially regarding subcutaneous depots. As they also tend to have larger adipocytes than Controls, this suggests lower adipocyte count per depot (same behaviour as MAT WTs).

Male FKOs are also slightly larger than Controls (n.s.), but their DW *decreases* with increasing BW, both for gonadal and subcutaneous depots. However, because of the intercepts in the models, their gonadal and subcutaneous DWs are within a similar range as Controls’ (smaller FKOs may have higher fat percentage, and larger FKOs, lower). Male FKO gonadal cell area is similar to Controls’ for any DW. Male FKO subcutaneous cell areas are similar to Controls’ for DW < 0.5 g, but then decrease or remain constant instead of growing with increasing DW. In summary, male FKOs are barely larger mice than Controls; their fat percentage decreases with increasing BW, but is on average similar to Controls’ fat percentage. There is a biological difference in cell areas, as in Controls, it increases with DW, but in FKOs it decreases or remains constant, suggesting a phenotype where DW increases are due to cell count rather than cell size. We also found that more traditional analysis, e.g. by Tukey HSD test of mean differences, most of the linear model phenotypes went undetected. This highlights the need to consider BW and DW as confounders in the analysis.

Putting all the phenotype analysis results together, our exploratory analysis reveals interesting leads by breaking down the study into animal, depot and cell level, and assessing genotype effects. The DW vs. cell area relationships could be directly connected to hypertrophy vs. hyperplasia. However, further investigation with more mice is necessary to be able to provide explanatory mechanisms for the phenotypes.

In future work, the pipeline can be redesigned with alternative deep learning approaches that find candidate objects and then segment them, as opposed to DeepCytometer, which first segments all objects and then classifies them, e.g. based on Faster R-CNN (Ren et al., 2017), Mask R-CNN (He et al., 2018), TensorMask (Chen et al., 2019) or XCiT (El-Nouby et al., 2021). Although we have proved that DeepCytometer can analyse hundreds of slides with standard resources, we would like to improve its architecture to scale it to studies with thousands of slides, without needing large cloud computing resources. Furthermore, the cell area heterogeneity we found in the heatmaps begs the question of whether a single whole slide provides an appropriate representation of a whole depot’s cell population. This could be decided by a study comparing depot cell populations from a single slide and from multiple slides, or by extending this work to segmentation of 3D image modalities. The latter approach is substantially harder, as it is necessary to use 3D imaging (usually with poorer definition than H&E histology), redesign the pipeline for 3D data (e.g. extending 3DeeCellTracker (Wen et al., 2021) to adipocytes and three orders of magnitude more cells), and creating a training data set in 3D is more challenging than in 2D (Dunn et al., 2019; Tasnadi et al., 2020). That notwithstanding, a 3D approach would allow the analysis of cell volumes instead of areas of intersections of cells with the cutting plane, as well as cell counts per depot. Moreover, studies derived from 2D images and depot weight such as ours can observe correlation between cell area and depot weight, but not determine whether all the depot weight gain is due to cell size increase or a net increase in combined cell volume + cell count. This limitation would be solved by full segmentation of 3D depot images.

## Acknowledgements

We would like to thank George Nicholson (Oxford Statistical Department) and Laura Bramley (BDI, University of Oxford) for their advice on Statistical methods; Stefano Malacrino (Dept. Engineering Science, University of Oxford), for his help with AIDA; and Liz Bentley, for her advice in the project.

Mouse Klf14^tm1(KOMP)Vlcg^ adipose samples, body and depot weight data were obtained from a previous study that was supported by UK Medical Research Council grant MR/J010642/1. RDC is supported by UK Medical Research Council funding MC_U142661184. Some of this work was supported by core funding from the UK Medical Research Council.

Computation used the Oxford Biomedical Research Computing (BMRC) facility, a joint development between the Wellcome Centre for Human Genetics and the Big Data Institute supported by Health Data Research UK and the NIHR Oxford Biomedical Research Centre. The views expressed are those of the author(s) and not necessarily those of the NHS, the NIHR or the Department of Health.

## Appendix A. Supplementary materials

Download 20190727_cytometer_paper_supplementary_materials.pdf. Supplementary Data S1. White adipocyte area heatmaps of all the whole slides automatically segmented by DeepCytometer.

## Appendix B. DeepCytometer technical details

### B.1. Relation between pixel-classification and EDT regression

We note that the pixel-classification (Van Valen et al., 2016) and EDT (Casero et al., 2019; Cientanni and Casero, 2018; Wang et al., 2019) approaches reviewed in the *Introduction* are related, as pixel-classification can be seen as a simplification of signed EDT regression by applying the sign function

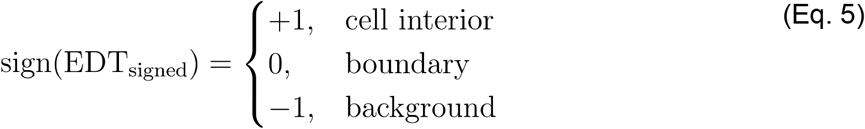

Thus, the EDT representation is richer, because it not only estimates whether a pixel is inside a cell, but how far from the membrane it is. This could account for the better segmentation results found by (Wang et al., 2019).

### B.2. Effective receptive field (ERF) of CNNs

The ERF of each CNN and fold was computed and is shown in Table 9. The maximum cell size in our manual data set is just under 20,000 μm^2^. This corresponds to a diameter of 160 μm for a circular cell, or 353 pixels for pixel size 0.454 μm. An ERF=131 pixels covers around 37.1% of the largest cell diameter.

**Table 9.** Effective Receptive Field (ERF) of the DeepCytometer pipeline CNNs. All sizes in pixels.

EDT CNN
| *Fold* | 0 | 1 | 2 | 3 | 4 | 5 | 6 | 7 | 8 | 9 |
| --- | --- | --- | --- | --- | --- | --- | --- | --- | --- | --- |
| *Height* | 128 | 131 | 131 | 131 | 131 | 131 | 131 | 130 | 131 | 124 |
| *Width* | 131 | 131 | 131 | 131 | 129 | 131 | 127 | 131 | 131 | 123 |

Contour CNN
| *Fold* | 0 | 1 | 2 | 3 | 4 | 5 | 6 | 7 | 8 | 9 |
| --- | --- | --- | --- | --- | --- | --- | --- | --- | --- | --- |
| *Height* | 131 | 131 | 131 | 131 | 131 | 131 | 131 | 131 | 131 | 131 |
| *Width* | 131 | 131 | 131 | 131 | 131 | 131 | 131 | 131 | 131 | 131 |

Tissue CNN
| *Fold* | 0 | 1 | 2 | 3 | 4 | 5 | 6 | 7 | 8 | 9 |
| --- | --- | --- | --- | --- | --- | --- | --- | --- | --- | --- |
| *Height* | 123 | 131 | 130 | 131 | 131 | 131 | 131 | 131 | 131 | 131 |
| *Width* | 131 | 131 | 131 | 131 | 131 | 131 | 131 | 131 | 131 | 131 |

| *Fold* | 0 | 1 | 2 | 3 | 4 | 5 | 6 | 7 | 8 | 9 |
| --- | --- | --- | --- | --- | --- | --- | --- | --- | --- | --- |
| *Height* | 131 | 131 | 131 | 131 | 131 | 131 | 131 | 131 | 131 | 131 |
| *Width* | 131 | 131 | 131 | 131 | 131 | 131 | 131 | 131 | 131 | 131 |

### B.3. Pseudocode for adaptive tiling

***adaptive_block_algorithm()*:**

1. Let

a. *histology_filename* be the filename of the full resolution histology slide.
b. *mask_todo_lores:*= coarse tissue mask computed in previous section.
c. *downsample_factor*:= 8
d. *max_window_size*:= [2751, 2751] # due to GPU memory limit
e. *border*:= [65, 65] # Overlap with other windows to account for receptive field
2. Open file pointer to full resolution histology without reading it into memory *im*:= OpenSlide(*histology_filename*)
3. Loop until *mask_todo_lores* is empty.

a. Compute coordinates of next processing block, both in full resolution and downsampled image coordinates [*box_coords_hires*, *box_coords_lores*]:= get_next_roi_to_process(*mask_todo_lores*, *downsample_factor, max_window_size*, *border*)
b. Load block to process from full resolution image *im_box*:= OpenSlide.read_region(*im*, *box_coords_hires*)
c. Extract low resolution mask for the block and upsample to full resolution *mask_box_lores*:= *mask_todo_lores[box_coords_lores*] *mask_box_hires*:= *resize(mask_box_lores, downsample_factor*, ‘nearest neighbour’)
d. Segment histology to obtain one label per cell, and mask of cells on the edge *labels_hires*, *mask_edge_hires*:= DeepCytometer_pipeline(*im_box*, *CNN_models, segmentation_parameters*)
e. If no cells found, wipe out current box from coarse tissue mask to avoid infinite loops *mask_todo_lores*[*box_coords_lores*]:= 0 go to next iteration in 3.
f. Downsample mask of edge objects so that the coarse tissue mask can be updated *mask_edge_lores*:= resize(*mask_edge_hires*, size(*box_coords_lores*), ‘nearest neighbour’)
g. Convert labels to contours for AIDA display
h. Update current block of coarse tissue mask with edge mask *mask_todo_lores[mask_box_lores]*:= *mask_edge_lores*

Function get_next_roi_to_process() convolves the coarse tissue mask with a vertical and horizontal line kernel and combines the outputs to find the location of the next processing block.

***get_next_roi_to_process()*:**

1. Compute convolution kernel size *L*:= *int((max_window_size* - 2 *border)* / *downsample_factor)*
2. Let *kh* and *kv* be convolution kernels with size *L*×*L* pixels. The kernels are all zeros except for a horizontal or vertical line of ones through the middle, respectively (Fig. 2c).
3. Compute Fast Fourier Transform (FFT) convolution of coarse tissue mask, with output cropped to mask size *zh*:= *mask_todo_lores 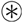 kh* *zv*:= *mask_todo_lores 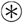 kv*
4. Compute hits where the processing block would both have a mask pixel on the top and left edges as the Hadamard or pointwise product (Fig. 2d) *hits_lores*:= *zh ∘ zv*
5. Choose first found hit *(x0, y0)* in *hits_lores* as top-left corner of block (Fig. 2e).
6. Choose bottom-right corner of block *xend*:= *x0* + *max_window_size* - 2 *border* *yend*:= *y0* + *max_window_size* - 2 *border*
7. Reduce block size if there are empty mask rows/columns at the bottom/right *xend*:= min(*xend*, last column in block with mask pixel) *yend*:= min(*yend*, last row in block with mask pixel)
8. Add a border around the block to account for the effective receptive field. Crop the border if it overflows the image edges, where the downsampled image has size (Rd, Cd) pixels *x0*:= max(*x0* - *border*, 0) *y0*:= max(*y0* - *border*, 0) *xend*:= min(*xend* + *border*, Cd-1) *yend*:= min(*yend* + *border*, Rd-1)
9. Upsample block coordinates for full resolution image *x0_hires*:= round(*x0* * *downsample_factor)* *y0_hires*:= round(*y0* * *downsample_factor)* *xend_hires*:= round(*xend* * *downsample_factor)* *yend_hires*:= round(*yend* * *downsample_factor*)

### B.4. Deep CNNs training methodology

#### B.4.1. Training for 10-fold cross validation

All CNNs were trained with the same 10-fold cross validation described in Table 1: 20 mice randomly partitioned in 10 sets of 18 mice for training and 2 for validation.

#### B.4.2. Training with a combination of labelled and unlabelled pixels

As advanced in the Introduction, it is convenient to use training images with a combination of labelled (‘White adipocyte’, ‘Background’, ‘Other tissue’) and unlabelled pixels. Instead of becoming a new class (‘Void’), unlabelled pixels should not contribute to the training process, as if they were not part of the training data set at all. This functionality is not available in Keras 2.2. Thus, we implemented an extension (Casero, 2019) that enables element-wise weighting of pixel-wise scores. Let *z* be the score matrix of size (*R, C*), where each element *z_i, j_*, *i* = 0,…, *R* – 1, *j* = 0,…, *C* – 1 is the contribution of an output pixel to the loss. Let *w* be a weighting matrix such that the loss is

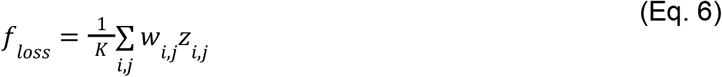

where *K* ≤ *RC* is the number of output pixels where *w_i,j_* ≠ 0. If

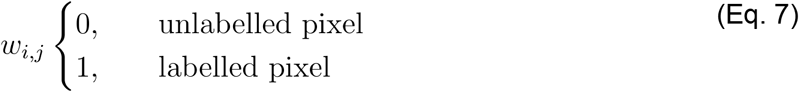

then *f_loss_* is the average score of labelled pixels.

#### B.4.3. EDT CNN and Contour CNN training data set

We used the 55 histology images with 2,117 ground truth white adipocyte (WA) hand traced contours from Table 1. The corresponding SVG files containing the description of the hand traced WA contours (as described in Section 2.2) were read. Each contour was rasterised as a closed polygon. Pixels that belonged to a single polygon were labelled as seeds. Then a watershed algorithm expanded the seeds over areas where polygons overlap. This effectively found a compromise boundary between overlapping cells. Boundaries were computed as pixels between two labels or between label and background. Ground truth EDTs were computed with respect to those boundaries (Fig. 3c). Boundaries were then dilated with a 3×3 kernel because we found that the Contour CNN training was unsatisfactory on 1-pixel thick boundaries. Masks of labelled pixels were computed for the loss function from the union of all polygons, and then were dilated with a 3×3 kernel. The histology windows, EDTs, boundaries and masks data set was then 10× augmented with random rotations up to ±90°, a scaling factor in [0.9, 1.1], and horizontal and vertical flips. The augmented images were split into 4 blocks due to GPU memory restrictions. Blocks where the mask had fewer than 1,900 pixels were discarded.

#### B.4.4. EDT CNN training

We trained the 10-fold CNNs using the augmented histology windows (input), ground truth EDTs (output), masks of labelled pixels (loss function masks), Adadelta optimisation (Zeiler, 2012) with He uniform variance scaling initialisation (He et al., 2015), mean absolute error (MAE) loss, MAE and mean squared error (MSE) metrics for validation with the left-out data, a batch size of 10 and 350 epochs until the loss and metrics converged.

#### B.4.5. Contour CNN training

We trained the 10-fold CNNs using the EDTs estimated by the previous CNN (input), ground truth dilated boundaries (output), masks of labelled pixels (loss function masks), Adadelta optimisation (Zeiler, 2012) with He uniform variance scaling initialisation (He et al., 2015), binary cross entropy loss, accuracy metric for validation with the left-out data, a batch size of 10 and 500 epochs until the loss and metric converged.

#### B.4.6. Tissue CNN training data set

We used all 126 histology images with hand segmentations in Table 1, containing a total of 2,117 “white adipocyte” (WA) objects and 232 “other” (NWA) objects. All WA and NWA contours were read. Each contour was rasterised as a polygon. Pixels within each polygon were labelled as “1” for WA and background, and “0” for other types of tissue to create the classifier ground truth. Pixels in overlap areas between a WA and NWA were considered NWA. The histology windows, classifier ground truth and masks data set were 10× augmented with random rotations up to ±90°, a scaling factor in [0.9, 1.1], horizontal and vertical flips and shear angle in [-15°, 15°] to an output shape of 1,416×1,416 to avoid cropping out training pixels. The augmented images were split into 4 blocks due to GPU memory restrictions.

#### B.4.7. Tissue CNN training

We trained the 10-fold CNNs using the augmented histology windows (input), classifier ground truth (output), masks of labelled pixels (loss function masks), Adadelta optimisation (Zeiler, 2012) with He uniform variance scaling initialisation (He et al., 2015), binary focal loss (Lin et al., 2020) with *γ* = 2, α = 0.4, accuracy metric for validation with the left-out data, a batch size of 8 and 37 epochs until the loss and metric converged. We used a cyclical learning rate (Smith, 2017) with a triangular cycle that scales initial amplitude by half each cycle, initial learning rate 10^-7^, upper boundary 10^-2^ and number of training iterations per half cycle equal to 8× training iterations in epoch.

#### B.4.8. Correction CNN training data set

We used the same 55 histology images, hand traced contours and rasterised polygons as for the EDT and Contour CNNs. Let *a_h_* be the hand traced contour polygon area, and 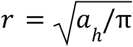 the radius of a circle with the same area. We generated a series of incorrect segmentations by eroding and dilating the ground truth segmentation using an *l_kernel_* × *l_kernel_* square kernel with length *l_kernel_* = ⌊2*r*|*k*| + 1⌋, where the parameter *k* ∈ {± 0.03, ± 0.07, ± 0.10, ± 0.15, ± 0.20}(*k* < 0 for erosion and *k* > 0 for dilation). Thus, we established a correspondence between a histology cropped image, the eroded/dilated segmentation mask *m_k_* and the segmentation error *m_k_* – *m_h_*, where *m_h_* is the hand traced contour polygon. Note that *m_k_* – *m_h_* is −1 for pixels that underestimate the segmentation, 0 for correctly segmented pixels and +1 for pixels that overestimate the segmentation. Next, we multiplied the histology by +1 within *m_k_* and by −1 without. We also computed a mask for the loss function by dilating *m_k_* ∪ *m_h_* by a factor *k*′ = 0.30. Finally, the resulting image, the segmentation error *m_k_* – *m_h_* and the loss function mask were cropped and resized according to the bounding box of *m_k’_* as described above in the Correction CNN architecture section.

#### B.4.9. Correction CNN training

We trained the 10-fold CNNs using the cropped and resized −1/+1 masked histology (input), segmentation error *m_k_* – *m_h_* (output) and loss function mask, the same cyclical learning rate as in the Tissue CNN, Adadelta optimisation (Zeiler, 2012) with He uniform variance scaling initialisation (He et al., 2015), mean squared error (MSE) loss, mean absolute error (MAE) and MSE metrics for validation with the left-out data, a batch size of 12 and 100 epochs until the loss and metrics converged.

#### B.4.10. DeepCytometer running times

We computed running times on a random sample of 95 automatically segmented whole slides. We measured the tissue area as the area covered by the coarse tissue mask. Ordinary Least Squares model (time ~ tissue area) showed a linear relationship with intercept β_0_=297.1 ± 2,495.8 s, slope β_1_=111.9 ± 10.6 s / mm^2^. Tissue areas were between 41.5 - 612.3 mm^2^ (201.5 Mpixel - 2973.2 Mpixel), corresponding to a computation time of 4,939 - 6,8784 s (1.4 - 19.1 h) in the linear model. The area HD quartiles were (Q1, Q2, Q3) = (102.4, 174.7, 276.2) mm^2^ corresponding to (3.3, 5.5, 8.7) h. It should be noted that each slide contained two tissue slices.

Segmentation times were calculated applying the Corrected method to the 60 training images for population studies. The Auto part of the pipeline took 43.9% ± 1.6% of the total time, and overlap correction took the other 56.1% ± 1.6%. Thus, overlap correction increased Auto computation time by a factor ×(2.28 ± 0.08).

