## Supplementary materials for "Phenotyping of *Klf14* mouse white adipose tissue enabled by whole slide segmentation with deep neural networks"

### S1. White adipocyte area heatmaps of all the whole slides automatically segmented by DeepCytometer

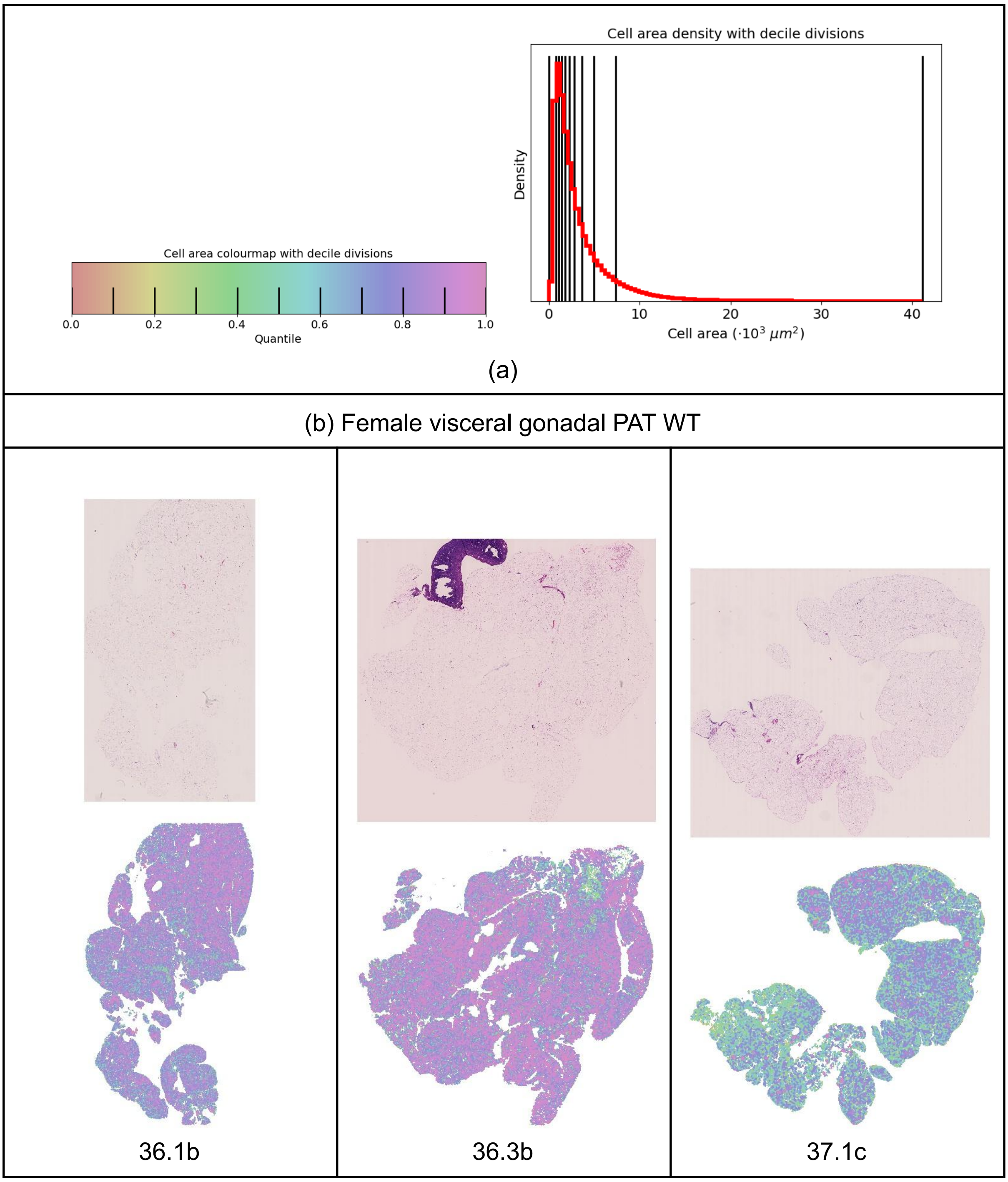

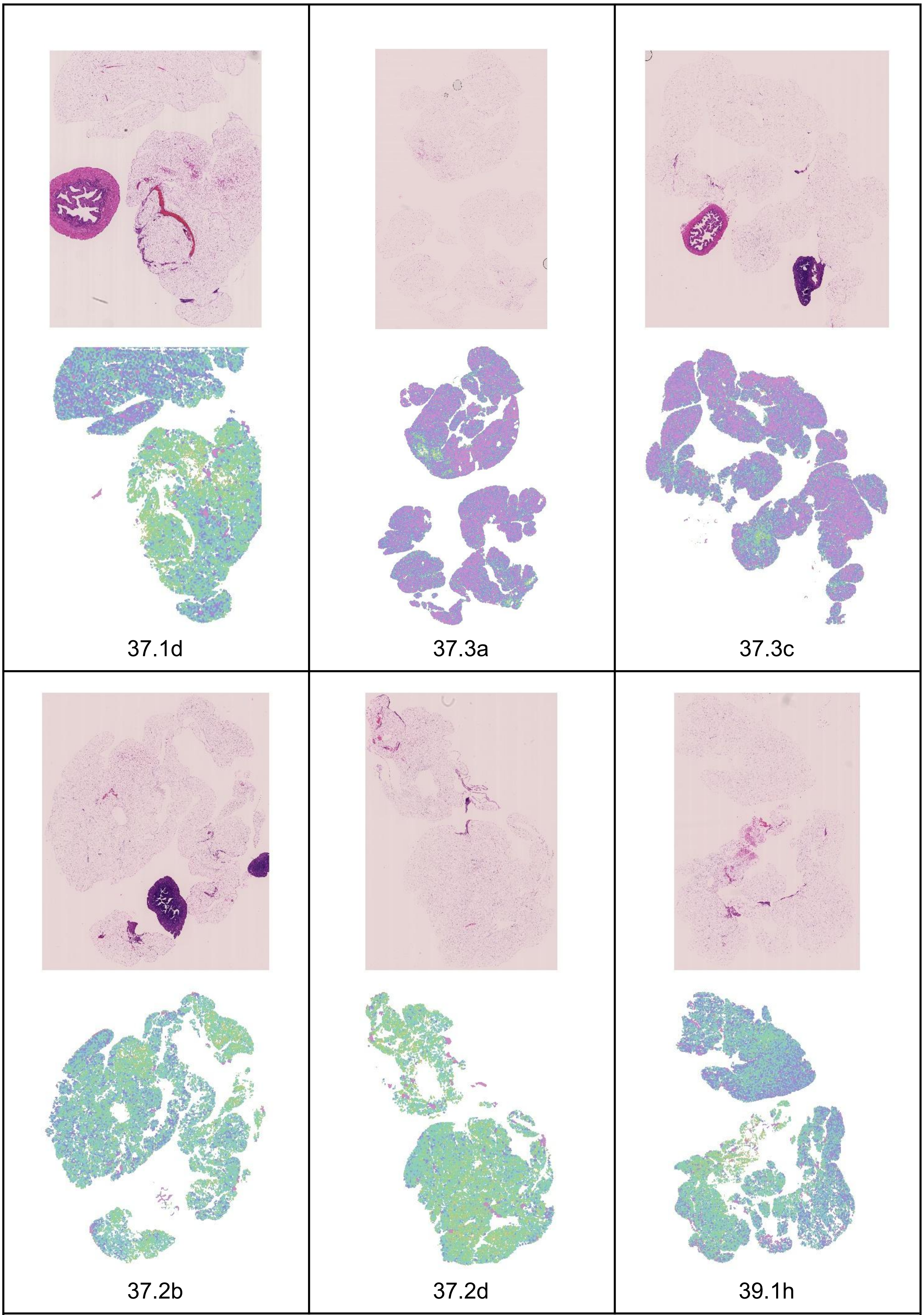


(c) Female visceral gonadal PAT Het

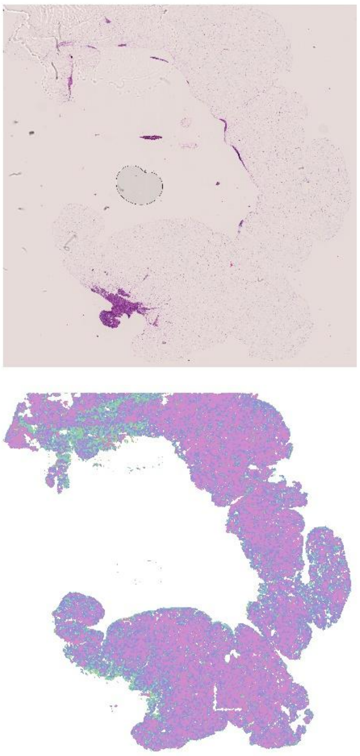

36.1a

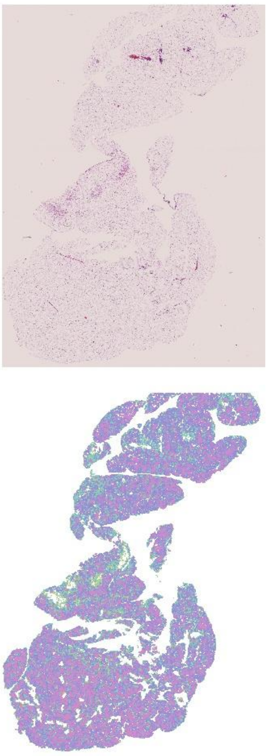

36.1c

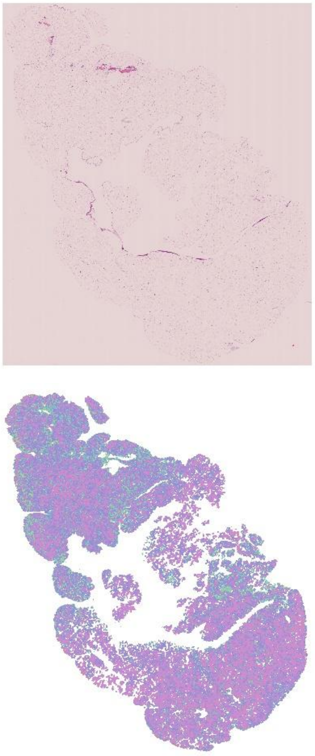

36.3a

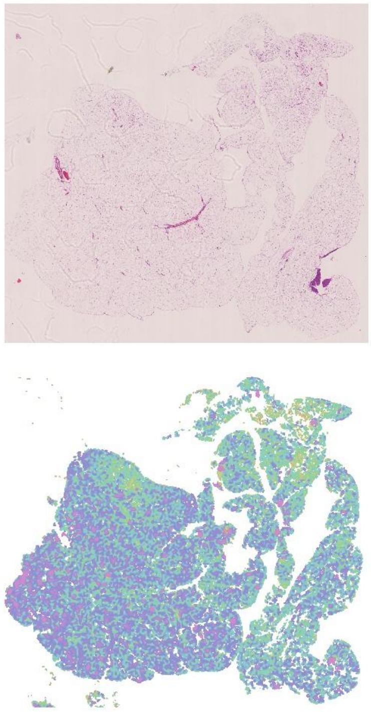

37.1a

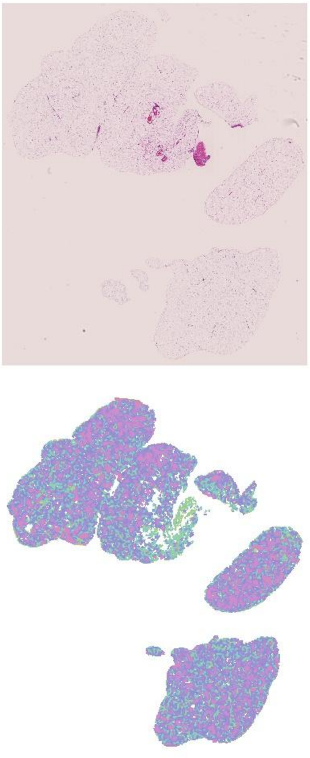

37.1e

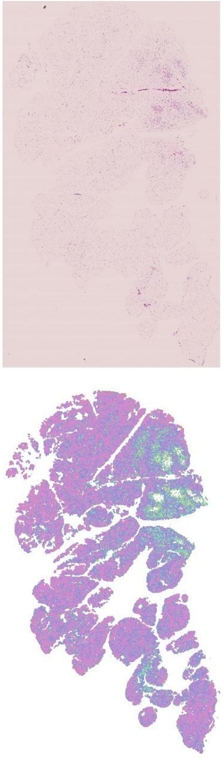

37.2e

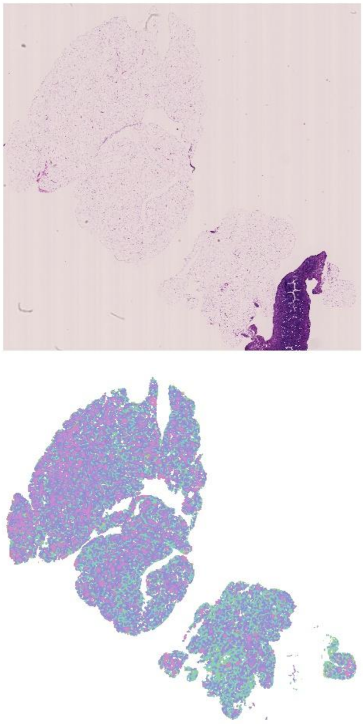

37.1b

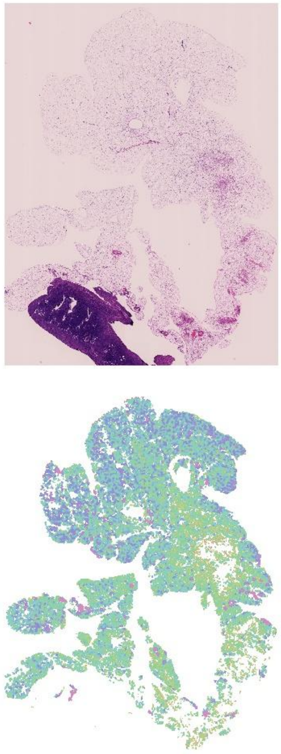

37.2c

(d) Female visceral gonadal MAT WT

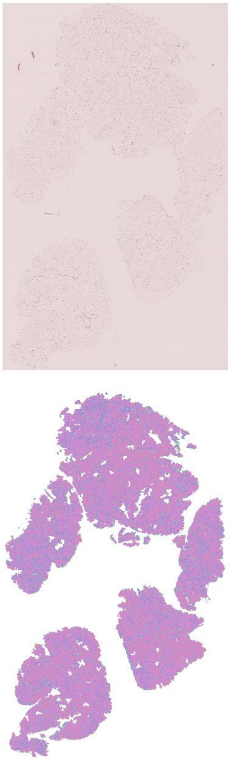

16.2a

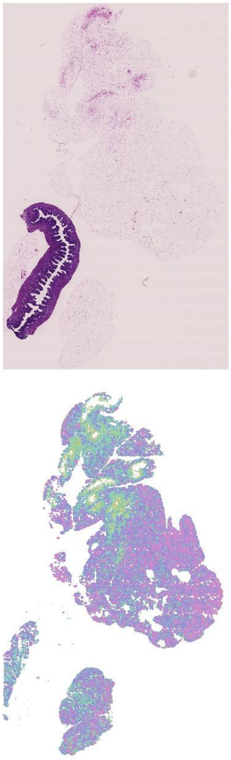

17.1d

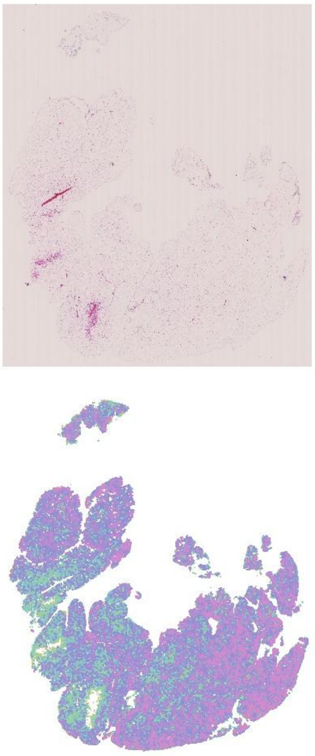

17.2b

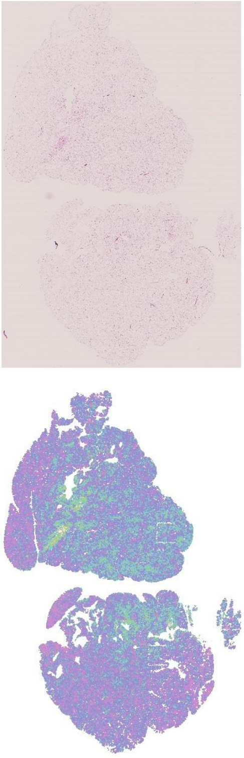

17.2d

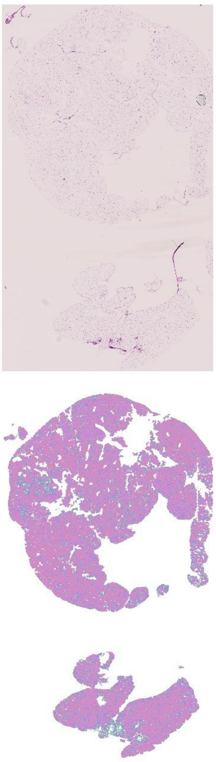

18.1c

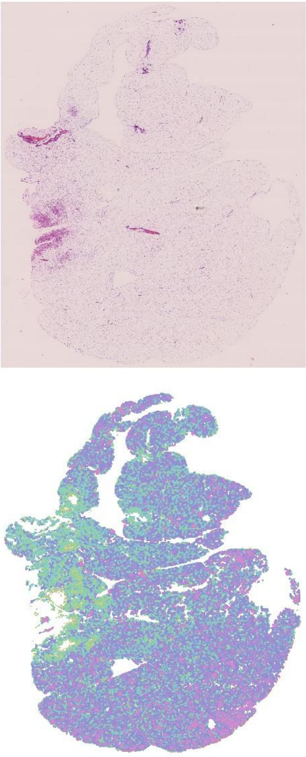

19.2b

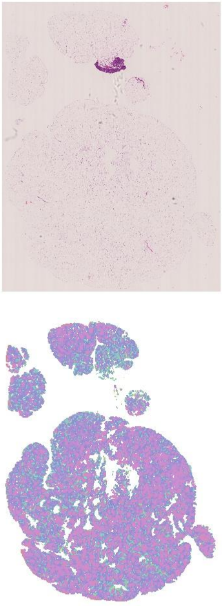

17.2a

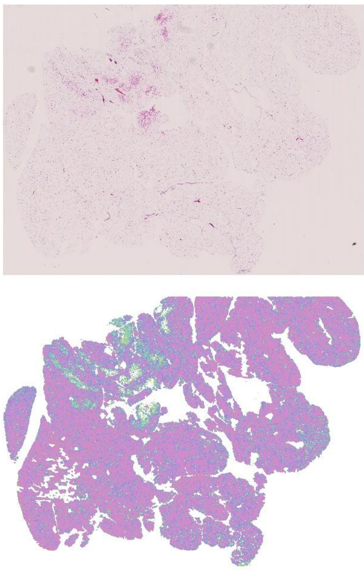

18.2c

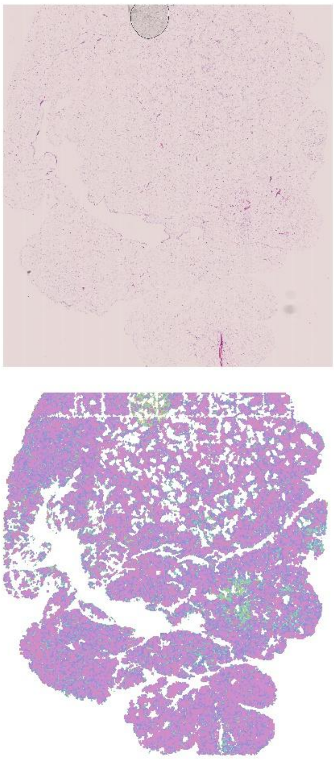

19.2c

<table>
<tr><td>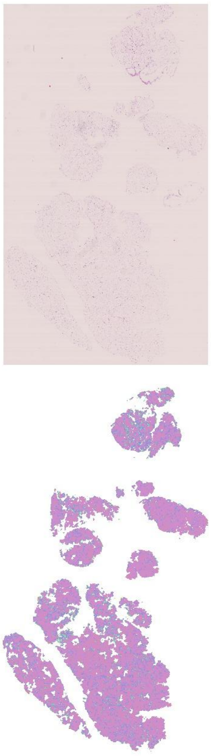<br>19.2e</td><td></td><td></td></tr>
<tr><td colspan="3">(e) Female visceral gonadal MAT Het</td></tr>
<tr><td>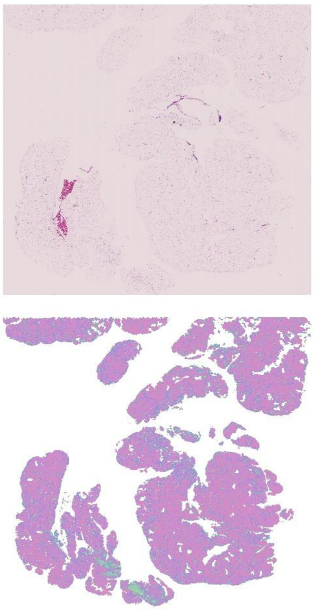<br>17.1b</td><td>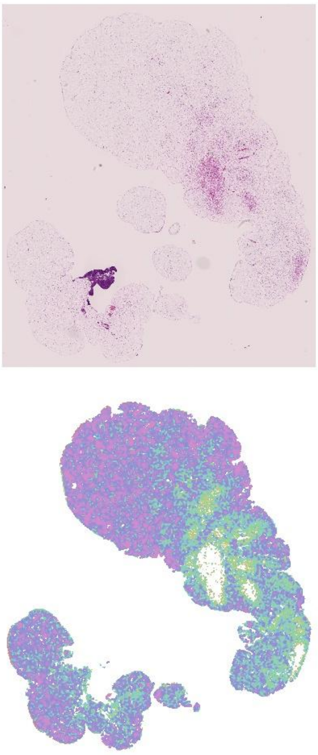<br>17.1c</td><td>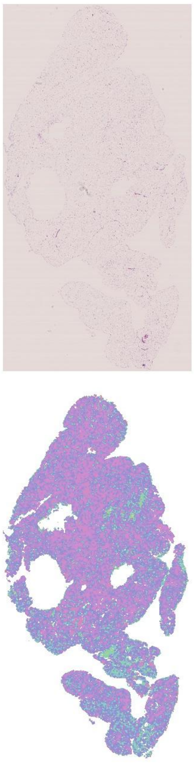<br>17.2c</td></tr>
</table>

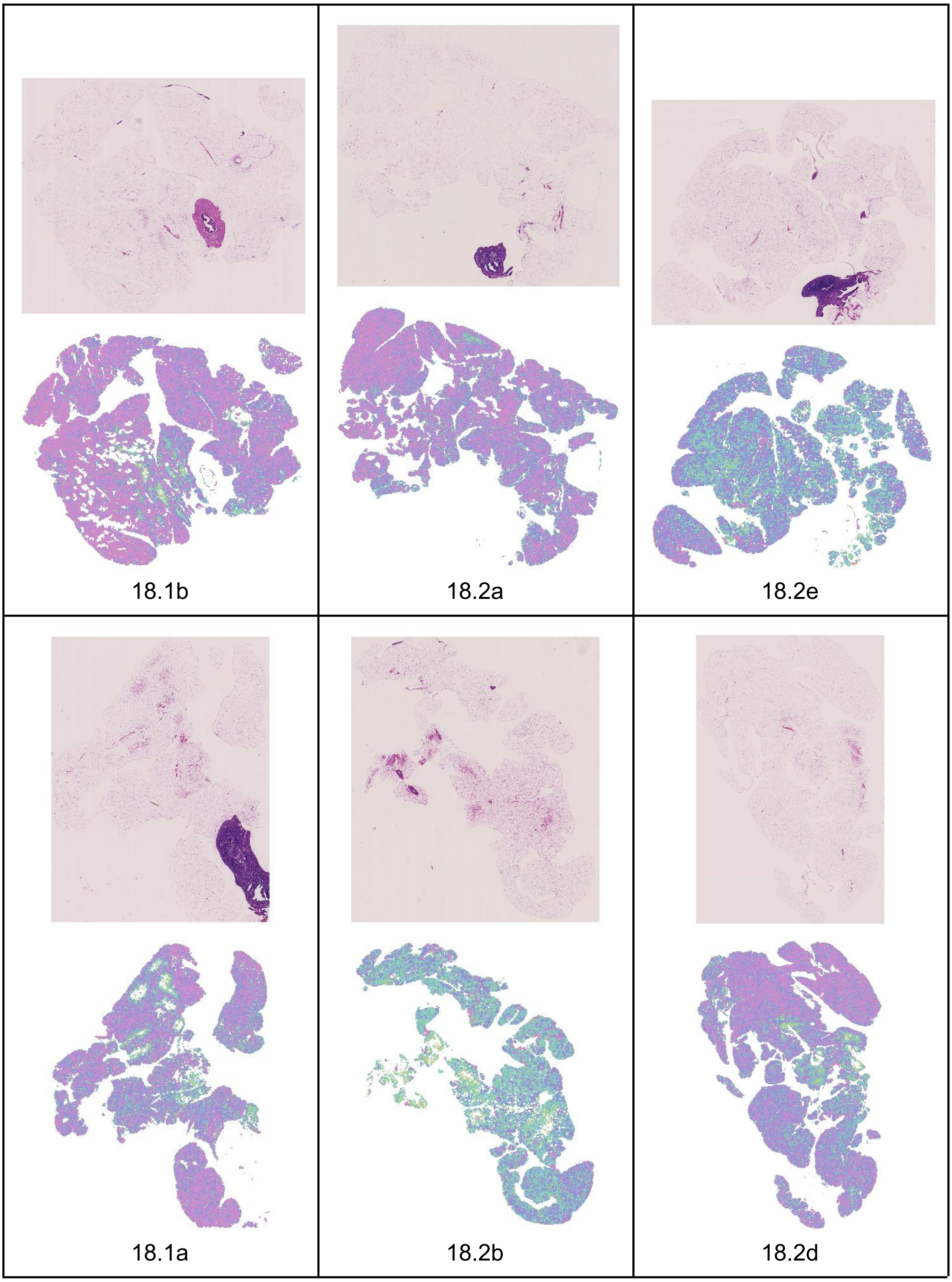

Fig. S1. White adipocyte tissue histology and area quantile heatmaps for female visceral gonadal depot. (a): Quantile colour map and cell area density for female Corrected segmentation with deciles plotted for reference (vertical black lines). (b): PAT WT. (c): PAT Het. (d): MAT WT. (e): MAT Het.

(a) Female inguinal subcutaneous PAT WT

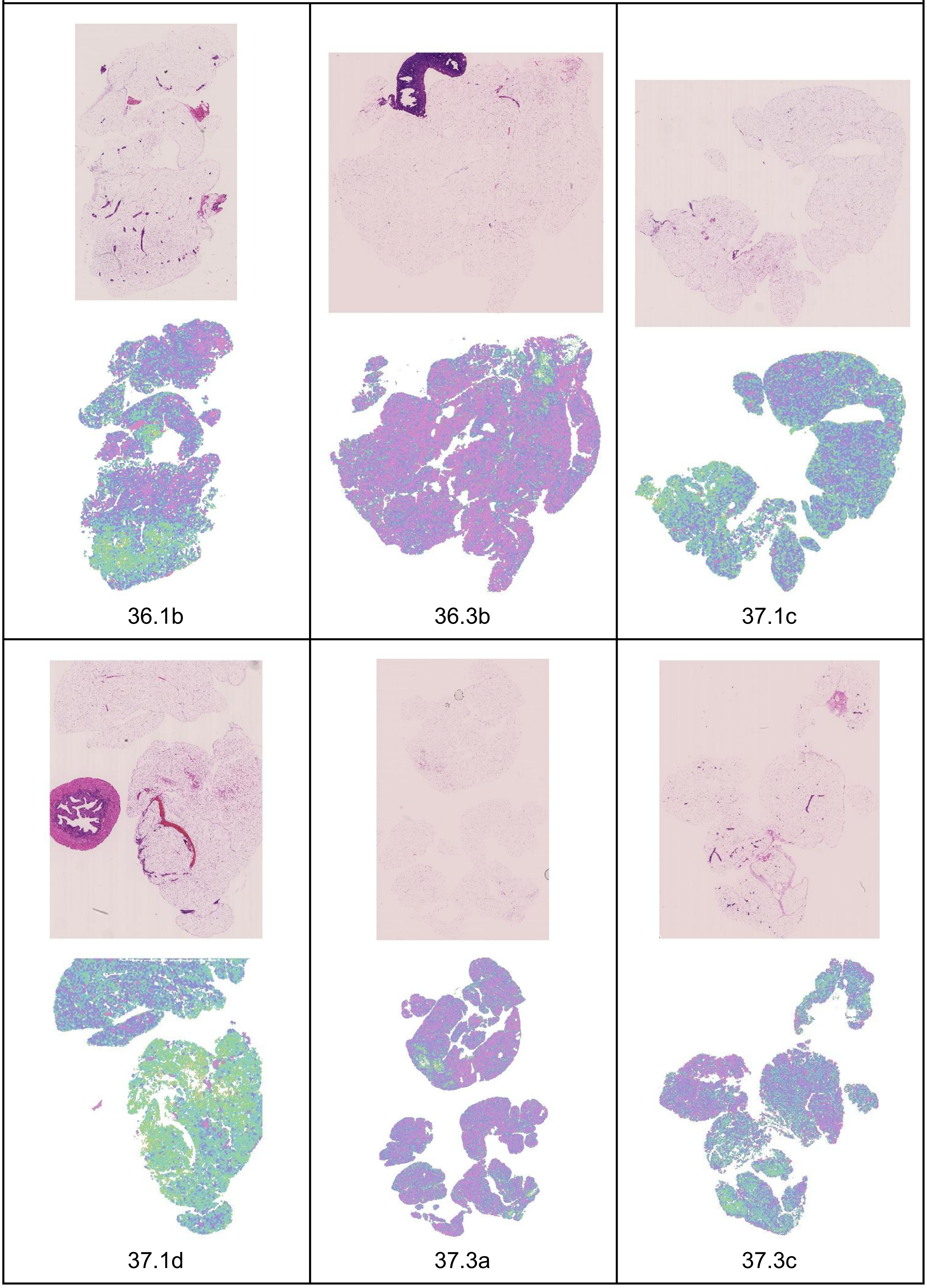

<table>
<tr><td>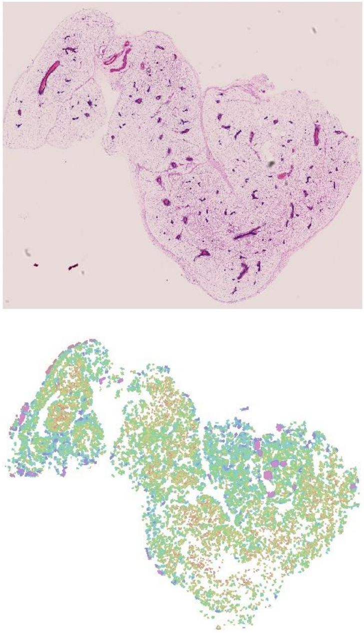<br>37.2b</td><td></td><td></td></tr>
<tr><td colspan="3">Female inguinal subcutaneous PAT Het</td></tr>
<tr><td>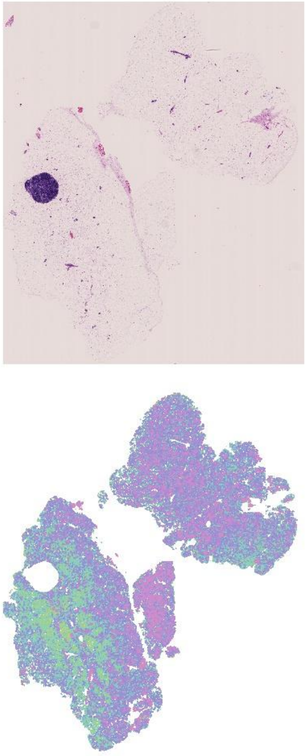<br>36.1a</td><td>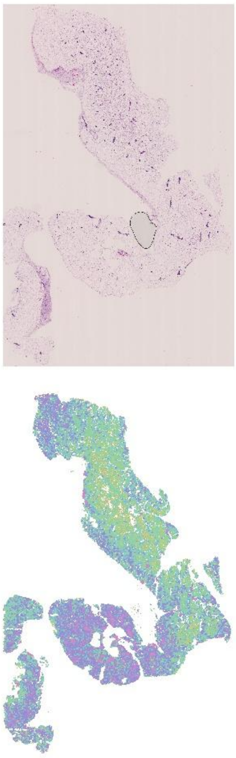<br>36.1c</td><td>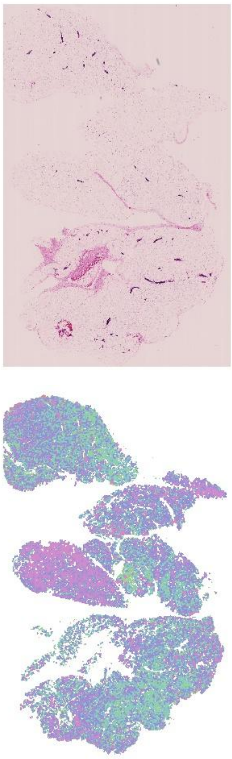<br>36.3a</td></tr>
</table>

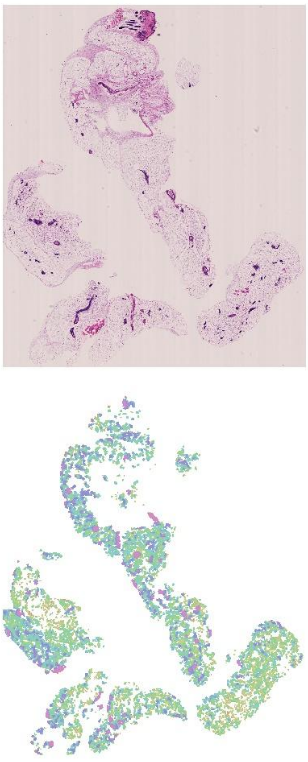

37.1a

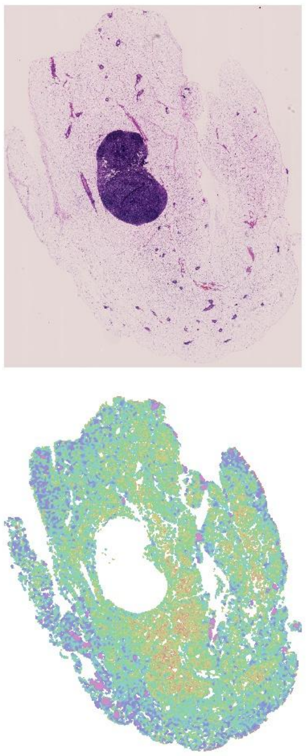

37.1e

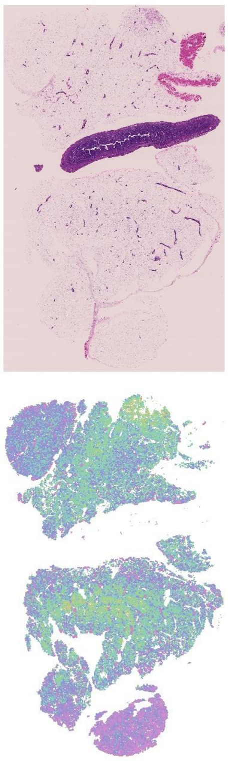

37.2e

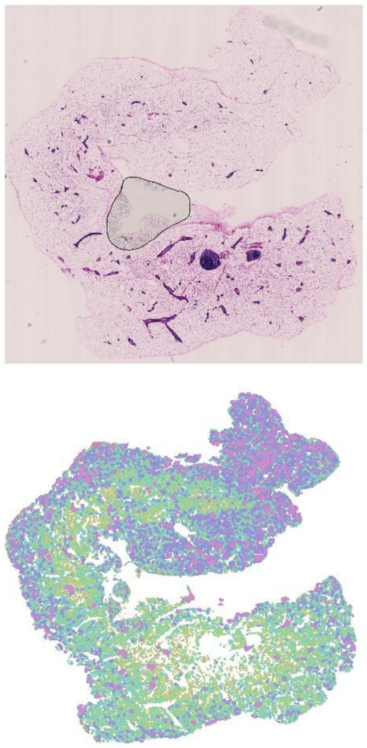

37.1b

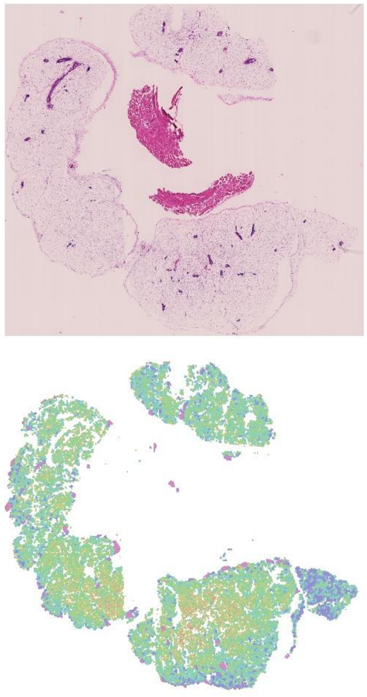

37.2a

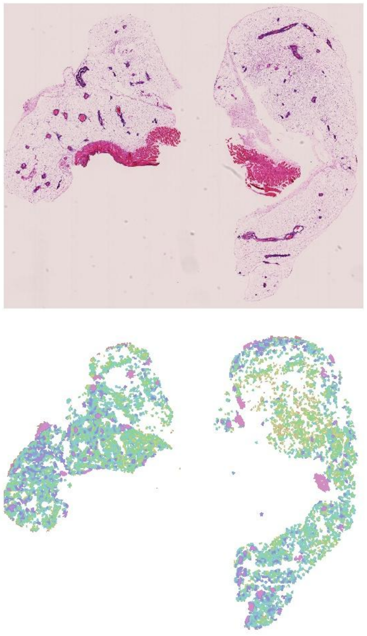

37.2c

<table>
<tr><td>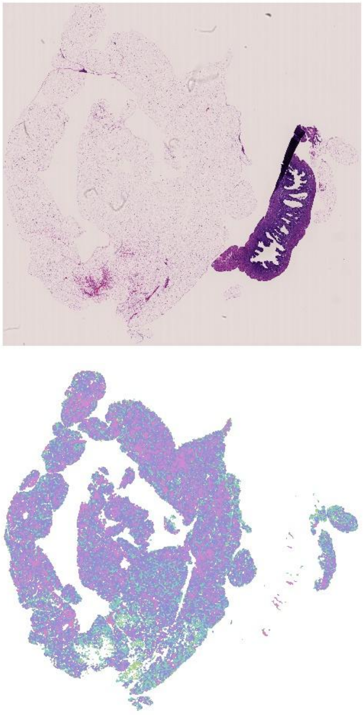<br>38.1a</td><td></td><td></td></tr>
<tr><td colspan="3">Female inguinal subcutaneous MAT WT</td></tr>
<tr><td>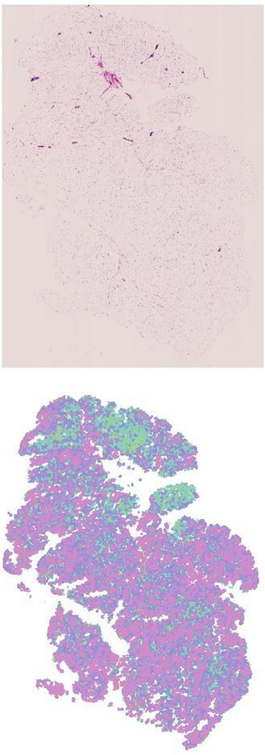<br>16.2a</td><td>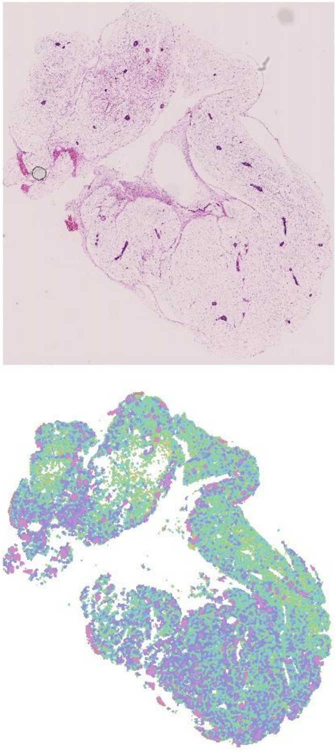<br>17.1d</td><td>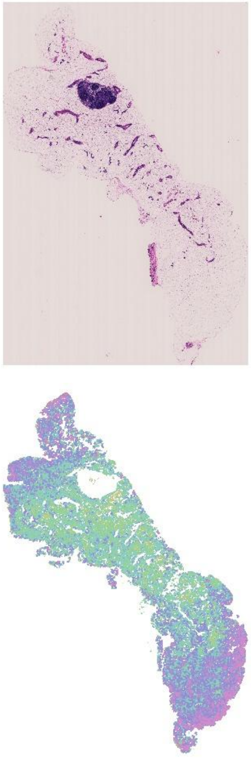<br>17.2b</td></tr>
</table>

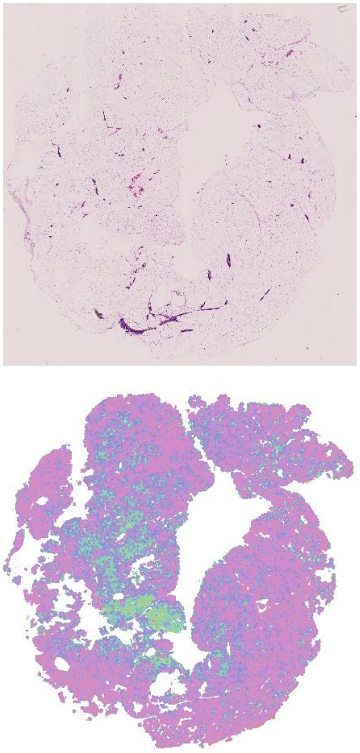

18.1c

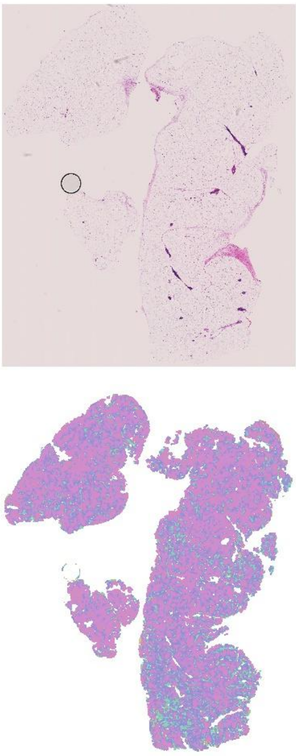

18.2c

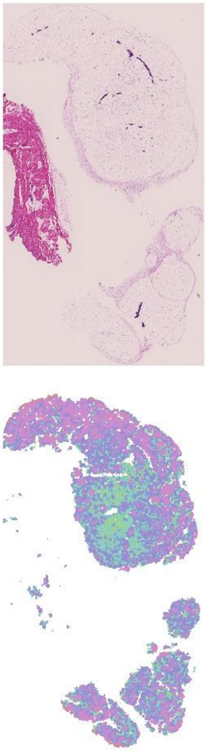

19.2c

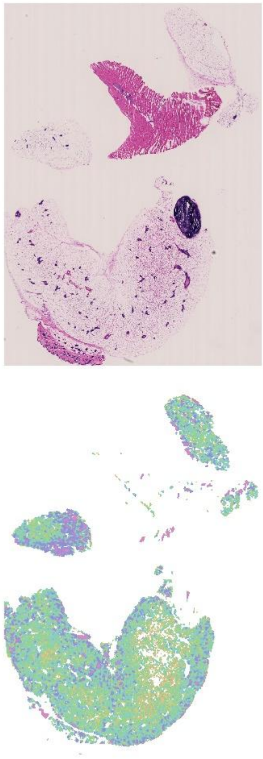

17.2a

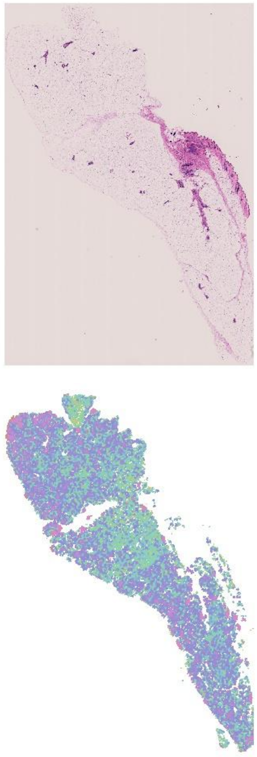

17.2d

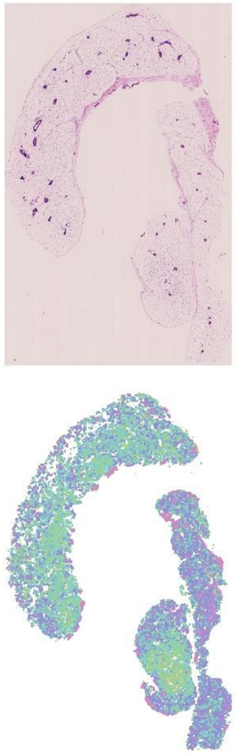

19.2b

| | | |
|---|---|---|
| 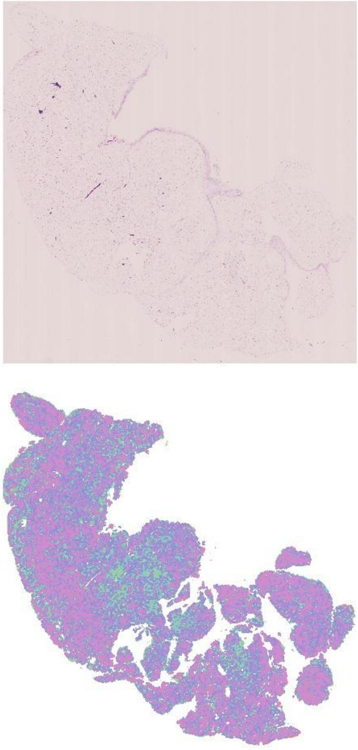<br>19.2e | | |
| Female inguinal subcutaneous MAT Het | | |
| 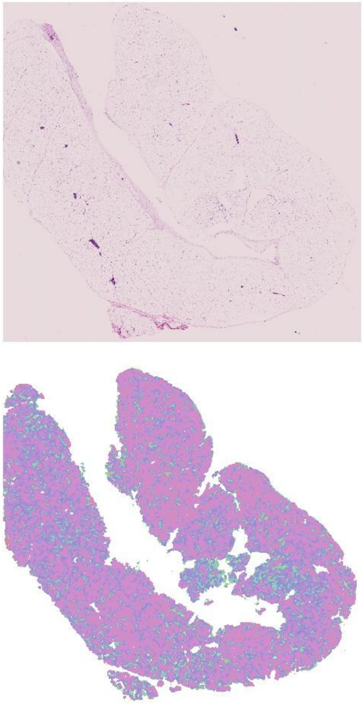<br>17.1b | 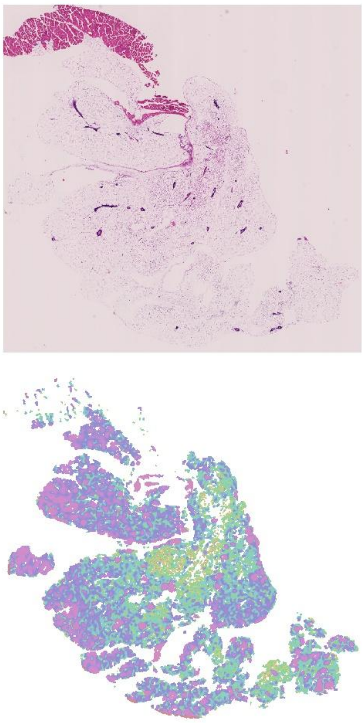<br>17.1c | 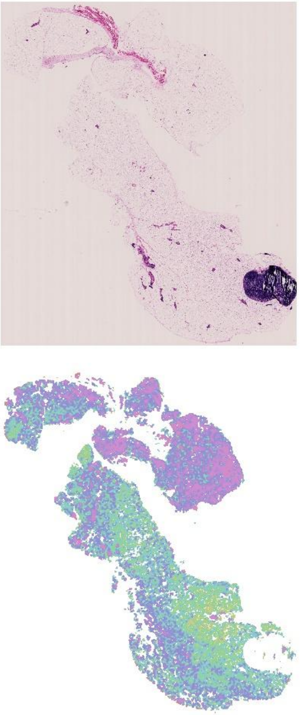<br>17.2c |

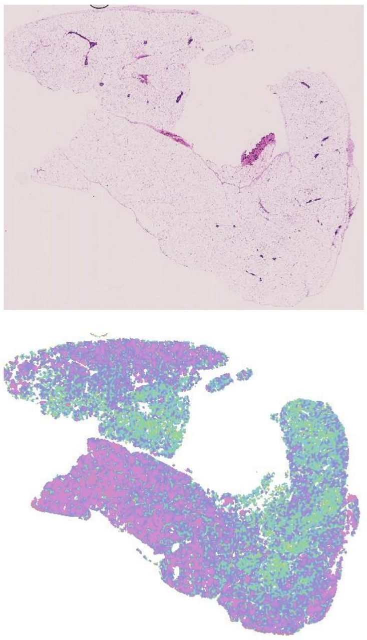

18.1b

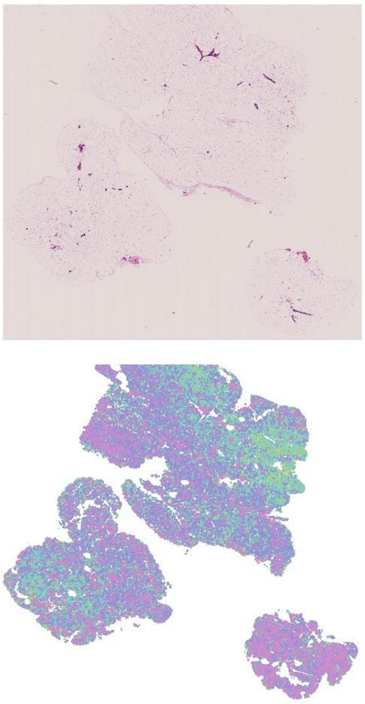

18.2a

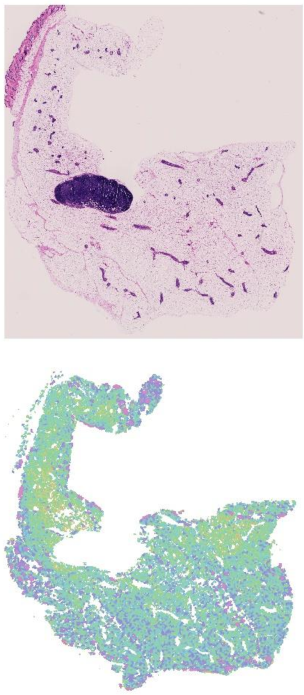

18.2e

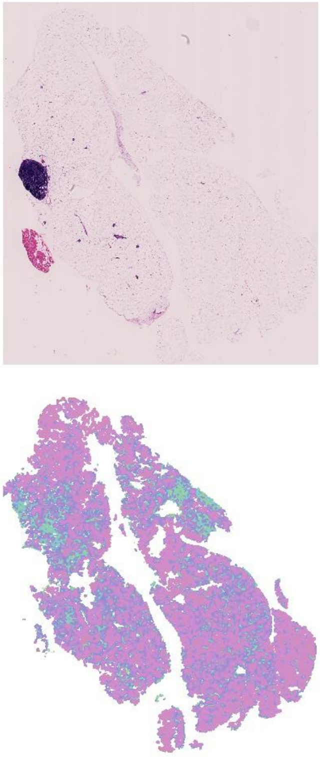

17.1a

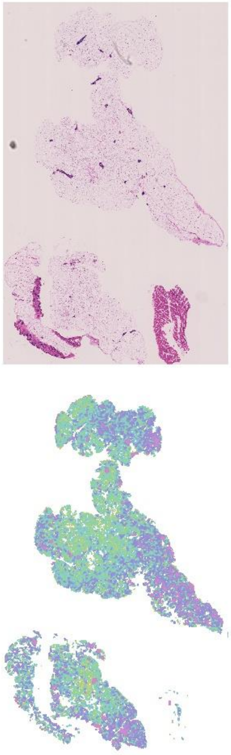

18.1a

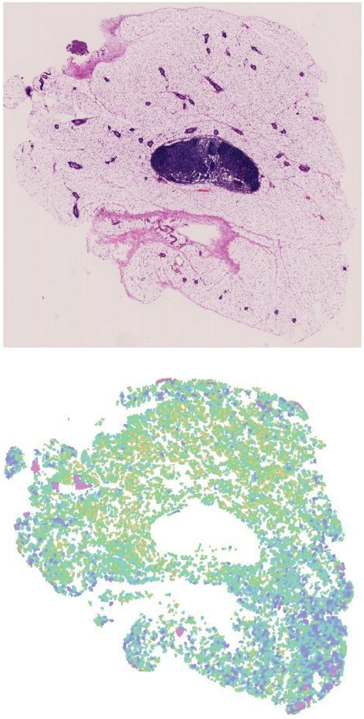

18.2b

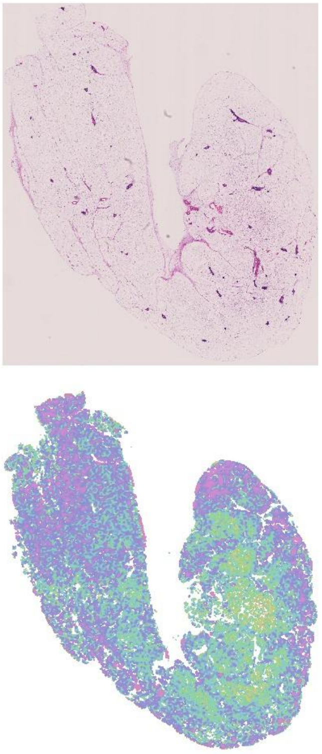

18.2d

Fig. S2. White adipocyte tissue histology and area quantile heatmaps for female inguinal subcutaneous depot. (a): PAT WT. (b): PAT Het. (c): MAT WT. (d): MAT Het.

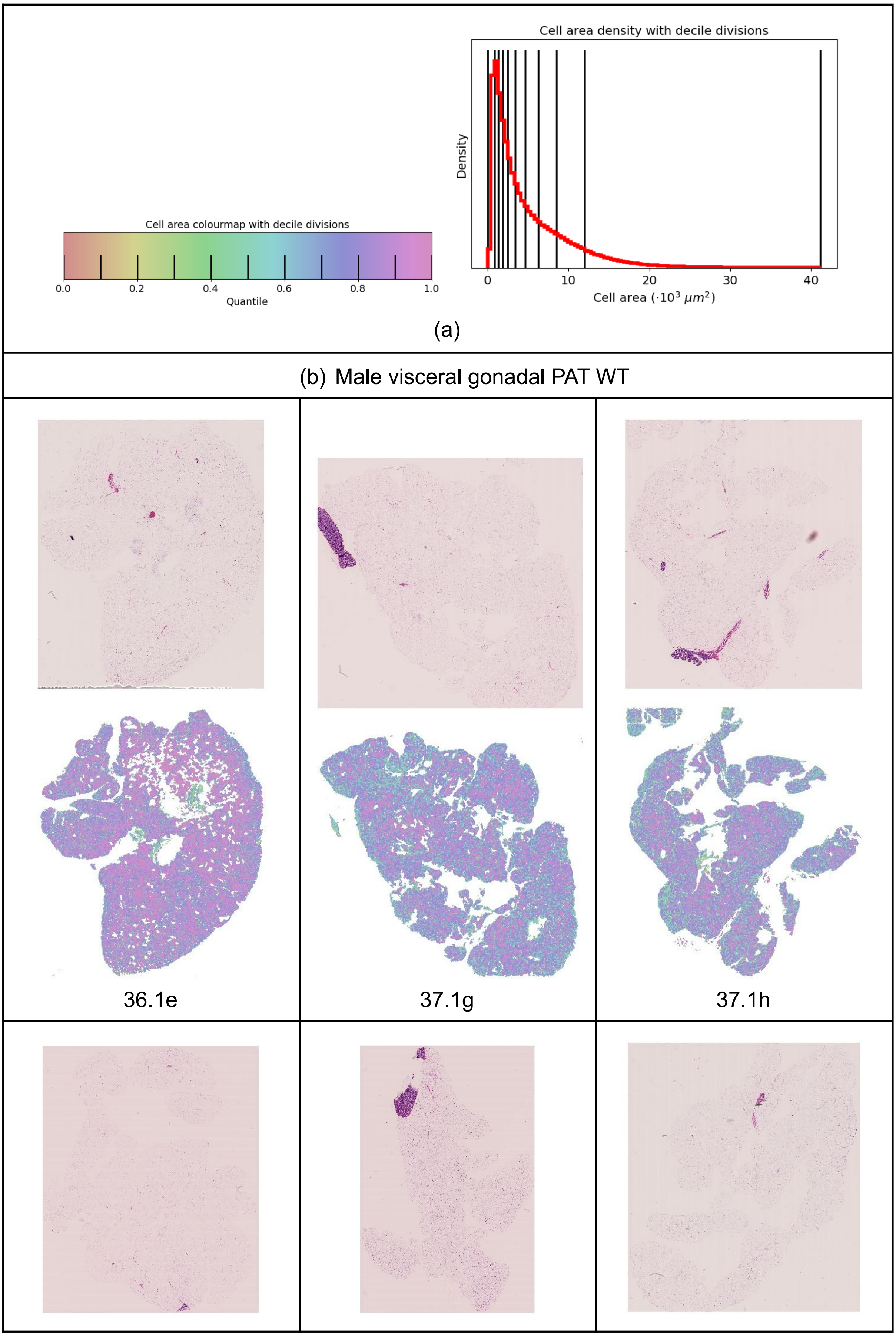
Cell area colourmap with decile divisions
0.0
0.2
0.4
0.6
0.8
1.0
Quantile
Cell area density with decile divisions
Density
0
10
20
30
40
Cell area (·10$^3$ $\mu m^2$)
(a)
(b) Male visceral gonadal PAT WT
36.1e
37.1g
37.1h

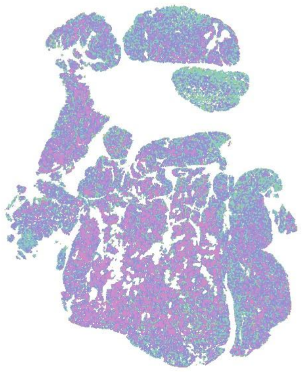

37.2h

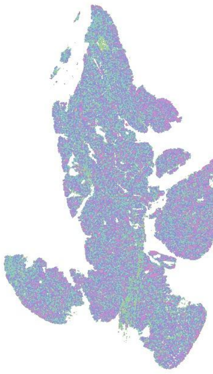

37.4b

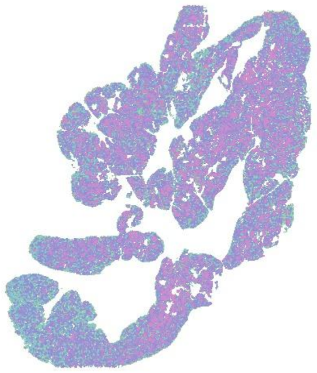

38.1f

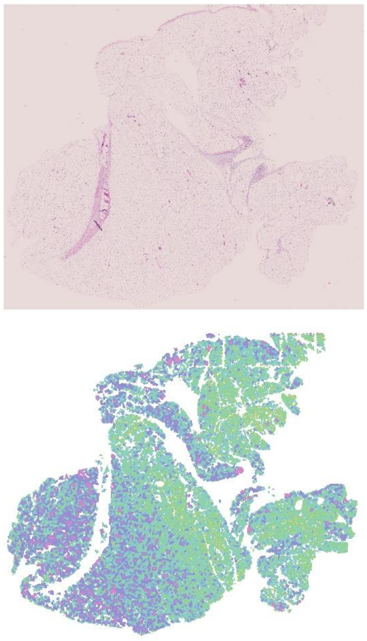

37.1f

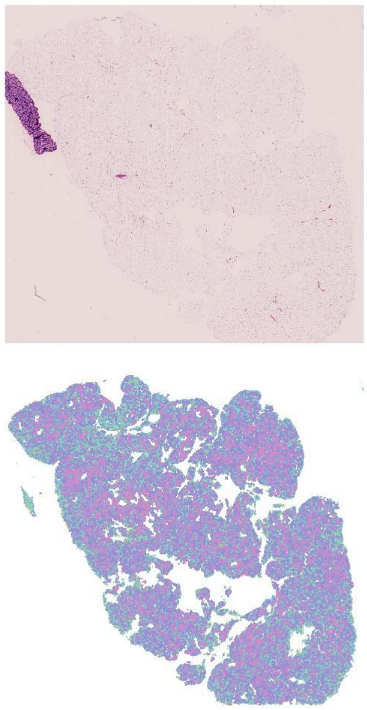

37.1g

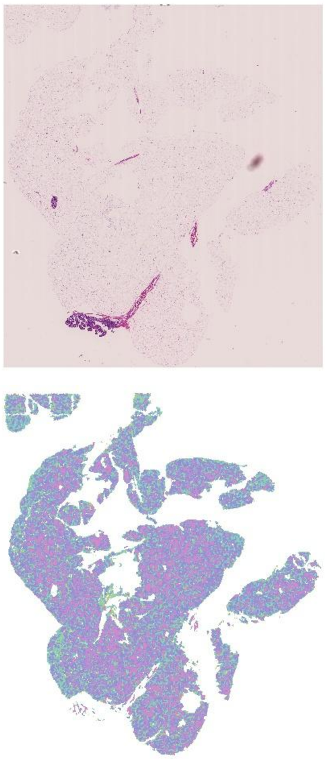

37.1h

(c) Male visceral gonadal PAT Het

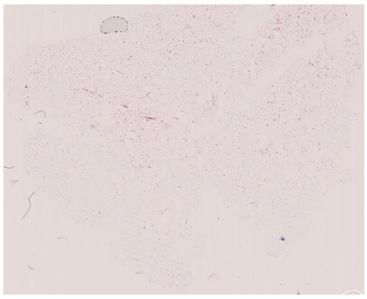

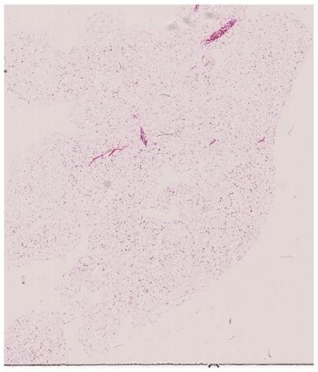

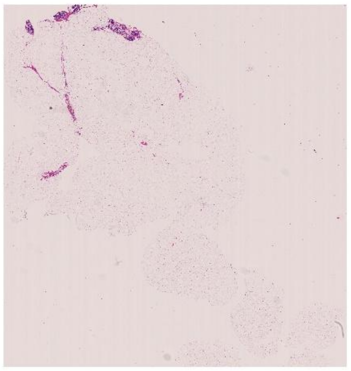

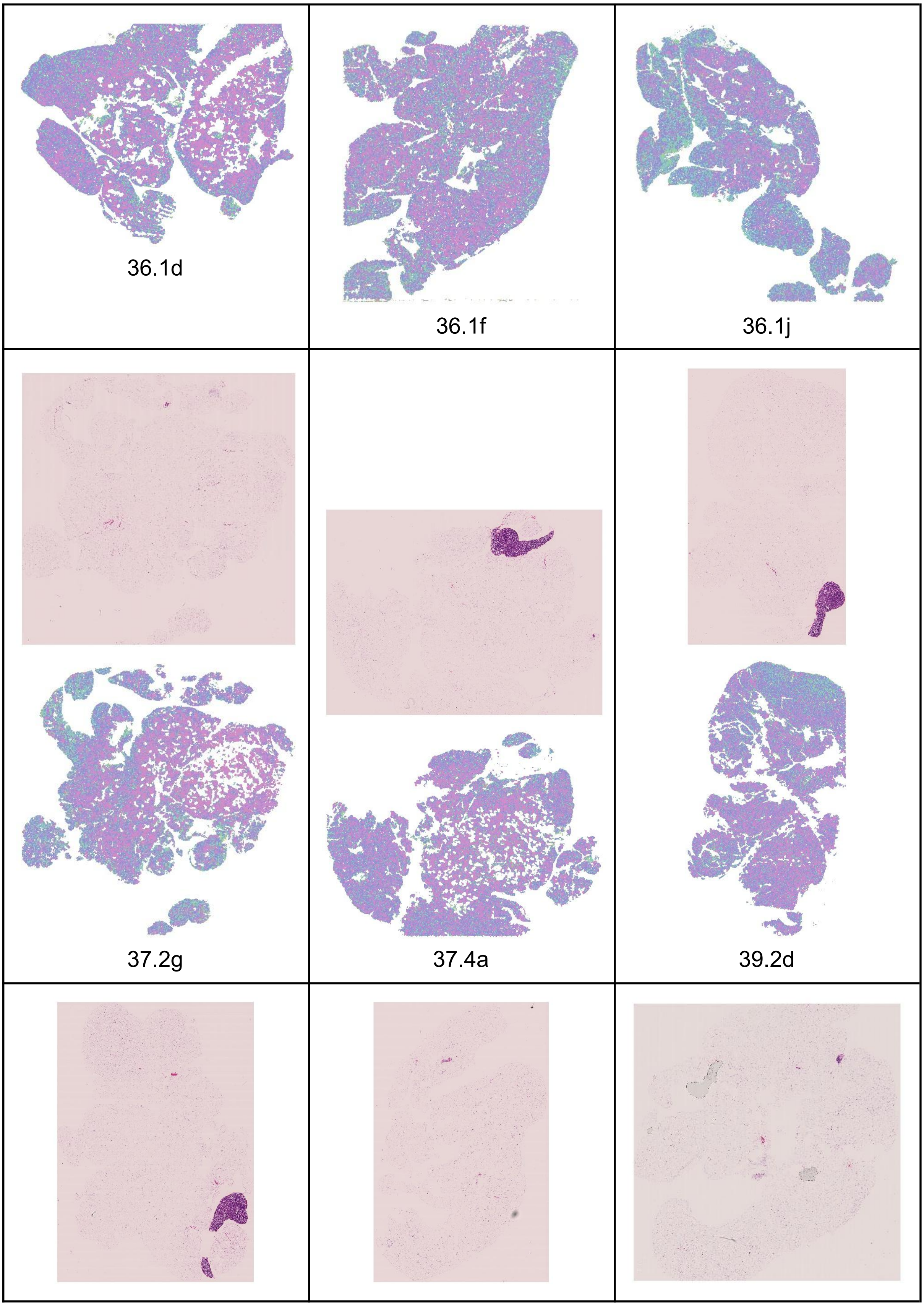
36.1d
36.1f
36.1j
37.2g
37.4a
39.2d

| | | |
|---|---|---|
| 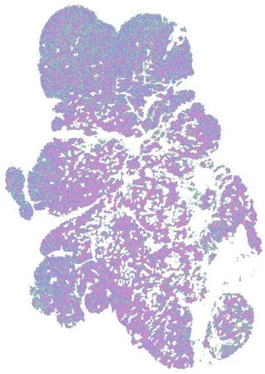 | 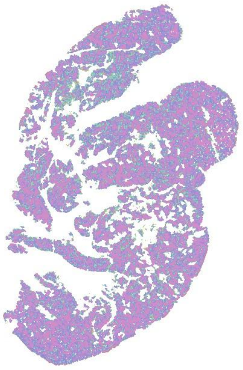 | 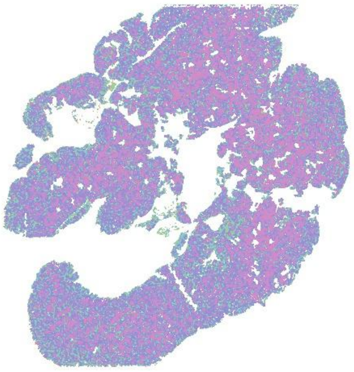 |
| 36.3d | 37.2f | 38.1e |

(d) Male visceral gonadal MAT WT

| | | |
|---|---|---|
| 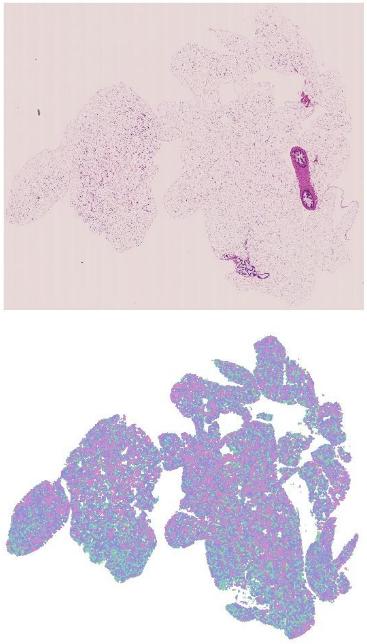 | 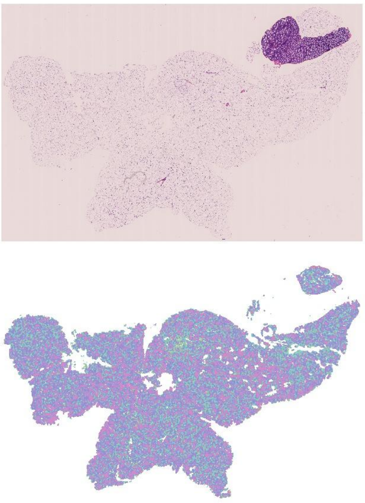 | 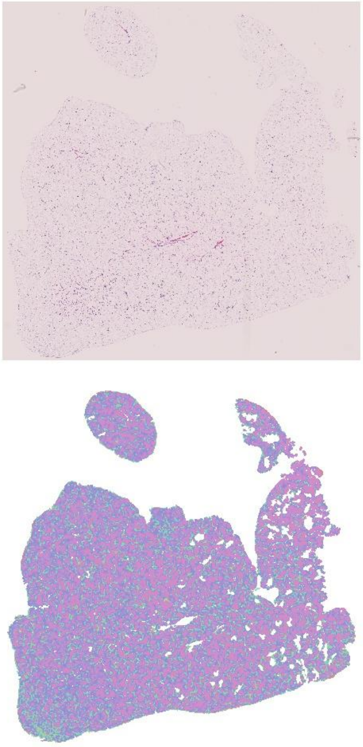 |
| 16.2b | 16.2e | 17.1e |
| 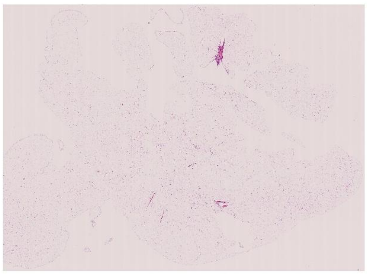 | 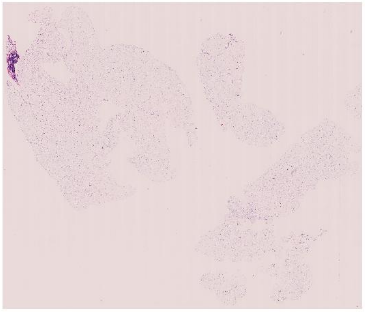 | 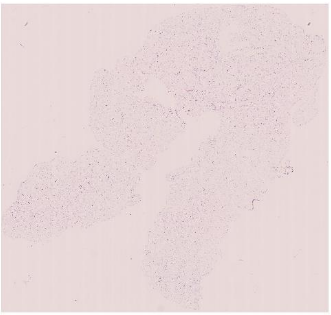 |

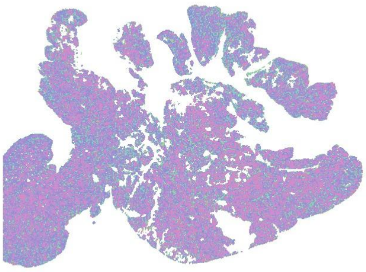

18.1d

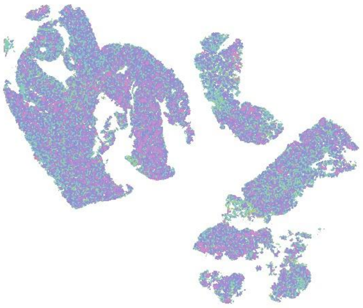

18.3c

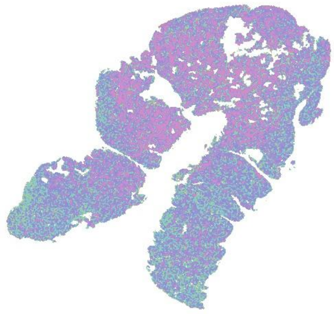

19.2f

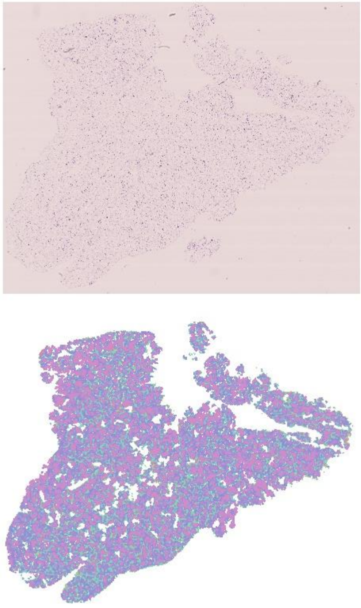

16.2f

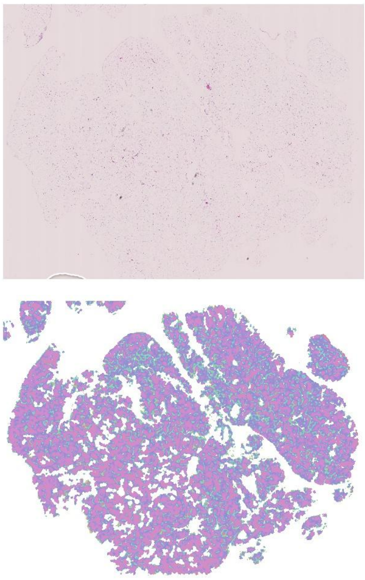

17.1f

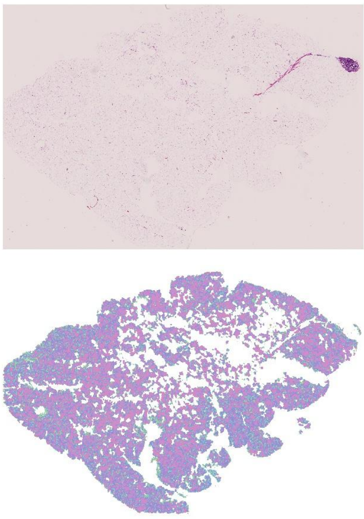

19.1a

(e) Male visceral gonadal MAT Het

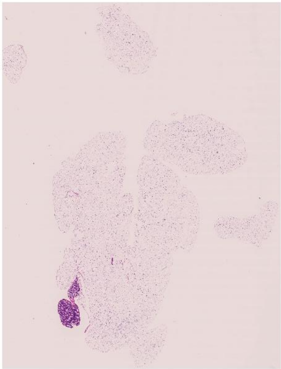

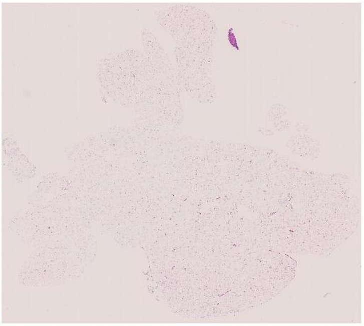

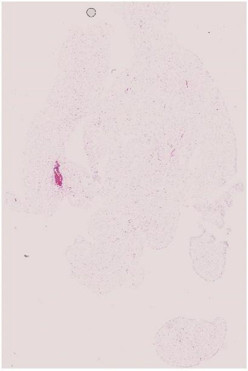

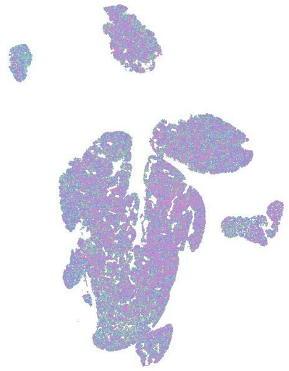

16.2c

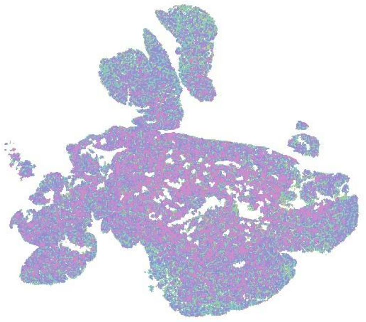

17.2f

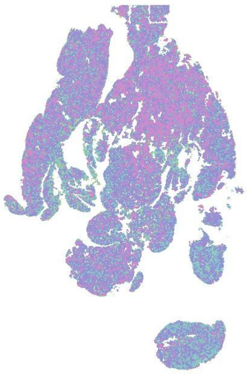

18.2f

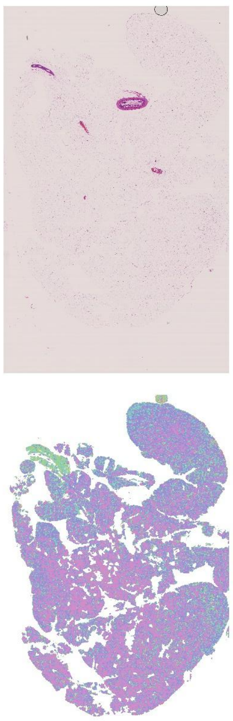

18.2g

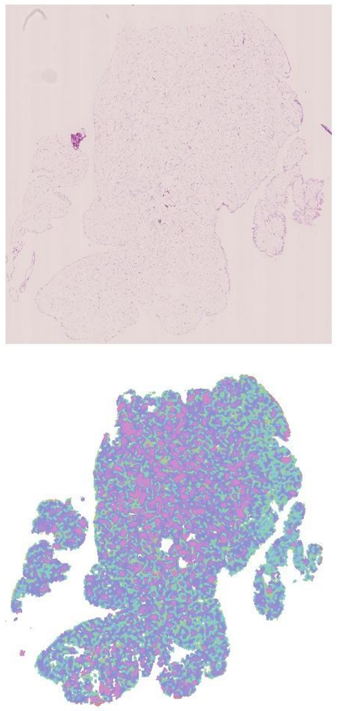

18.3b

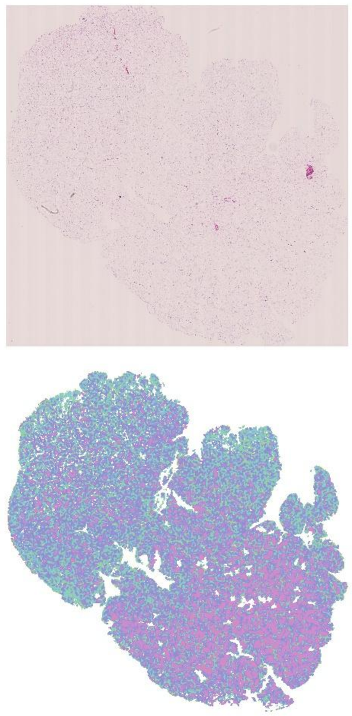

18.3d

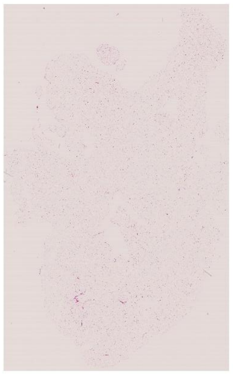

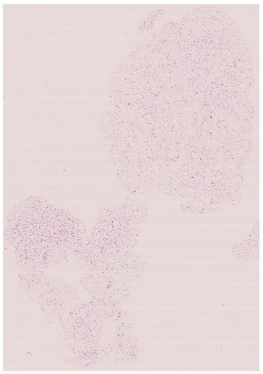

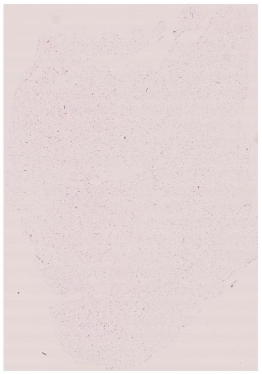

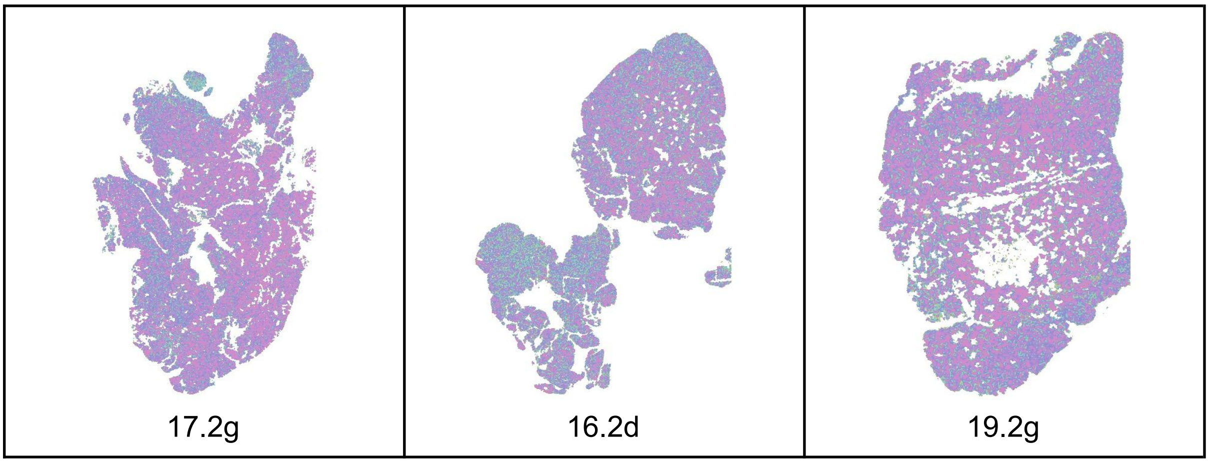


Fig. S3. White adipocyte tissue histology and area quantile heatmaps for male visceral gonadal depot. (a): Quantile colour map and cell area density for male Corrected segmentation with deciles plotted for reference (vertical black lines). (b): PAT WT. (c): PAT Het. (d): MAT WT. (e): MAT Het.

| Male inguinal subcutaneous PAT WT | | |
|---|---|---|
| 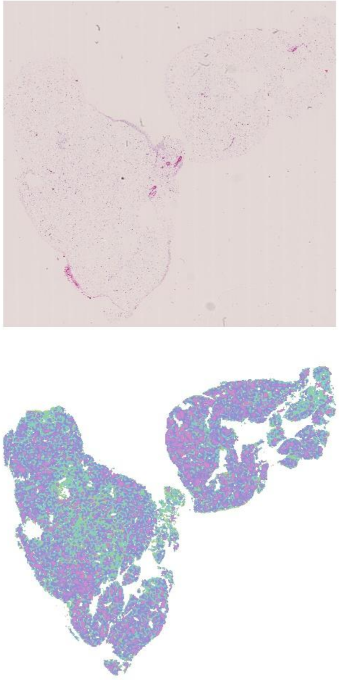 36.1e | 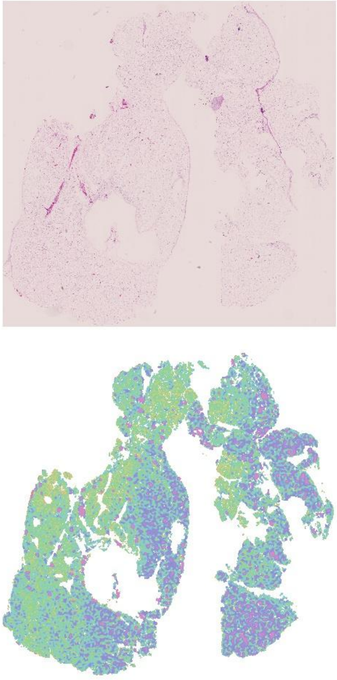 37.1g | 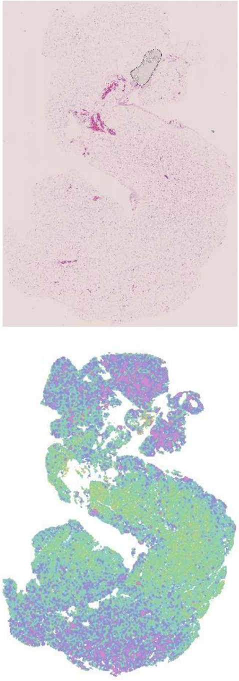 37.1h |
| 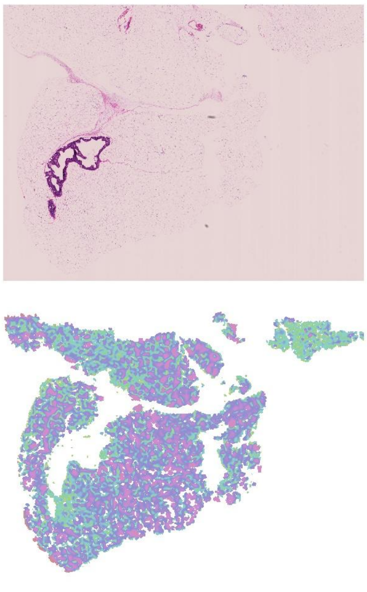 37.2h | 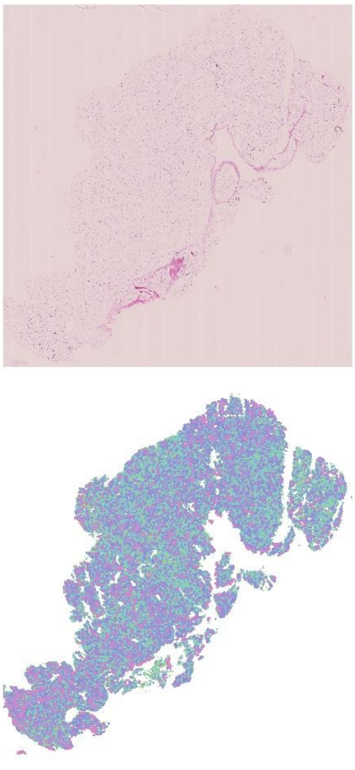 37.4b | 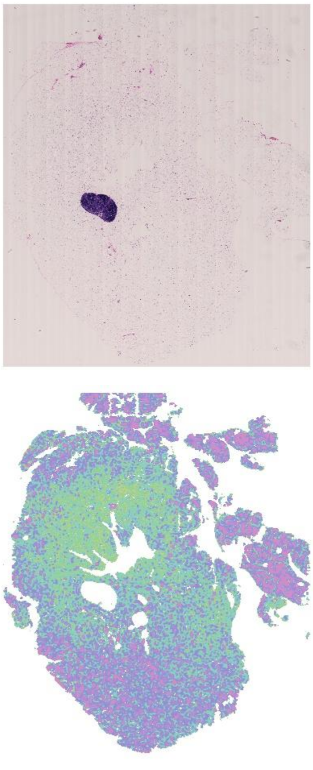 38.1f |

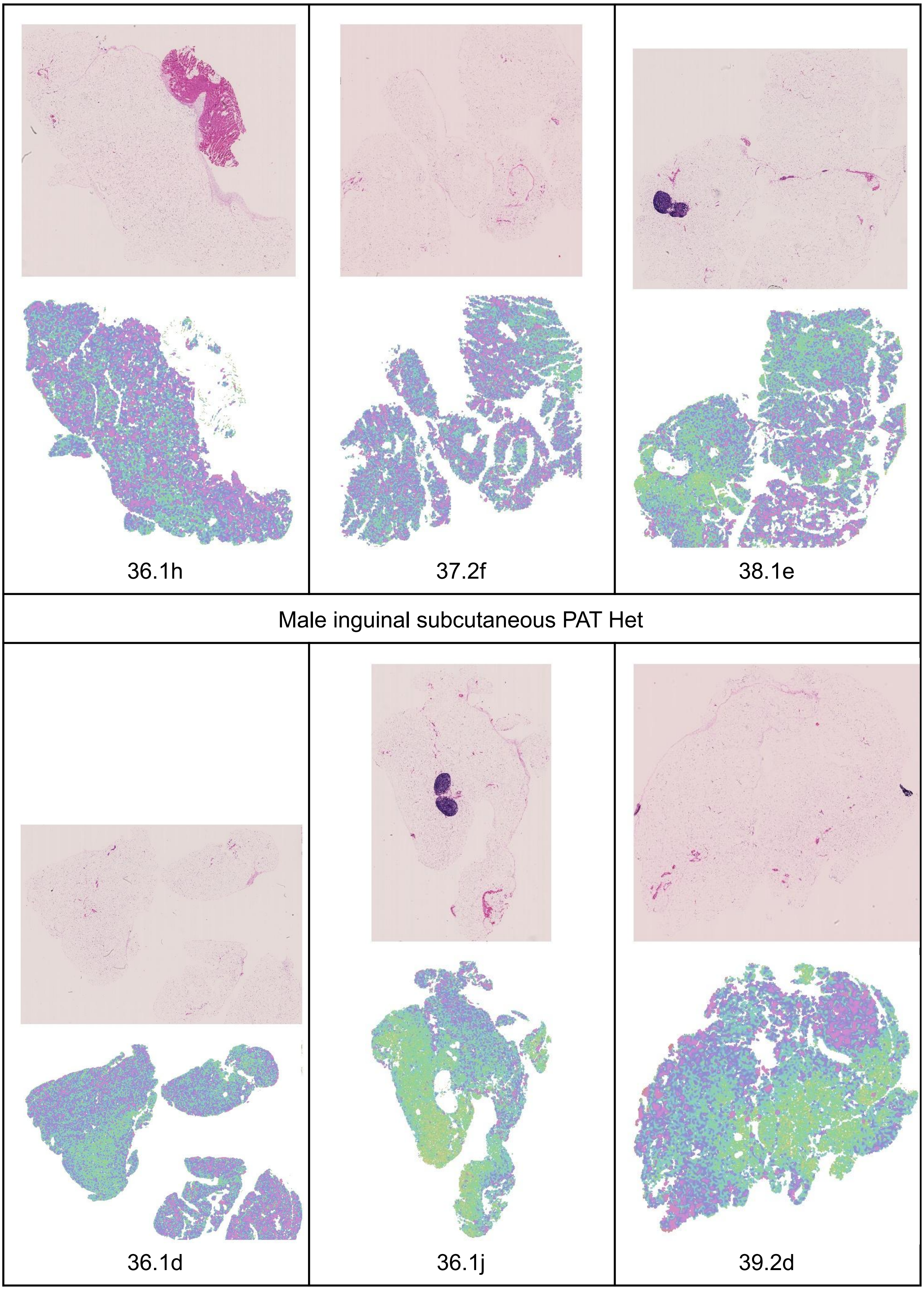
36.1h
37.2f
38.1e
Male inguinal subcutaneous PAT Het
36.1d
36.1j
39.2d

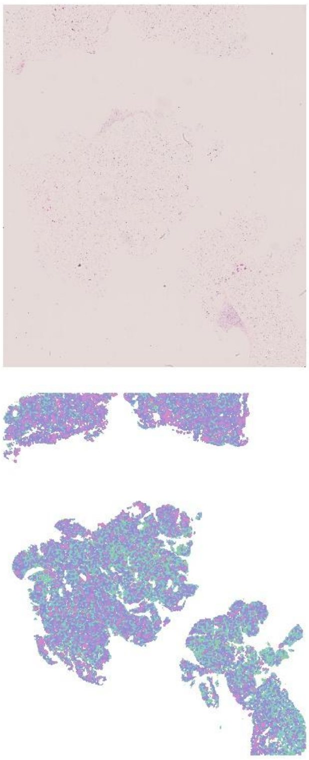

36.1f

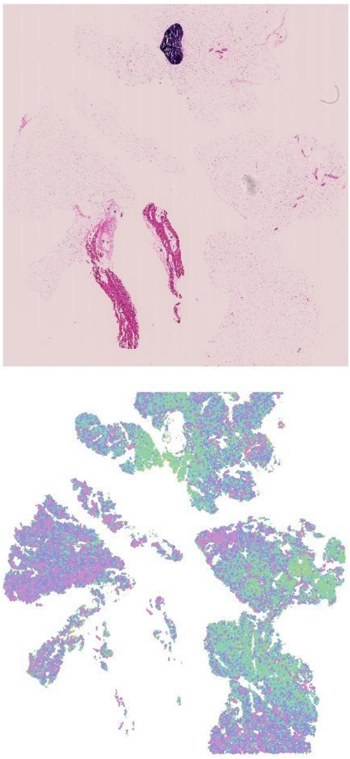

37.2g

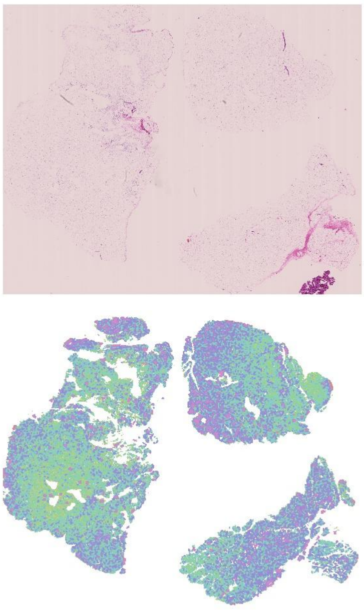

37.4a

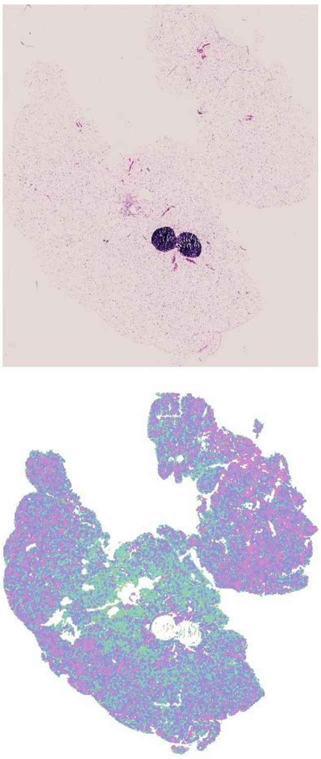

36.1g

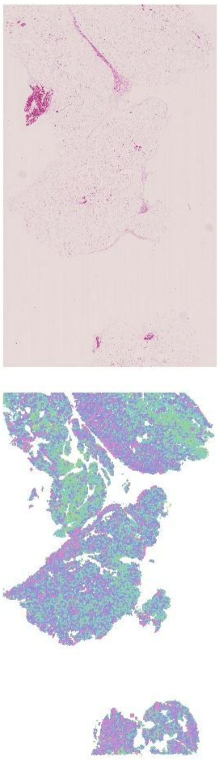

36.1i

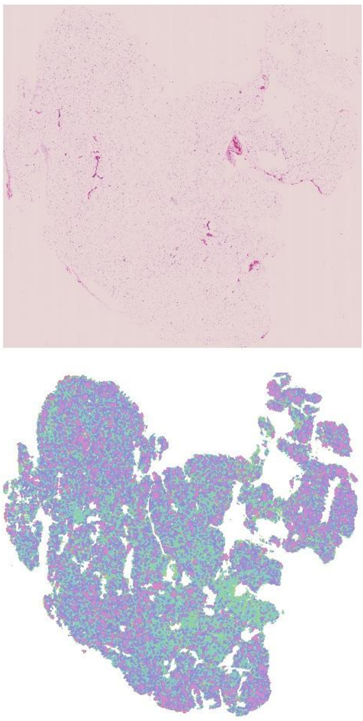

36.3d

Male inguinal subcutaneous MAT WT

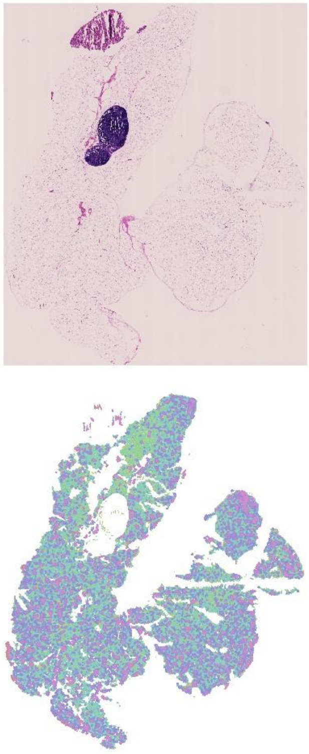

16.2b

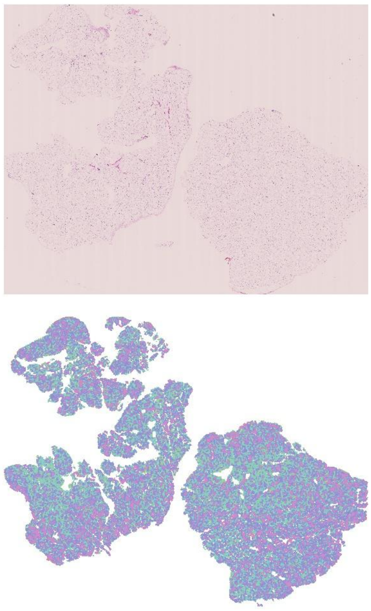

16.2e

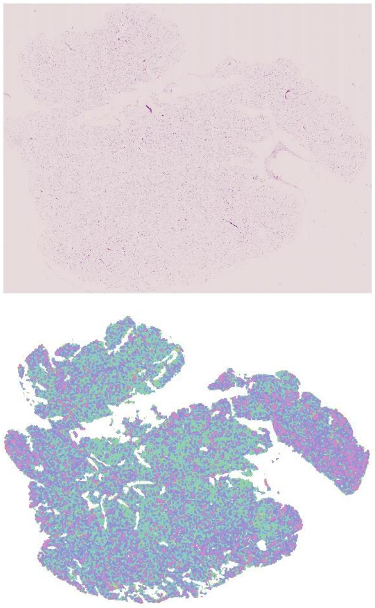

17.1e

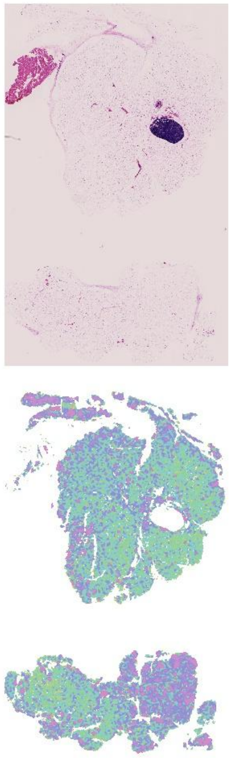

18.1d

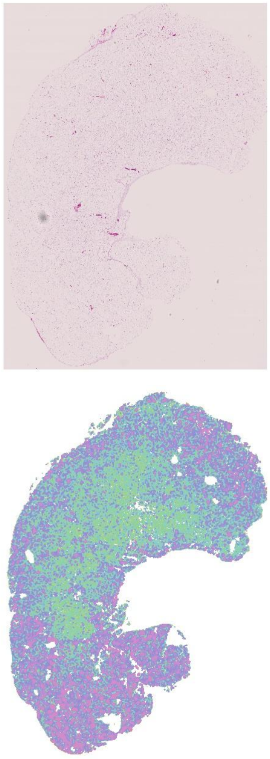

18.1f

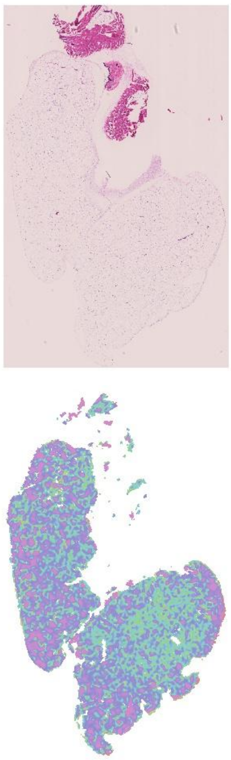

18.3c

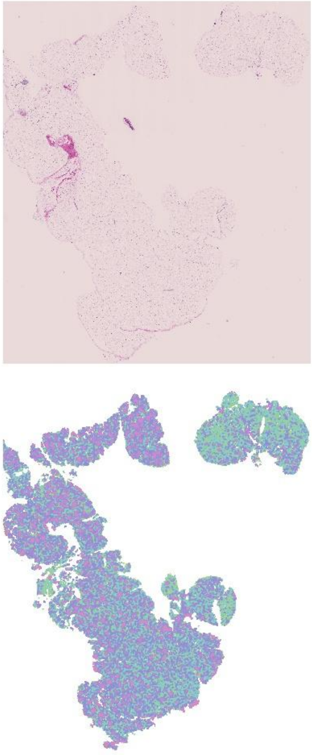

16.2f

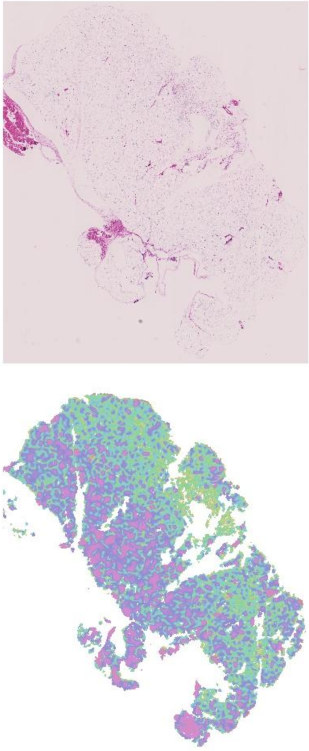

17.1f

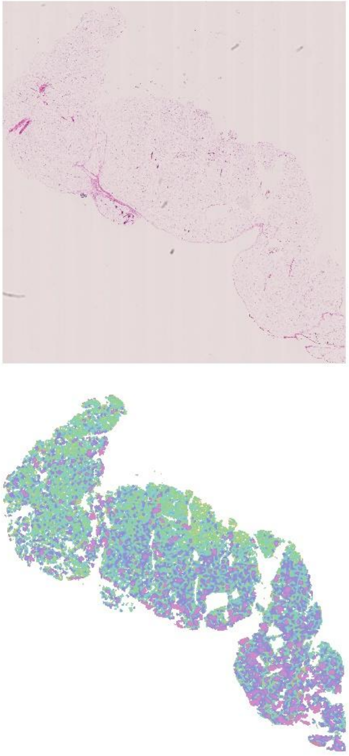

19.1a

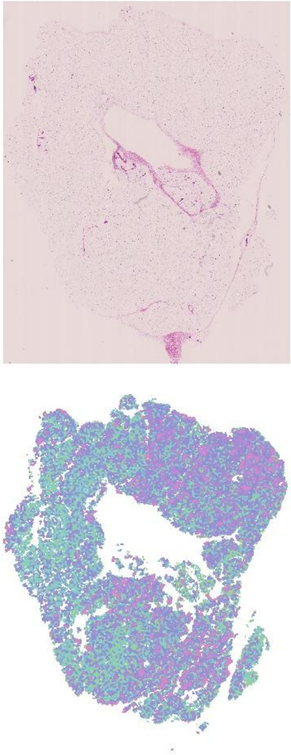

19.2f

Male inguinal subcutaneous MAT Het

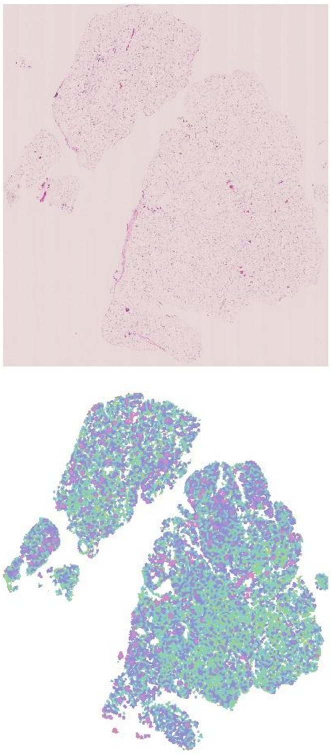

16.2c

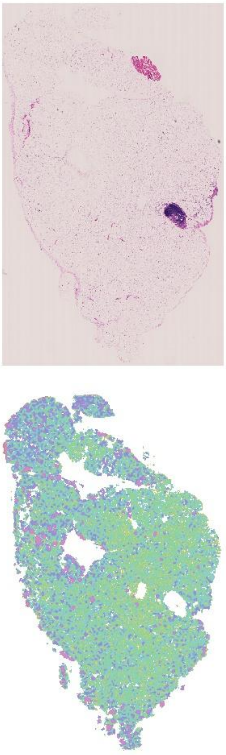

17.2f

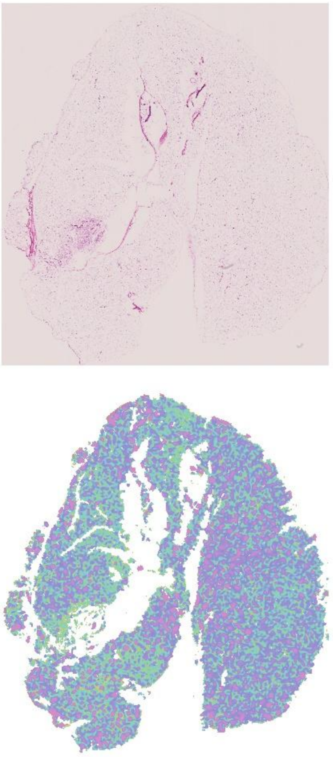

18.2f

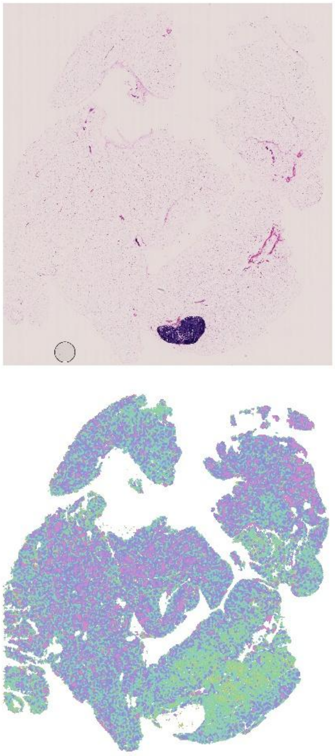

18.2g

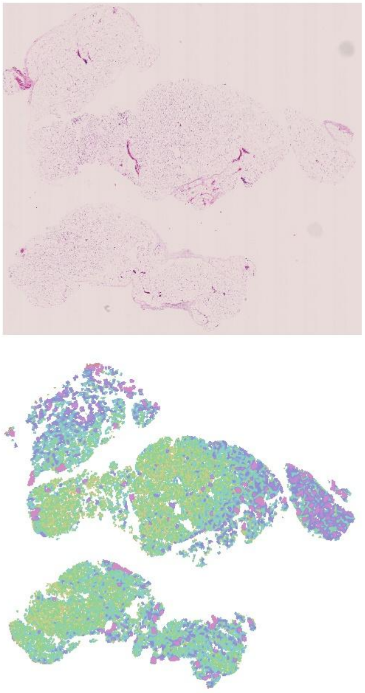

18.3b

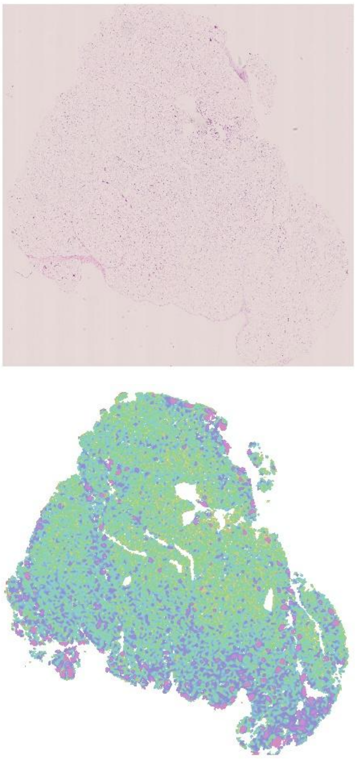

18.3d

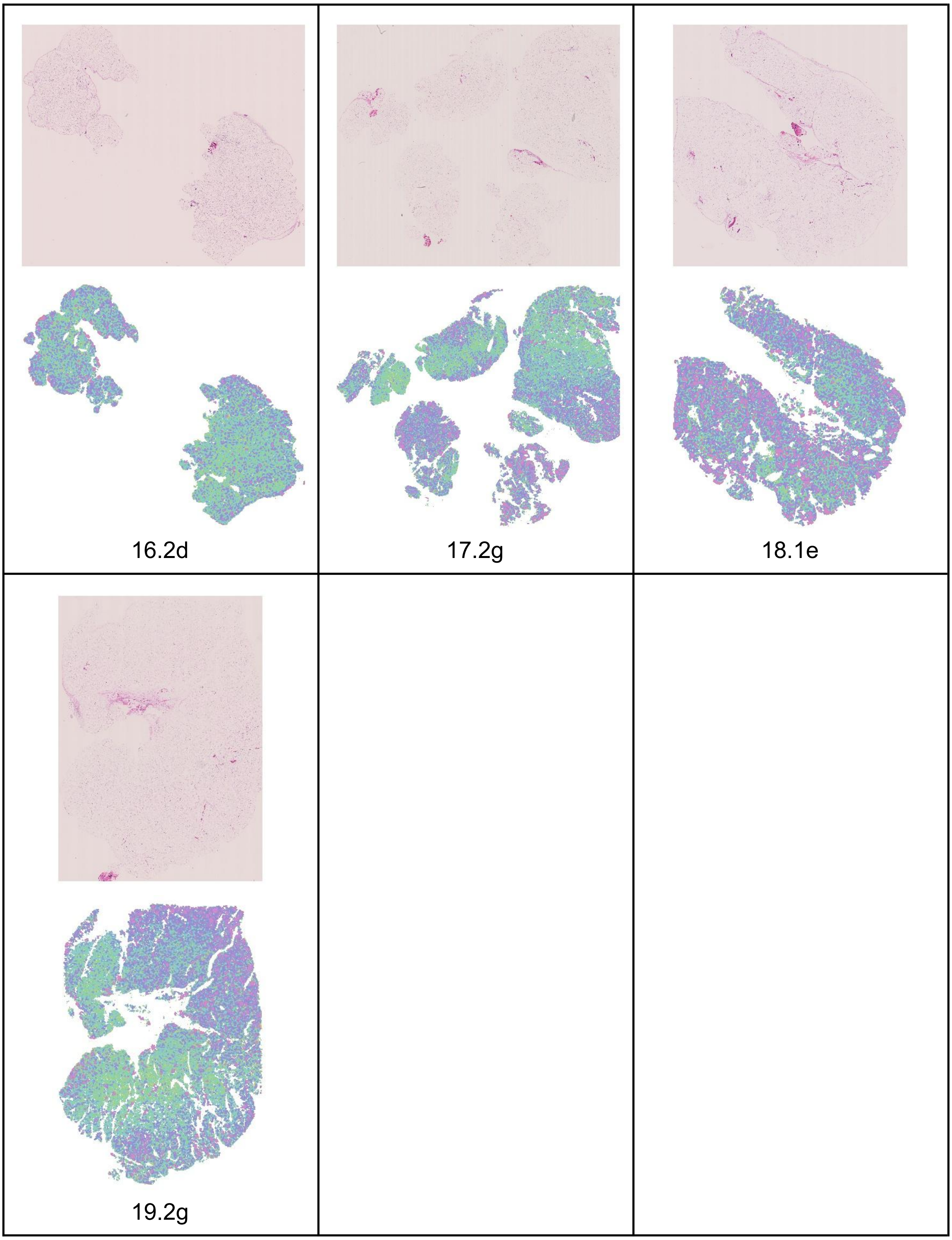


Fig. S4. White adipocyte tissue histology and area quantile heatmaps for male inguinal subcutaneous depot. (a): PAT WT. (b): PAT Het. (c): MAT WT. (d): MAT Het.
